# Ecological changes have driven biotic exchanges across the Indian Ocean

**DOI:** 10.1101/2021.11.18.469077

**Authors:** Samuel C. Bernardes, Kristina von Rintelen, Thomas von Rintelen, Almir R. Pepato, Timothy J. Page, Mark de Bruyn

**Affiliations:** Museum für Naturkunde, Leibniz Institute for Evolution and Biodiversity Science, Berlin, Germany; Laboratório de Acarologia, Instituto de Ciências Biológicas, Universidade Federal de Minas Gerais, Belo Horizonte, Brazil; Australian Rivers Institute, Griffith University, Queensland, Australia; School of Life and Environmental Sciences, University of Sydney, Sydney, Australia

**Keywords:** Vicariance, dispersal, transoceanic, Laurasia, Gondwana, India, Madagascar, Indo-Australian Archipelago

## Abstract

The Indian Ocean has a complex geological history that has drawn the attention of naturalists for almost a century now. Due to its tectonic history, many geological elements and processes have been evoked to explain the exchange of species between landmasses. Here, we revisited previous studies on twenty-three taxa to investigate trends across time since the Gondwana breakup. We investigated these datasets by applying a time-calibrated Bayesian framework to them and reconstructing their ancestral ranges. We conclude that ecological transformations have presented opportunities for the establishment of migrants. The role of donating and receiving migrants has shifted several times according to these transformations. Time-specific trends show weak evidence for the stepping-stones commonly suggested as physical routes between landmasses. However, before its collision with Asia, India may have served as an intermediary for such exchanges.

## Introduction

The Indian Ocean (hereafter IO) is the smallest, youngest and warmest of the three major oceans. IO’s boundaries include continents (Africa, Asia, and Australia), continental fragments (India, Madagascar, the Mascarene Plateau including Seychelles), continental islands (many in the Indo-Australian Archipelago, IAA) and oceanic islands (Comoros, the Mascarene Islands, and the IAA), which sum up to high geological complexity. Due to its geological and environmental characteristics, the IO has been a prolific natural laboratory for studies on continental drift (see ref. ^1^), climatology (e.g. ref. ^2^), geology (e.g. ref. ^3^) and, of course, biogeography (e.g. ref. ^4^).

When Pangaea was splitting into Laurasia and Gondwana, at 160 Ma (millions of years ago), a series of break-up events would trigger the IO’s formation: Australia and India first separated from the rest of Gondwana (Fig. 1a) and, in the Early Cretaceous, from each other (140 Ma; Fig. 1b); meanwhile, India carried Madagascar northwards (Fig. 1c) until their break-up some 90 Ma (Fig. 1d)^5,6^. As India drifted northwards, it affected oceanic circulation, the climate, and the shape of the continents, such as through collision with Eurasia – which, for instance, uplifted the Oman-Kohistan-Dras Island Arc (OKD) (Fig. 1d,e)^1,7,8^. With the northward movement of the Australian Plate at ~14 Ma, the IO was re-organised and the Miocene Indian Ocean Equatorial Jet (MIOJet), a strong westward current, was in place and lasted for over 10 Myr (Fig. 1f)^9^. The IAA originated through the movement of the Australian Plate towards the Sunda Shelf, modifying landscapes and oceanic circulation (Fig. 1e)^5,10^. Around the Cretaceous–Palaeogene boundary (K-Pg), the Mascarene Plateau separated from India (Fig. 1e)^11^. The northern part of the plateau (including Seychelles) is a granitic 750 million years (Myr) old micro-continent^12^; the southern part (including the Mascarene Islands, which began their emergence in the Eocene) is a considerably younger volcanic trace of the Réunion hotspot^13^.

**Figure 1.**
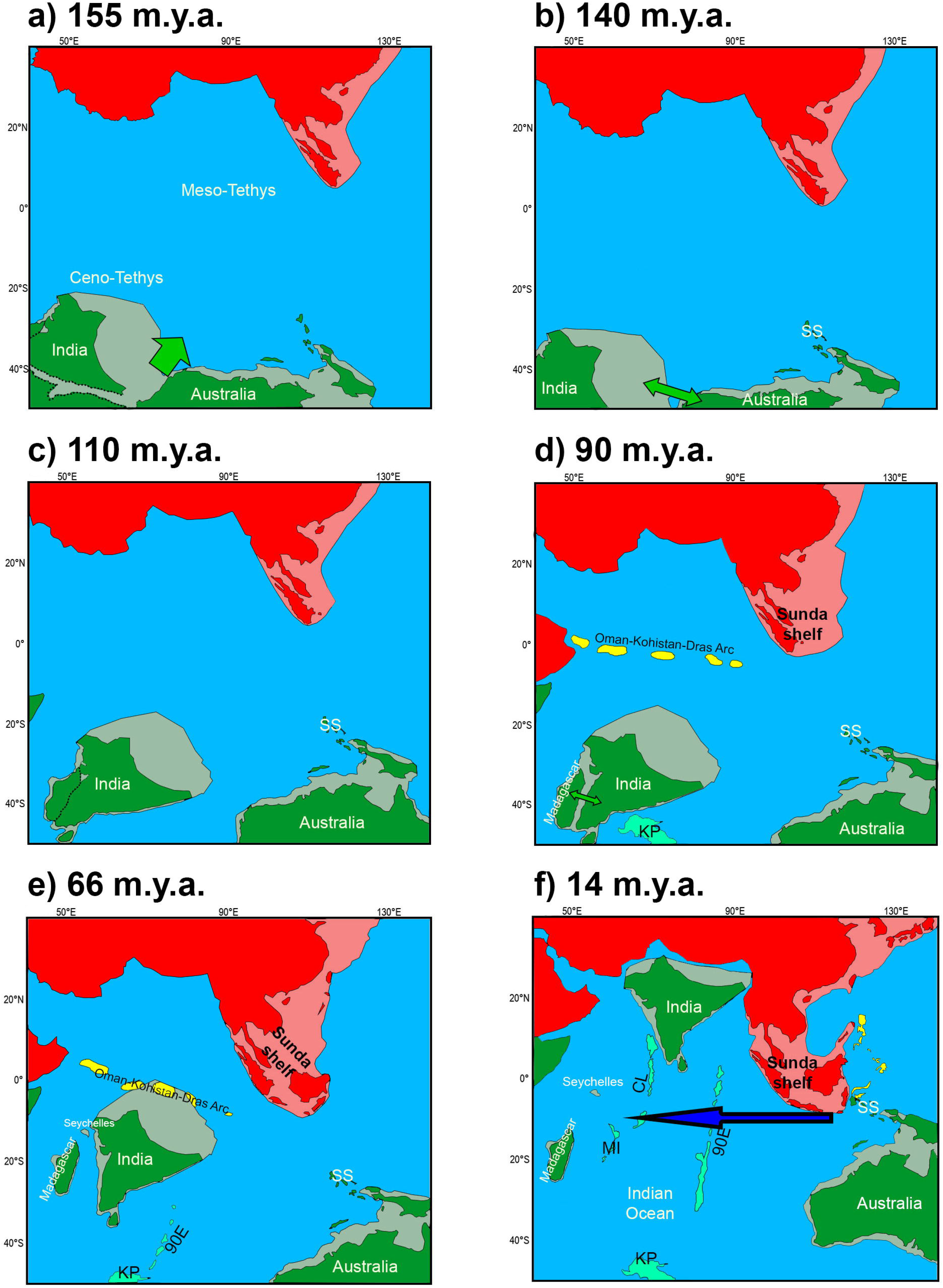
Geologic reconstruction of the Indian Ocean’s continents through the Mesozoic and part of the Cenozoic at 155 Ma (a), 140 (b), 110 (c), 90 Ma (d), 66 Ma (e), and 14 Ma (f). Laurasian landmasses are represented in red and Gondwanan landmasses in green. Areas in yellow are volcanic islands and in cyan are volcanic ridges (90E: Ninetyeast Ridge; CL: Chagos-Laccadive Ridge; MI: Mascarene Islands) and plateaus (KP: Kerguelen Plateau). The lightly coloured areas next to the continents represent continental shelves, which may or may not have been emergent depending on the timeframe. Dashed lines represent boundaries of continental break-ups; green arrows represent major break-up events; and the blue arrow represents the MIOJet. SS: Sula Spur. Redrawn from refs. ^1,5,8^.

Several major biogeographical regions converge on and appear to be interconnected by the IO (Australasian, Oriental, Afrotropical, Madagascan, Saharo-Arabian, and even Oceanian). They have been subjects for many studies in biogeography, especially after the emergence of vicariant theory and panbiogeography, and the oceanic dispersal paradigm^14^. For instance, while Madagascar has one of the highest current levels of endemism in the world^15^, India, despite over 100 Myr of isolation, is the focus of a longstanding debate regarding the lack of endemicity^16^. Both landmasses present mixed (Gondwanan and Laurasian) taxa^16,17^, which brought theoretical land bridges such as the so-called ‘Lemurian stepping-stones’ into the debate^18^ (Fig. 1d).

Divergence time inferences and molecular phylogenies have presented a turn of the tide in the vicariant-dispersal dichotomy, but very few works have looked past the continental drift problem. Even extensive systematic reviews failed to evaluate the timing of the events, and, as the methods developed, molecular clock calibration approaches have been rarely reviewed across taxonomic analyses (e.g. ref. ^4,19^. New bridges and stepping-stones are constantly proposed and often evoked as explanations for current taxon distributions, but the rationale they provide may be more problematic to parse than that of long-distance dispersal^20,21^. Furthermore, more recent methods such as ancestral area reconstructions remain largely absent from comprehensive studies of IO biogeography. Here, we review and evaluate the different pressures and geological and ecological conditions that enabled the colonisation of the Indian Ocean region through time. We also assess the role of the proposed bridges and stepping-stones by re-examining the time frame of migration events and ancestral range evolution across taxa.

## Materials and Methods

### Study selection

We conducted literature searches for potential studies to be used in this meta-analysis. Four selection criteria for inclusion were identified *a priori:* (1) a focus on either terrestrial or freshwater taxa; (2) inclusion of molecular genetic data; (3) a focus on phylogenetic or phylogeographic reconstructions; and (4) inclusion of at least five of the biogeographically relevant areas delimited here: continental Africa, continental Asia, Australia, Comoros, India, IAA, Madagascar, Mascarene Islands, and Seychelles.

Various studies have focused on the West Indian Ocean, Madagascar, India or IAA. We, on the other hand, wanted to identify what kind of patterns can be seen in a broader sense. Thus, the above criteria aimed to identify changes in trends along a phylogeny, and to then examine the concordance of trends associated with various time frames across different phylogenies. We chose studies that allowed us to (1) replicate the earlier phylogenetic reconstructions – since we needed the raw phylogenetic trees as input for ancestral range inference – and (2) allowed for calibration of molecular clocks (see below).

Initially, 54 studies fit our search criteria. Studies that were excluded and justification for exclusion can be seen in Appendix S1. Four studies were excluded initially because we were not able to replicate the original analyses and/or could not apply a reasonable calibration strategy. Twenty-three studies were included in the final dataset (Table 1). Some of the datasets were supplemented with information from other studies to fill taxonomic and/or geographic gaps (Appendix S2).

**Table 1.** Taxa included in this study with a summary of the calibration method for the molecular clock, type and number of markers used, divergences dates. Ages are the most recent common ancestor for the given taxon. References of the articles on which the analyses were based are available in the Appendix S0. Some of the analyses include data from other sources and these are specified in Appendix S2.

| Taxon | Taxonomic level | Calibration method | Markers |  | Taxon age in m.y.a. (95% CI) |  |
| --- | --- | --- | --- | --- | --- | --- |
|  |  |  | mtDNA/cpDNA | nDNA | Source | This study |
| Aves: Acrocephalidae | Family | Substitution rate | 3 | 5 | 14 (12-16) | 8 (6-9) |
| Aves: Anatini | Tribe | Fossil | 5 | 0 | ~18 | 22 (18.5-26) |
| Aves: Falconidae | Family | Fossil | 0 | 8 | 28 (23-35) | 34 (29-38) |
| Aves: Palaeognathae | Infraclass | Fossil | 1* | 0 | 73 (63-84) | 80 (73-89) |
| Aves: Psittacoidea | Superfamily | Fossil | 3 | 4 | 59 (45-72) | 50 (40-60) |
| Aves: Pycnonotidae | Family | Substitution rate,<br>Secondary calibration | 7 | 7 | – | 11 (9-13) |
| Crustacea: Atyidae | Family | Fossil,<br>Secondary calibration | 1 | 1 | 161 (69-257) | 141.5 (125-158) |
| Lissamphibia: Caecilia | Order | Fossil | 3 | 6 | ~109 | 178 (174-184) |
| Lissamphibia: Neobatrachia | Suborder | Fossil | 0 | 95** | 128 (120-135) | – |
| Mammalia: Emballonurinae | Subfamily | Fossil | 1 | 1 | 56 (47.5-69) | 54 (47-63) |
| Mammalia: Strepsirrhini | Suborder | Fossil | 2 | 4 | 61 (55-67) | 58 (50-65) |
| Mammalia: Pteropodidae | Family | Substitution rate,<br>Secondary calibration | 2 | 4 | 8 (5.5-11) | 7 (5-8)<sup>2</sup> |
| Squamata: Boidae | Family | Fossil | 1 | 5 | 77 (68-89) | 72 (67-79) |
| Squamata: Chamaeleoninae | Subfamily | Fossil | 3 | 10 | 65 (55-75) | 66 (53-75) |
| Squamata: Gekkonidae | Family | Fossil,<br>Secondary calibration | 3 | 4 | – | 100 (97-103) |
| Squamata: Natricinae | Subfamily | Fossil | 6 | 1 | 33 (28-40) | 56 (48-66) |
| Squamata: Typhlopoidea | Superfamily | Fossil | 0 | 5 | 106 (91-121) | 108 (93-124) |
| Testudines: Testudinidae | Family | Fossil | 3 | 2 | – | 50 (37-66) |
| Angiosperms: *Aphananthe* | Genus | Fossil | 4 | 0 | 19 (12-27.5) | 13 (9-16.5) |
| Angiosperms: *Exacum* | Genus | Secondary calibration | 2 | 0 | 21 (8-36) | 41 (32-50) |
| Angiosperms: *Ficus* | Genus | Fossil | 0 | 4 | 70 (60-92.5) | 68 (60-82) |
| Angiosperms: Loranthaceae | Family | Fossil | 3 | 2 | 59 (53-66) | 72 (71-72.5) |
| Angiosperms: Rubiaceae | Family | Secondary calibration<sup>1</sup> | 4 | 0 | 25 (30-21) | 25 (28-20) |

\* The marker used was the entire mitogenome;
\*\* This dataset was very large and, thus, we had to reduce and restrain it to set up the analysis (see Appendices);
<sup>1</sup> To simplify the analysis, we separated the tribes that were present in diverse biogeographic regions (Coffeae, Bertiereae, Octotropideae, and Pavetteae) and built the trees independently;
<sup>2</sup> Age of TMRCA between *Pteropus* and *Acerodon*.

### Phylogenetic analyses

All phylogenetic analyses were conducted in either BEAST or *BEAST as implemented in BEAST2 2.4.0 (*BEAST was used when there was allele and/or sampling heterogeneity within a single species)^22^, with the primary aim to replicate the original phylogenetic analyses. Due to the large number of analyses we conducted, we simplified some parameters to facilitate convergence. Because of the abovementioned issues, we established the following pipeline for our analyses:

#### (I) Data handling

we used all information made available by the authors in our first approach to the data (alignment, partitioning, priors, etc.). Nonetheless, we altered what we did not consider as the best practice for data analysis (Appendix S2).

#### (II) Substitution model

we used a reversible-jump method to choose the best model^23,24^. The reversible-jump method was chosen because it (1) facilitates convergence by finding the least parametrised model to fit the data and (2) makes phylogenetic estimation independent of a specific model by integrating it over the parametric uncertainty. However, our previous experience shows that better convergence is obtained when offset values for the model parameters are empirically obtained through jModelTest 2.1^25^; thus, we combined both methods.

#### (III) Tree priors

both Yule and birth-death tree priors were used for all analyses, but no significant differences were found. The priors were all initially set to a lognormal distribution. The distribution and parameter values of the priors were then modified when needed according to assessments made through Tracer 1.6^26^. Analyses of datasets comprising multiple sequences per species terminal were run with all gene trees unlinked, using a coalescent model for species trees inference^27^. Otherwise, it was assumed that concatenated gene trees correspond to species trees, despite separate substitution models being applied to each gene.

#### (IV) Estimation of divergence times

all analyses were run with an uncorrelated lognormal relaxed-clock model. Although the relaxed and strict models are commonly tested against each other, the accuracy of such statistical comparisons has been debated and, currently, seems to be accompanied by computational intricacy^28^. Since the strict clock is a special case of the relaxed clock where there is no variation of the rates along the branches, it is reasonable to start with a relaxed model and only test for the strict model those markers that presented the median of the standard deviation’s distribution tending to zero. Tests were made through a simple AICM approach^29^.

#### (V) Calibration

we sought available molecular-clock calibration methods from the literature. Calibration methods of three types were identified to be informative: fossil, molecular (substitution) rates, and secondary calibrations obtained from broader phylogenetic studies. Geological events such as island emergence were not used to calibrate the age of an island clade to avoid circular reasoning^19^. We modified prior settings where necessary to allow soft constraints on the calibrated node^30^.

Alignments were done through MAFFT 6^31^, plus visual alignment (using secondary structure) for ribosomal markers^32^. All Bayesian phylogenetic analyses were run for 100-500 million generations sampling every 10-50 thousand generations. Runs were visually assessed in Tracer with a 25% burn-in. Final maximum clade credibility trees were obtained through TreeAnnotator (BEAST package) by using the same burn-in. All but the phylogeographic analyses were run on the CIPRES Science Gateway web portal^33^ (Appendix S2).

### Biogeographic analyses

We inferred ancestral areas in RASP 4.0^34^, which also implements BioGeoBEARS 1.1^35^. Species and population distributions were obtained either from the original papers or the literature. These localities were designated to one or more of the areas defined above. The model testing implemented by BioGeoBEARS was also used, but since it only accounts for the ‘model-like’ approaches implemented by the software, we ran the original versions of DEC^36^ and BayArea^37^ as well as the similar approaches implemented by BioGeoBEARS. To avoid the limitations imposed by DiVA^36^, we only used BioGeoBEARS’ DIVALIKE method. BayArea was run both with and without coordinates (Appendix S3) to identify incongruences, for 20 million generations sampling every 4,000; both DEC and BayArea were run with unconstrained settings for dispersal with a maximum range size of four areas.

## Results

### Phylogenetic analyses

We were able to recover similar topology and divergence times compared to the original studies for 14 datasets (Table 1). Some of the datasets were largely altered in terms of terminals and priors (Appendix S2) and it led to major differences in the topology and/or divergence times.

### Biogeographic analyses

Based on the BioGeoBEARS model test as implemented in RASP, the *j* parameter had a positive effect on every dataset’s AIC, except for the Strepsirrhini (lemurs). However, even in the lemur dataset, the *j* parameter was not completely discarded (Table 2). This parameter models speciation caused by long-distance dispersal and it has been shown to have a strong effect over other parameters used in classic approaches to biogeographic reconstruction^35^. DEC+*j* was the most commonly favoured model although with some notable exceptions (Table 2).

**Table 2.**
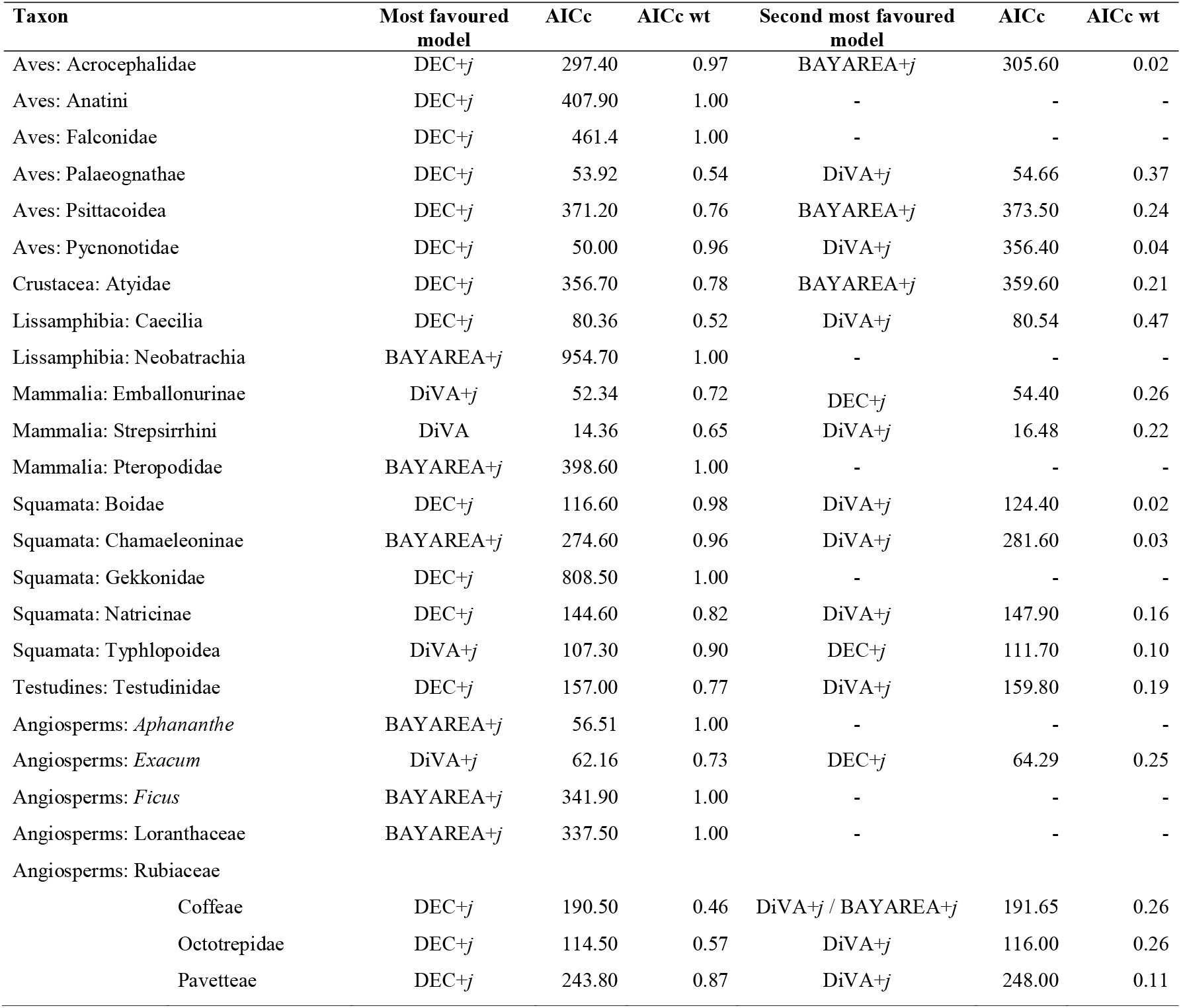
Summary of the BioGeoBEARS model test results per taxon/dataset as well as AICc/AICc weights for each model considered.

Figure 2 summarises vicariance and dispersal events as inferred from the ancestral area estimation, assigned to six relevant timeframes: Cretaceous including most of the Gondwanan break-up (82–71 Ma); the K-Pg Boundary (70–60 Ma); Palaeocene and Early Eocene when many of the proposed bridges were in place (57–46 Ma); Late Eocene when there were multiple ecological and geological shifts (42–32 Ma); Oligocene and Early Miocene when there were dramatic landscape and oceanic circulation changes (31–20 Ma); and Neogene (19–5 Ma). Two of the analysed taxa (Pycnonotidae and Falconidae) are not represented in Figure 2 because they have a very recent biogeographic history with dispersal events dated <4 Ma. The Comoros are also not represented in Figure 2 for the same reason.

**Figure 2.**
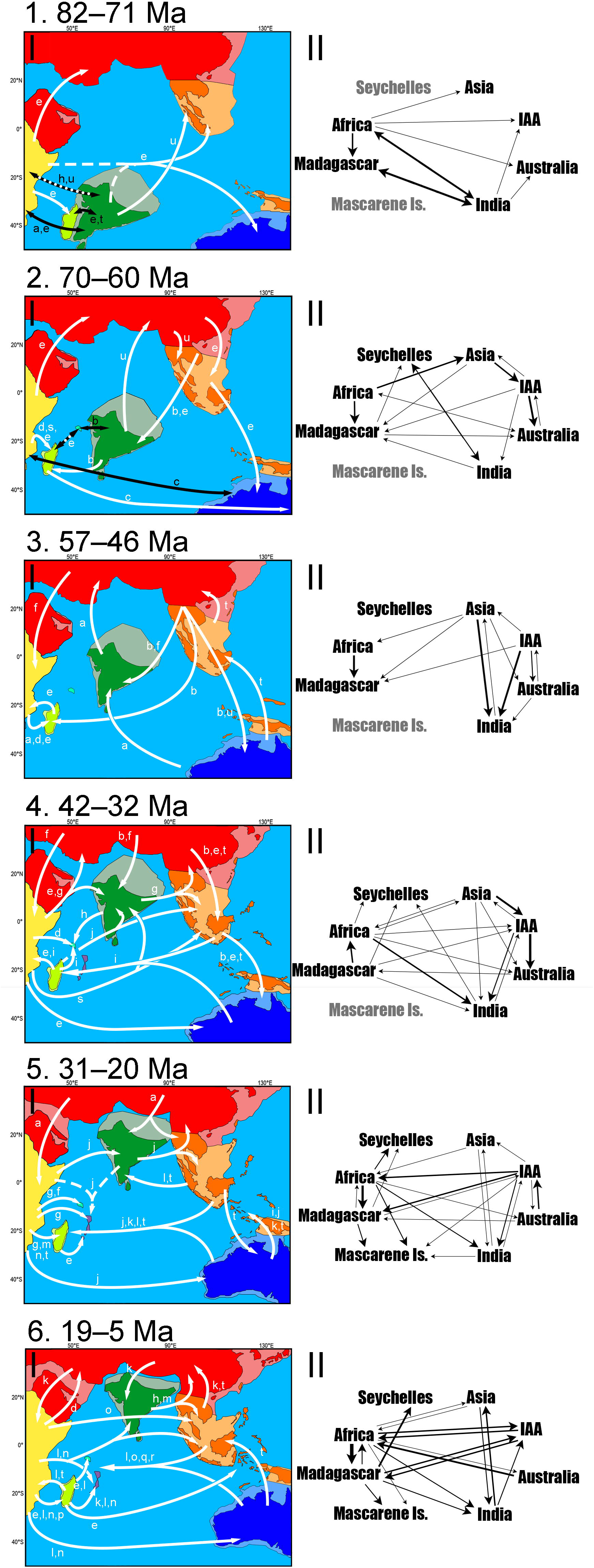
Dated results from BayArea and DEC at ~80 Ma (1), ~66 Ma (2), ~55 Ma (3), ~40 Ma (4), ~25 Ma (5), and ~15 Ma (6). (I) Mapped colonisation routes by taxon. Represented Biogeographic regions are colour-coded: Africa in yellow, Asia in red, Australia in blue, IAA in orange, India in green, Madagascar in light green, MI in purple and Seychelles in cyan (lighter colours represent continental shelves). Taxa represented are Boidae (a), Neobatrachia (b), Palaeognathae (c), Chamaleoninae (d), Gekkonidae (e), Natricinae (f), Testudinidae (g), Caecilia (h), Emballonuridae (i), Ficus (j), Psittacoidea (k), Atyidae (l), *Exacum* (m), Rubiaceae (n), Pteropodidae (o), Acrocephalidae (p), Anatini (q), *Aphananthe* (r), Strepsirrhini (s), Loranthaceae (t), and Typhlopoidea (u). White arrows represent dispersal and black arrows represent vicariance. Dashed arrows represent competing ancestral areas that may have composed an entire ancestral range. Note that the white and black arrow in (1) represents the uncertainty between vicariance and dispersal due to the confidence interval of the nodal dates. The map only illustrates relative positions and does not show the precise size and shapes of the continents. Based on refs. ^1,5,8^. (II) Total weighted colonisation routes. One-headed arrows represent dispersal and two-headed arrows represent vicariance. Line thickness corresponds to the number of colonisations. Regions with their names in grey were not present at the given timeframe. The arrows do not have any specific order inside each represented timeframe. For specific dates or taxa, please refer to Appendix S4.

According to our dates, few of the dispersal routes presented in Figure 2 could have relied on the stepping stones found in the biogeography literature. Cretaceous events in the IO are mainly Gondwanan by default, but long-distance dispersal increased around the K-Pg boundary (Fig. 2.2). Exchanges with Laurasia increased through the Cenozoic, particularly via the IAA (Fig. 2.3 and 2.4). These exchanges increased through time, with shifts in which landmasses served as centres of origin.

The analysed taxa exhibit quite different migration pathways, although they converge to similar locations in similar periods (see Discussion). Although it appears unlikely that one or more routes persisted for extended periods, a few trends can be seen in Figure 2: India has served as a stepping stone both between eastern and western IO and between Laurasia and Gondwana; Africa and Asia persisted as a centre of origin for most of the Cenozoic; the IAA increased exchange between Asia and Australasia until the collision of Sundaland and the Sula Spur. In the Late Oligocene, a westward route persisting through the Miocene is supported by the data (Fig. 2.5); by the end of the Miocene, eastward migration increased and a bidirectional pattern was in place (Fig. 2.6).

## Discussion

Previous studies have shown that, despite the IO’s long and complex geological history, the importance of transoceanic dispersal pre-dates the establishment of the IO even during the flourishing continental break-up in the Mesozoic. However, due to the IO’s composite nature, involving many continents, subcontinents and islands, most of the studies are restricted to smaller sections of the IO, such as single regions or eastern/western provinces or islands (e.g. refs. ^4,16,38^).

The review by Agnarsson and Kuntner^4^ is an exemplary of such trends. This is one of the most extensive and robust reviews of the Western IO islands (Madagascar, Seychelles, Mascarenes, Comoros, Zanzibar and Socotra) including about 100 studies. However, the authors themselves underline that they have used no information on the timing of the biogeographic events as such data was scarce at the time. Although there are some strong similarities between their Figure 2 and our results on the relationships between the regions examined here, the complexity of those relationships remains largely underestimated.

Here, we have a much wider biogeographic focus than most of the previous studies, but, more importantly, we focussed on the timing of trends and estimation of ancestral range evolution to find common patterns across geological ages. Our results, rather than another nail in the coffin of the vicariance vs. dispersal debate, illustrates how taxa spread across the IO since its formation in the Mesozoic as well as the drivers of biogeographic events.

The Late Cretaceous represented a warm environment with lower sea levels^39^ and the K-Pg extinction event created a transforming environment with ample opportunity for community turnovers^40,41^. The shallow shelves and exposed land bridges left plenty of pathways connecting Gondwanan fragments^42,43^ and westward sea currents flowing through SE Laurasia^44^ could have facilitated long-distance dispersal by, e.g., ratites, lemurs, and herpetofauna (Fig. 2.1; 2.2), as well as abelisaurid dinosaurs^45^.

The biogeographical history we reconstructed for herpetofauna and angiosperms (Fig. 2.3a,b,f,t,u) concur with previously documented biotic interchanges between Laurasia and Gondwana. This pattern is particularly true for Afro-Indian flora colonisation in Asia^46^. For a long time, there was little evidence for some of the landmasses related to any Laurasia-Gondwana routes across the IO^8^ being subaerial at the time, but recent reconstructions of paleotopography show solid pathways, e.g., through the OKD^43^. However, the presence of these routes alone does not explain the timing of the biotic exchanges. Alternatively, we argue that the thermal maximum at the Palaeocene-Eocene boundary (PETM)^47^ might have facilitated the establishment of some taxa not only by exposing stepping-stones through oscillations in sea level^48^ but also by triggering migrations and prompting new suitable environments via climatic changes.

Similarly, our results replicated a known interchange between Asia and India^8,46^ with an intensification of migrations post-Middle Eocene (Fig, 2.4). Considering the confidence interval for the dates, these events could be related to the Middle Eocene climatic optimum^49^ and/or to the Eocene-Oligocene extinction event^50,51^. Unlike the PETM, the Middle Eocene climatic optimum was a much slower and longer-lasting global warming period^49^ and led to a cooling period that, among other things, triggered the Antarctic glaciation^52^. The cooler global climate caused aridification of the landscape and expansion of seasonal vegetation across India and the IAA^46,53^. This could have offered some habitat heterogeneity and new ecological opportunities in the tropics^54^ while turning Asia into a faunal centre of origin^55^ (Fig. 2.4b,e,f,t). Moreover, the transformations in the Antarctic lowered sea levels and could have exposed elements of the OKD and IAA that allowed the colonisation of the South Hemisphere lowlands.

Madagascar, which had both a long-lasting link with Africa^56,57^ and a more puzzling relationship with Asia/India^38^, has presented some interesting results across the Eocene: the Palaeocene trend of colonisation by African taxa is maintained in the Palaeocene-Eocene boundary (Fig. 2.3). This trend is, then, reversed during the Eocene (Fig. 2.4) and reverted again in the Oligocene (Fig. 2.5). The availability of pathways between Africa and Madagascar coincides with the periods when Madagascar received immigrants^56^, but it does not offer any clarification for the reversal during the Eocene. If we look at the availability of ecological opportunities, on the other hand, these three periods fit three different climatic profiles: the PETM, the Middle Eocene climatic optimum and the Eocene-Oligocene cooling triggered by the Antarctic glaciation^52,55^. These changes probably caused biotic turnovers^40,58^ aided by Madagascar’s northward movement, thereby leaving the influence of the subtropical arid belt and becoming wetter. The increase in the diversification rate caused by this transformation could explain the brief emigration wave from Madagascar. The Eocene-Oligocene extinction could also be related to the faunal transformation and intense immigration observed in the Oligocene^59^, but the floral signature in the transformation is very clear in our results (Fig. 2.5).

The IAA, with a close association with Asia and Australasia, functioned as a pathway before it had a unique biogeographic identity. Nonetheless, around the Oligocene-Miocene boundary (Fig. 2.5), the IAA’s heterogeneous geological and climatic history plus the high number of geologically independent islands allowed intense diversification and endemicity that would transform it into a centre of origin^53^. The changes in sea level and oceanic circulation created a transoceanic route between Australia-IAA and the Western IO that might have been intensified by the MIOJet. At the time, the Mascarene Islands were emerging and one could argue that they could have aided such a trend by serving as a stepping-stone, but, according to our reconstruction, they seem to have functioned as a sink (Fig. 2.5j,k,l,t; Fig. 2.6l,o,q,r).

In the Late Oligocene, the Antarctic thawing caused rapid changes in sea level^52^ and was followed by a cooling period that led to the expansion of the grasslands and the appearance of more heterogeneous ecosystems^64^. Both India and Madagascar had their ecology vastly transformed due to their climatic instability during the Eocene and again in the Miocene^17,40^. Such instability was likely accentuated by the latitudinal changes in these two islands and might have favoured some groups of angiosperms (e.g. Fig. 2.5j,t) and allowed the faunal composition to be reshaped. The ‘Asian flavour’ in the endemic biota of these two landmasses was acquired very early in the Cenozoic and accentuated through time^16,38^.

While the distinction between eastern and western IO became clearer as the IAA was formed and the colonisation of the western IO increased in the Miocene, incursions into India remained common (Fig. 2.3-6). The extant endemicity in India sometimes conflicts with a fossil record teeming with non-Indian taxa and several land bridges have been invoked as an explanation (e.g. refs. ^1,8,60^). However, shaped by geological and latitudinal changes, India was a transforming environment that represented either a source or a recipient at different times^46,61^. Routes between India and Gondwana were constantly available until at least the Eocene (maybe Oligocene) and were still present when the connection with Asia became wider^43^. These connections and the northward movement of India that put it in the middle of Cenozoic oceanic currents between Eastern and Western IO^9,62^, turned India into some sort of gatekeeper both between east and west and between Laurasia and Gondwana. Accordingly, this role was gradually diminished as India became fully connected to Asia. Like an ark, immigrants could colonise Asia more effectively after docking^6^.

BioGeoBEARS’ new parameter to measure founder-effect speciation (*j*) was favoured in all but one of our analyses (namely, the lemurs; Fig. 2s). This is interesting because, without founder-effect, we are left with either dispersal followed by extinction or vicariance. According to our analysis, the Strepsirrhini originated in the Late Cretaceous and diversified near the K-Pg boundary. This dating suggests that lemurs might have had a Gondwanan ancestor and reached Madagascar through vicariance either from Africa or from India, or possibly had a more pan-Gondwanan distribution followed by extinction outside Madagascar. This result underlines the importance of ancestral range reconstruction rather than most parsimonious approximations derived from time-calibrated phylogenies.

The biogeographical reconstruction of these regions is difficult for two main reasons. First, extinction has been neglected in biogeography^19^ and the observed increasing trend of dispersal events through time (Fig. 2) may be an artefact of not including extinct taxa in the analyses. Secondly, the further back we go into Earth’s history, geography becomes more challenging to reconstruct^1,57,60^. While geology and palaeontology present us with some well-recorded events and the timing thereof, evidence from alternative sources has driven debates about dates, locations, and causes of extant biodiversity patterns (e.g. ref. ^7^). A record of major evolutionary events is sometimes kept by the fossil assemblages and sometimes retrieved by the molecular data, but, since biotic turnovers are dependent on the size and isolation of the island, high extinction rates may be why we were not able to infer as many ancient colonisation events on smaller ancient islands such as the Seychelles.

This adds to the debate on the ability of different taxa to cross different sections of the IO^14,65,66^. Although biogeographers historically resorted to stepping-stones, the necessity of such paths for colonisation of isolated islands has already been brought to question^20^. Here, we present strong evidence for only two systems: IAA elements (between Asia and Australia) and Indian elements (between Asia/IAA and western IO) (Fig.2). Other paths involving stepping-stones that are no longer available are hard to assess due to the extinct intermediate populations. Like the Krakatau case study, there might have been divergent environments that would not allow colonisation by similar populations^20^. Alternatively, sea and air currents could have served as constant pathways for rafts and migratory animals and/or phoresis and the establishment of the migrants would then be regulated by niche availability.

Thus, the establishment of the taxa might be equally or even more relevant than the mechanism by which they got there in explaining present-day biogeography: ecological transformations may contribute significantly to presenting opportunities for migration and establishment, whereas biological properties such as life-history strategies may account for differences between organisms in vagility or competition. Here, we managed to reconstruct timely trends that concurred across different taxa and coincided with known environmental transformations. Further research could investigate specific cases both across time and across geography.

## Conclusion

The role of continental vicariance is ancient and has left little signal in the present-day Indian Ocean biota. The IO has been constantly shaped by geological and climatic changes, with transforming landscapes and ongoing ecological opportunities. Although its continents, particularly the Gondwanan landmasses, have been widely studied and debated, they have been examined in terms of ‘how did they get there,’ which is certainly important but neglects the relevance of ecological factors and evolutionary events over dispersal mode. Here, we argue that the ‘why were they able to establish there’ should be looked at more carefully. Future research should attempt greater taxonomic breadth, not only in the sense of including more taxa but also in the sense of better assessing key features of the biota such as extinction. These studies should also aim for more depth: molecular dating and biogeographic models have advanced greatly as well as paleogeographic and paleoclimatic reconstructions. Our results show that making use of these advances can challenge previous hypotheses based on parsimony. Furthermore, the Indian Ocean’s biogeographic history is intrinsically connected to the tropics, where older taxonomic diversity and higher speciation rates provide ecological diversity and the origins of many non-tropical lineages.

## Acknowledgements

We warmly thank Dr Björn Stelbrink and Dr Nicholas Matzke for discussions leading to major insights in this study. We also thank the reviewers, whose comments have been valuable to enrich this work. The authors declare that they have no conflicts of interest regarding this publication.

## Data availability statement

Data analysed and produced by this study can be downloaded from Dryad repository. This includes: Appendix S1 with the justification for non-inclusion of the studies in this meta-analysis; Appendix S2 with all the parameters and details for the phylogenetic and phylogeographic analyses; Appendix S3 with the coordinates used in the phylogeographic analyses; and Appendix S4 with the final trees and ancestral ranges reconstructions.

