## Supplementary material S0 to S4 for "Ecological changes have driven biotic exchanges across the Indian Ocean": S0.pdf

### Appendix S0: Studies used as a basis for the main analyses


1. Aves: Acrocephalidae

   Arbabi T., Gonzalez J., Wink M. 2014. A re-evaluation of phylogenetic relationships within reed warblers (Aves: Acrocephalidae) based on eight molecular loci and ISSR profiles. *Mol. Phylogenet. Evol.* **78**:304–313.

2. Aves: Palaeognathae

   Mitchell K.J., Llamas B., Soubrier J., Rawlence N.J., Worthy T.H., Wood J., Lee M.S.Y., Cooper A. 2014a. Ancient DNA reveals elephant birds and kiwi are sister taxa and clarifies ratite bird evolution. *Science.* **344**:898–900.

3. Aves: Anatini

   Mitchell K.J., Wood J.R., Scofield R.P., Llamas B., Cooper A. 2014b. Ancient mitochondrial genome reveals unsuspected taxonomic affinity of the extinct Chatham duck (*Pachyanas chathamica*) and resolves divergence times for New Zealand and sub-Antarctic brown teals. *Mol. Phylogenet. Evol.* **70**:420–428.

4. Aves: Falconidae

   Fuchs, J., Johnson, J. A., Mindell, D. P. 2015. Rapid diversification of falcons (Aves: Falconidae) due to expansion of open habitats in the Late Miocene. *Mol. Phylogenet. Evol.* **82(A)**:166–182.

5. Aves: Psittacoidea

   Schweizer M., Seehausen O., Hertwig S.T. 2011. Macroevolutionary patterns in the diversification of parrots: effects of climate change, geological events and key innovations. *J. Biogeogr.* **38**:2176–2194.

6. Aves: Pycnonotidae

   Shakya S.B., Sheldon F.H. 2017. The phylogeny of the world's bulbuls (Pycnonotidae) inferred using a supermatrix approach. *Ibis.* **159**:498–509.

7. Crustacea: Atyidae

   von Rintelen K., Page T.J., Cai Y., Roe K., Stelbrink B., Kuhajda B.R., Iliffe T.M., Hughes J., von Rintelen T. 2012. Drawn to the dark side: A molecular phylogeny of freshwater shrimps (Crustacea: Decapoda: Caridea: Atyidae) reveals frequent cave invasions and challenges current taxonomic hypotheses. *Mol. Phylogenet. Evol.* **63**:82–96.

8. Crustacea: Atyidae

   Bernardes S.C., Pepato A.R., von Rintelen T., von Rintelen K., Page T.J., Freitag H., de Bruyn M. 2017. The complex evolutionary history and phylogeography of *Caridina typus* (Crustacea: Decapoda): long-distance dispersal and cryptic allopatric species. *Sci. Rep.* **7**:9044.

9. Lissamphibia: Caecilia

Pyron R.A. 2014. Biogeographic analysis reveals ancient continental vicariance and recent oceanic dispersal in amphibians. *Syst. Biol.* **63**:779–797.

10. Squamata: Gekkonidae

    Pyron R.A., Burbrink F.T., Wiens J.J. 2013. A phylogeny and revised classification of Squamata, including 4161 species of lizards and snakes. *BMC Evol. Biol.* **13**:93.

11. Lissamphibia: Neobatrachia

    Li J.-T., Li Y., Klaus S., Rao D.-Q., Hillis D.M., Zhang Y.-P. 2013. Diversification of rhacophorid frogs provides evidence for accelerated faunal exchange between India and Eurasia during the Oligocene. *PNAS.* **110**:3441–3446.

12. Lissamphibia: Neobatrachia

    Feng Y.-J., Blackburn D.C., Liang D., Hillis D.M., Wake D.B., Cannatella D.C., Zhang P. 2017. Phylogenomics reveals rapid, simultaneous diversification of three major clades of Gondwanan frogs at the Cretaceous–Paleogene boundary. *PNAS.* **114**:E5864–E5870.

13. Mammalia: Emballonurinae

    Ruedi M., Friedli-Weyeneth N., Teeling E.C., Puechmaille S.J., Goodman S.M. 2012. Biogeography of Old World emballonurine bats (Chiroptera: Emballonuridae) inferred with mitochondrial and nuclear DNA. *Mol. Phylogenet. Evol.* **64**:204–211.

14. Mammalia: Strepsirrhini

    Herrera J.P., Dávalos L.M. 2016. Phylogeny and divergence times of lemurs inferred with recent and ancient fossils in the tree. *Syst. Biol.* **65**:772–791.

15. Mammalia: Pteropodidae

    O'Brien J., Mariani C., Olson L., Russell A.L., Say L., Yoder A.D., Hayden T.J. 2009. Multiple colonisations of the western Indian Ocean by *Pteropus* fruit bats (Megachiroptera: Pteropodidae): The furthest islands were colonised first. *Mol. Phylogenet. Evol.* **51**:294–303.

16. Mammalia: Pteropodidae

    Almeida F.C., Giannini N.P., DeSalle R., Simmons N.B. 2011. Evolutionary relationships of the old world fruit bats (Chiroptera, Pteropodidae): Another star phylogeny? *BMC Evol. Biol.* **11**:281.

17. Mammalia: Pteropodidae

    Almeida F.C., Giannini N.P., Simmons N.B., Helgen K.M. 2014. Each flying fox on its own branch: A phylogenetic tree for *Pteropus* and related genera (Chiroptera: Pteropodidae). *Mol. Phylogenet. Evol.* **77**:83–95.

18. Squamata: Boidae

    Noonan B.P., Chippindale P.T. 2006. Dispersal and vicariance: The complex evolutionary history of boid snakes. *Mol. Phylogenet. Evol.* **40**:347–358.

19. Squamata: Chamaeleoninae

Tolley K.A., Townsend T.M., Vences M. 2013. Large-scale phylogeny of chameleons suggests African origins and Eocene diversification. *Proc. R. Soc. B.* **280**:20130184.

20. Squamata: Typhlopoidea

Vidal, N., Marin, J., Morini, M., Donnellan, S., Branch, W. R., Thomas, R., Vences, M., Wynn, A., Cruaud, C., and Hedges, S. B. (2010). Blindsnake evolutionary tree reveals long history on Gondwana. *Biol. Lett*. **6(4)**:558–561.

21. Squamata: Natricinae

Guo P., Liu Q., Xu Y., Jiang K., Hou M., Ding L., Alexander Pyron R., Burbrink F.T. 2012. Out of Asia: Natricine snakes support the Cenozoic Beringian dispersal hypothesis. *Mol. Phylogenet. Evol.* **63**:825–833.

22. Testudines: Testudinidae

Le M., Raxworthy C.J., McCord W.P., Mertz L. 2006. A molecular phylogeny of tortoises (Testudines: Testudinidae) based on mitochondrial and nuclear genes. *Mol. Phylogenet. Evol.* **40**:517–531.

23. Testudines: Testudinidae

Joyce W.G., Parham J.F., Lyson T.R., Warnock R.C.M., Donoghue P.C.J. 2013. A divergence dating analysis of turtles using fossil calibrations: an example of best practices. *J. Paleontol.* **87**:612–634.

24. Angiosperms: *Aphananthe*

Yang M.-Q., Li D.-Z., Wen J., Yi T.-S. 2017. Phylogeny and biogeography of the amphi-Pacific genus *Aphananthe*. *PLOS ONE.* **12**:e0171405.

25. Angiosperms: *Exacum*

Yuan Y.-M., Wohlhauser S., Möller M., Klackenberg J., Callmander M.W., Küpfer P. 2005. Phylogeny and biogeography of *Exacum* (Gentianaceae): A disjunctive distribution in the Indian Ocean basin resulted from long distance dispersal and extensive radiation. *Syst. Biol.* **54**:21–34.

26. Angiosperms: Rubiaceae

Kainulainen K., Razafimandimbison S.G., Wikström N., Bremer B. 2017. Island hopping, long-distance dispersal and species radiation in the Western Indian Ocean: historical biogeography of the *Coffeeae* alliance (Rubiaceae). *J. Biogeogr.* **44**:1966–1979.

27. Angiosperms: Loranthaceae

Liu, B., Le, C. T., Barrett, R. L., Nickrent, D. L., Chen, Z., Lu, L., Vidal-Russel, R. 2018. Historical biogeography of Loranthaceae (Santalales): Diversification agrees with emergence of tropical forests and radiation of songbirds. *Mol. Phylogenet. Evol.* **124**:199–212.

28. Angiosperms: *Ficus*

Chantarasuwan B., Rønsted N., Kjellberg F., Sungkaew S., van Welzen P.C. 2016. Palaeotropical intercontinental disjunctions revisited using a dated phylogenetic hypothesis with nearly complete species level sampling of Ficus subsect. *Urostigma* (Moraceae). *J. Biogeogr.* **43**:384–397.
