## Supplementary material S0 to S4 for "Ecological changes have driven biotic exchanges across the Indian Ocean": S1.pdf

### Appendix 1: Excluded works and justification for exclusion

Bauret, L., Gaudeul, M., Sundue, M. A., Parris, B. S., Ranker, T. A., Rakotondrainibe, F., … Rouhan, G. (2017). Madagascar sheds new light on the molecular systematics and biogeography of grammitid ferns: New unexpected lineages and numerous long-distance dispersal events. *Molecular Phylogenetics and Evolution*, *111*, 1–17. doi: 10.1016/j.ympev.2017.03.005

- Global biogeography with limited geographic coverage of the Indian Ocean region

Bauret, L., Rouhan, G., Hirai, R. Y., Perrie, L., Prado, J., Salino, A., … Gaudeul, M. (2017). Molecular data, based on an exhaustive species sampling of the fern genus *Rumohra* (Dryopteridaceae), reveal a biogeographical history mostly shaped by dispersal and several cryptic species in the widely distributed Rumohra adiantiformis. *Botanical Journal of the Linnean Society*, *185*(4), 463–481. doi: 10.1093/botlinnean/box072

- Global biogeography with limited geographic coverage of the Indian Ocean region

Bauret, L., Field, A. R., Gaudeul, M., Selosse, M.-A., & Rouhan, G. (2018). First insights on the biogeographical history of Phlegmariurus (Lycopodiaceae), with a focus on Madagascar. *Molecular Phylogenetics and Evolution*, *127*, 488–501. doi: 10.1016/j.ympev.2018.05.004

- Global biogeography with limited geographic coverage of the Indian Ocean region

Bourguignon, T., Lo, N., Šobotník, J., Ho, S. Y. W., Iqbal, N., Coissac, E., … Evans, T. A. (2017). Mitochondrial Phylogenomics Resolves the Global Spread of Higher Termites, Ecosystem Engineers of the Tropics. *Molecular Biology and Evolution*, *34*(3), 589–597. doi: 10.1093/molbev/msw253

- Global biogeography with limited geographic coverage of the Indian Ocean region

Buyck, B., Kauff, F., Eyssartier, G., Couloux, A., & Hofstetter, V. (2014). A multilocus phylogeny for worldwide *Cantharellus* (Cantharellales, Agaricomycetidae). *Fungal Diversity*, *64*(1), 101–121. doi: 10.1007/s13225-013-0272-3

- Too restricted inside the Indian Ocean region (includes less than five of the nine delimited areas)

Chacón, J., & Renner, S. S. (2014). Assessing model sensitivity in ancestral area reconstruction using Lagrange: a case study using the Colchicaceae family. *Journal of Biogeography*, *41*(7), 1414–1427. doi: 10.1111/jbi.12301

- Global biogeography with limited geographic coverage of the Indian Ocean region

Dee, R., Malakasi, P., Rakotoarisoa, S. E., & Grace, O. M. (2018). A phylogenetic analysis of the genus *Aloe* (Asphodelaceae) in Madagascar and the Mascarene Islands. *Botanical Journal of the Linnean Society*, *187*(3), 428–440. doi: 10.1093/botlinnean/boy026

- Too restricted inside the Indian Ocean region (includes less than five of the nine delimited areas)

Fernández, R. & Giribet, G. (2015). Unnoticed in the tropics: phylogenomic resolution of the poorly known arachnid order Ricinulei (Arachnida). *Royal Society Open Science*, *2*(6), 150065. doi: 10.1098/rsos.150065

- Global biogeography with limited geographic coverage of the Indian Ocean region

Forthman, M., & Weirauch, C. (2016). Phylogenetics and biogeography of the endemic Madagascan millipede assassin bugs (Hemiptera: Reduviidae: Ectrichodiinae). *Molecular Phylogenetics and Evolution*, *100*, 219–233. doi: 10.1016/j.ympev.2016.03.011

- Global biogeography with limited geographic coverage of the Indian Ocean region
- We were unable to replicate the author's results and we could not decide if the lack of morphological data in the original work or the use of non-conserved regions of ribosomal loci was responsible for this incongruence.

Giribet, G., Sharma, P. P., Benavides, L. R., Boyer, S. L., Clouse, R. M., De Bivort, B. L., … Schwendinger, P. J. (2012). Evolutionary and biogeographical history of an ancient and global group of arachnids (Arachnida: Opiliones: Cyphophthalmi) with a new taxonomic arrangement. *Biological Journal of the Linnean Society*, *105*(1), 92–130. doi: 10.1111/j.1095-8312.2011.01774.x

- Global biogeography with limited geographic and phylogenetic coverage of the Indian Ocean region

Gomard, Y., Cornuault, J., Licciardi, S., Lagadec, E., Belqat, B., Dsouli, N., … Tortosa, P. (2018). Evidence of multiple colonizations as a driver of black fly diversification in an oceanic island. *PLOS ONE*, *13*(8), e0202015. doi: 10.1371/journal.pone.0202015

- Too restricted inside the Indian Ocean region (includes less than five of the nine delimited areas)

Guilbert, E. (2012). Biogeography of the Cantacaderinae Stål (Insecta: Heteroptera: Tingidae) revisited. *Invertebrate Systematics*, *26*(3), 316–322. doi: 10.1071/IS12010

- Too restricted inside the Indian Ocean region (includes less than five of the nine delimited areas)

Haber, E. A., Kainulainen, K., Van Ee, B. W., Oyserman, B. O., & Berry, P. E. (2017). Phylogenetic relationships of a major diversification of Croton (Euphorbiaceae) in the western Indian Ocean region. *Botanical Journal of the Linnean Society*, *183*(4), 532–544. doi: 10.1093/botlinnean/box004

- Lacks a calibration method to date the phylogenies.

Harrison, S. E., Harvey, M. S., Cooper, S. J. B., Austin, A. D., & Rix, M. G. (2017). Across the Indian Ocean: A remarkable example of trans-oceanic dispersal in an austral mygalomorph spider. *PLOS ONE*, *12*(8), e0180139. doi: 10.1371/journal.pone.0180139

- Too restricted inside the Indian Ocean region (includes less than five of the nine delimited areas)

Jønsson, K. A., Borregaard, M. K., Carstensen, D. W., Hansen, L. A., Kennedy, J. D., Machac, A., … Rahbek, C. (2017). Biogeography and Biotic Assembly of Indo-Pacific Corvoid Passerine Birds. *Annual Review of Ecology, Evolution, and Systematics*, *48*(1), 231–253. doi: 10.1146/annurev-ecolsys-110316-022813

- Too restricted inside the Indian Ocean region (includes less than five of the nine delimited areas)

Kuntner, M., & Agnarsson, I. (2011). Phylogeography of a successful aerial disperser: the golden orb spider *Nephila* on Indian Ocean islands. *BMC Evolutionary Biology*, *11*(1), 119. doi: 10.1186/1471-2148-11-119

- Too restricted inside the Indian Ocean region (includes less than five of the nine delimited areas)

Lo, E. Y., Duke, N. C., & Sun, M. (2014). Phylogeographic pattern of *Rhizophora* (Rhizophoraceae) reveals the importance of both vicariance and long-distance oceanic dispersal to modern mangrove distribution. *BMC Evolutionary Biology*, *14*(1), 83. doi: 10.1186/1471-2148-14-83

- Global biogeography with limited geographic and phylogenetic coverage of the Indian Ocean region

Mast, A. R., Willis, C. L., Jones, E. H., Downs, K. M., & Weston, P. H. (2008). A smaller Macadamia from a more vagile tribe: inference of phylogenetic relationships, divergence times, and diaspore evolution in Macadamia and relatives (tribe Macadamieae; Proteaceae). *American Journal of Botany*, *95*(7), 843–870. doi: 10.3732/ajb.0700006

- Global biogeography with limited geographic and phylogenetic coverage of the Indian Ocean region

Matos-Maraví, P., Matzke, N. J., Larabee, F. J., Clouse, R. M., Wheeler, W. C., Sorger, D. M., … Janda, M. (2018). Taxon cycle predictions supported by model-based inference in Indo-Pacific trap-jaw ants (Hymenoptera: Formicidae: *Odontomachus*). *Molecular Ecology*, *27*(20), 4090–4107. doi: 10.1111/mec.14835

- Too restricted inside the Indian Ocean region (includes less than five of the nine delimited areas)

Müller, C. J., Wahlberg, N., & Beheregaray, L. B. (2010). 'After Africa': the evolutionary history and systematics of the genus *Charaxes* Ochsenheimer (Lepidoptera: Nymphalidae) in the Indo-Pacific region. *Biological Journal of the Linnean Society*, *100*(2), 457–481. doi: 10.1111/j.1095-8312.2010.01426.x

- Too restricted inside the Indian Ocean region (includes less than five of the nine delimited areas)

Le Péchon, T., Zhang, L., He, H., Zhou, X.-M., Bytebier, B., Gao, X.-F., & Zhang, L.-B. (2016). A well-sampled phylogenetic analysis of the polystichoid ferns (Dryopteridaceae) suggests a complex biogeographical history involving both boreotropical migrations and recent transoceanic dispersals. *Molecular Phylogenetics and Evolution*, *98*, 324–336. doi: 10.1016/j.ympev.2016.02.018

- Global biogeography with limited geographic and phylogenetic coverage of the Indian Ocean region

Renner, S. S. (2004). Bayesian analysis of combined chloroplast loci, using multiple calibrations, supports the recent arrival of Melastomataceae in Africa and Madagascar. *American Journal of Botany*, *91*(9), 1427–1435. doi: 10.3732/ajb.91.9.1427

- Too restricted inside the Indian Ocean region (includes less than five of the nine delimited areas)

Sharma, P. P., Santiago, M. A., Kriebel, R., Lipps, S. M., Buenavente, P. A. C., Diesmos, A. C., … Wheeler, W. C. (2017). A multilocus phylogeny of Podoctidae (Arachnida, Opiliones, Laniatores) and parametric shape analysis reveal the disutility of subfamilial nomenclature in armored harvestman systematics. *Molecular Phylogenetics and Evolution*, *106*, 164–173. doi: 10.1016/j.ympev.2016.09.019

- Too restricted inside the Indian Ocean region (includes less than five of the nine delimited areas)

Simonsen, T. J., Zakharov, E. V., Djernaes, M., Cotton, A. M., Vane-Wright, R. I., & Sperling, F. A. H. (2011). Phylogenetics and divergence times of Papilioninae (Lepidoptera) with special reference to the enigmatic genera *Teinopalpus* and *Meandrusa*. *Cladistics*, *27*(2), 113–137. doi: 10.1111/j.1096-0031.2010.00326.x

- Global biogeography with limited geographic and phylogenetic coverage of the Indian Ocean region

Strandberg, J., & Johanson, K. A. (2011). The historical biogeography of *Apsilochorema* (Trichoptera, Hydrobiosidae) revised, following molecular studies. *Journal of Zoological Systematics and Evolutionary Research*, *49*(2), 110–118. doi: 10.1111/j.1439-0469.2010.00578.x

- Too restricted inside the Indian Ocean region (includes less than five of the nine delimited areas)

Strijk, J. S., Bone, R. E., Thébaud, C., Buerki, S., Fritsch, P. W., Hodkinson, T. R., & Strasberg, D. (2014). Timing and tempo of evolutionary diversification in a biodiversity hotspot: Primulaceae on Indian Ocean islands. *Journal of Biogeography*, *41*(4), 810–822. doi: 10.1111/jbi.12259

- Too restricted inside the Indian Ocean region (includes less than five of the nine delimited areas)

Strümpher, W. P., Sole, C. L., Villet, M. H., & Scholtz, C. H. (2014). Phylogeny of the family Trogidae (Coleoptera: Scarabaeoidea) inferred from mitochondrial and nuclear ribosomal DNA sequence data. *Systematic Entomology*, *39*(3), 548–562. doi: 10.1111/syen.12074

- Global biogeography with limited geographic and phylogenetic coverage of the Indian Ocean region

Sun, Y., He, X., & Glenny, D. (2014). Transantarctic disjunctions in Schistochilaceae (Marchantiophyta) explained by early extinction events, post-Gondwanan radiations and palaeoclimatic changes. *Molecular Phylogenetics and Evolution*, *76*, 189–201. doi: 10.1016/j.ympev.2014.03.018

- Global biogeography with limited geographic and phylogenetic coverage of the Indian Ocean region

Thomas, N., Bruhl, J. J., Ford, A., & Weston, P. H. (2014). Molecular dating of Winteraceae reveals a complex biogeographical history involving both ancient Gondwanan vicariance and long-distance dispersal. *Journal of Biogeography*, *41*(5), 894–904. doi: 10.1111/jbi.12265

- Too restricted inside the Indian Ocean region (includes less than five of the nine delimited areas)

Toussaint, E. F. A., & Short, A. E. Z. (2018). Transoceanic Stepping–stones between Cretaceous waterfalls? The enigmatic biogeography of pantropical *Oocyclus* cascade beetles. *Molecular Phylogenetics and Evolution*, *127*, 416–428. doi: 10.1016/j.ympev.2018.04.023

- Global biogeography with limited geographic and phylogenetic coverage of the Indian Ocean region

Wood, H. M., Gillespie, R. G., Griswold, C. E., & Wainwright, P. C. (2015). Why is Madagascar special? The extraordinarily slow evolution of pelican spiders (Araneae, Archaeidae). *Evolution*, *69*(2), 462–481. doi: 10.1111/evo.12578

- Too restricted inside the Indian Ocean region (includes less than five of the nine delimited areas)

Zhang, L., & Zhang, L.-B. (2017). A Classification of the Fern Genus *Tectaria* (Tectariaceae: Polypodiales) Based on Molecular and Morphological Evidence1. *Annals of the Missouri Botanical Garden*, *103*(2), 188–199. doi: 10.3417/2017007
Incomplete biogeographic information and lack of a calibration method to date the phylogenies
