## Supplementary material S0 to S4 for "Ecological changes have driven biotic exchanges across the Indian Ocean": S2.pdf

### Appendix 2: Specifics of the analyses

Please note that the analyses that followed the exact methodology described in the main text with the parameters supplied by the original authors list only the taxa available. Those analyses where we had to modify or add calibration points, or change priors to obtain chain convergence, have a short text explaining our modifications to allow replication of our results.

#### 1. Acrocephalidae

Table S1: Genbank numbers for the sequences used in the Acrocephalidae analyses with the distribution attributed to each terminal.

| Species | Locality | CytB | COI | ND2 | LDH | ODC | Myo | FIB5 | RAG1 |
|---|---|---|---|---|---|---|---|---|---|
| *Acrocephalus aedon* | Asia | FJ883020 AJ004778 | HQ608890 | DQ125987 | FJ883054 | FJ883126 | FJ883104 | | |
| *Acrocephalus aequinoctialis* | Pacific | EF156278 | | EF156314 | | | | | |
| *Acrocephalus agricola* | Asia | FJ883021 AJ004775 | KJ453121 | KJ453197 | FJ883055 | FJ883127 | FJ883108 | KJ453263 | KJ453456 |
| *Acrocephalus arundinaceus* | Africa Asia Europe | FJ883022 AJ004784 | FR847226 | AB621334 | FJ883056 | FJ883128 | FJ883098 | KJ453264 | KJ453460 |
| *Acrocephalus atyphus* | Pacific | EF156297 | | HQ844387 | | | | | |
| *Acrocephalus australis* | Australia | FJ883023 AJ004786 | KJ453124 | KJ453200 | FJ883057 | FJ883129 | FJ883097 | KJ453267 | KJ453459 |
| *Acrocephalus baeticatus* | Africa | KF547904 | KJ453126 | KJ453201 | FJ883058 | FJ883130 | FJ883103 | KJ453268 | KJ453462 |
| *Acrocephalus bistrigiceps* | Asia | FJ883025 AJ004766 | KJ453127 | KJ453203 | FJ883059 | FJ883131 | FJ883109 | KJ453270 | KJ453464 |
| *Acrocephalus brevipennis* | Africa | FJ883026 | KJ453129 | KJ453205 | FJ883060 | FJ883132 | FJ883099 | KJ453273 | KJ453467 |
| *Acrocephalus caffer* | Pacific | EF156308 | | EF156348 | | | | | |
| *Acrocephalus concinens* | Asia | FJ883027 | KJ453131 | KJ453206 | FJ883061 | FJ883133 | FJ883101 | KJ453274 | KJ453468 |
| *Acrocephalus dumetorum* | Asia Europe India | FJ883028 AJ004773 | KJ453133 | KJ453208 | FJ883062 | FJ883134 | FJ883105 | EF626749 | FJ358146 |
| *Acrocephalus familiaris* | Pacific | EU119965 | | EU119962 | | | | | |
| *Acrocephalus gracilirostris* | Africa | FJ883029 AJ004782 | KJ453135 | KJ453210 | FJ883063 | FJ883135 | FJ883095 | KJ453277 | KJ453470 |
| *Acrocephalus griseldis* | Africa | FJ883030 AJ004779 | KJ453138 | KJ453212 | FJ883064 | FJ883136 | FJ883092 | KJ453279 | KJ453472 |
| *Acrocephalus kerearako* | Pacific | EF156292 | | HQ844403 | | | | | |
| *Acrocephalus melanopogon* | Europe Asia | FJ883032 AJ004767 | KJ453139 | KJ453214 | FJ883066 | FJ883138 | FJ883093 | KJ453281 | KJ453474 |
| *Acrocephalus mendanae* | Pacific | EF156310 | | EF156322 | | | | | |
| *Acrocephalus newtoni* | Madagascar | FJ883033 AJ004780 | KJ453141 | KJ453216 | FJ883067 | FJ883139 | FJ883123 | KJ453283 | AY319972 |
| *Acrocephalus orientalis* | Asia | FJ883034 AJ004785 | JF499074 | KJ453218 | FJ883068 | FJ883140 | FJ883102 | KJ453285 | KJ453477 |
| *Acrocephalus orinus* | Asia | DQ681065 | | | | | | | |
| *Acrocephalus paludicola* | Africa Asia Europe | FJ883035 AJ004768 | KJ453144 | KJ453220 | FJ883069 | FJ883141 | FJ883112 | KJ453287 | KJ453479 |
| *Acrocephalus palustris* | Africa Asia Europe | FJ883036 AJ004774 | KJ453146 | KJ453222 | FJ883070 | FJ883142 | FJ883106 | KJ453289 | KJ453481 |
| *Acrocephalus percernis* | Pacific | EF156298 | | HQ844418 | | | | | |
| *Acrocephalus rimitarae* | Pacific | EF156306 | | EF156346 | | | | | |
| *Acrocephalus rufescens* | Africa | FJ883037 Z73475 | KJ453148 | KJ453224 | FJ883071 | FJ883143 | FJ883124 | KJ453291 | KJ453484 |
| *Acrocephalus schoenobaenus* | Asia | FJ883038 | KJ453150 | KJ453226 | FJ883072 | FJ883144 | FJ883107 | KJ453293 | KJ453486 |
| *Acrocephalus scirpaceus fuscus* | Asia Africa | KF547898 | KJ453154 | KJ453230 | KJ453349 | KJ453433 | KJ453390 | KJ453300 | KJ453488 |
| *Acrocephalus scirpaceus scirpaceus* | Europe Africa | KF547878 Z73483 | KJ453152 | KJ453229 | FJ883073 | FJ883145 | FJ883111 | KJ453302 | KJ453491 |

| | | | | | | | | | | |
|---|---|---|---|---|---|---|---|---|---|---|
| *Acrocephalus sechellensis* | Seychelles | FJ883040 | AJ004781 | KJ453156 | KJ453232 | FJ883074 | FJ883146 | FJ883122 | KJ453295 | KJ453492 |
| *Acrocephalus stentoreus* | Arabic Peninsula | FJ883031 | AJ004787 | FJ465293 | KJ453233 | FJ883065 | FJ883137 | FJ883100 | KJ453296 | KJ453495 |
| *Acrocephalus taiti* | Pacific | FJ883042 | AJ004777 | KJ453161 | HQ844427 | FJ883076 | FJ883148 | FJ883096 | | |
| *Acrocephalus tangorum* | Asia IAA | FJ883041 | | KJ453158 | KJ453235 | FJ883075 | FJ883147 | FJ883110 | KJ453305 | KJ453497 |
| *Acrocephalus vaughani* | Pacific | KJ453185 | | KJ453161 | KJ453237 | KJ453354 | KJ453440 | KJ453394 | KJ453299 | KJ453493 |
| *Bradypterus baboecala* | Africa | FJ883053 | | | AY382344 | FJ883090 | FJ883162 | DQ008525 | | JX236420 |
| *Calamonastides gracilirostris* | Africa | FJ883043 | | KJ453162 | KJ453239 | FJ883077 | FJ883149 | FJ883113 | KJ453306 | KJ453499 |
| *Hippolais icterina* | Europe Africa | FJ883046 | DQ008479 | KJ453164 | GQ242091 | FJ883078 | FJ883153 | FJ883120 | EU680651 | KJ453516 |
| *Hippolais languida* | Africa Arabic Peninsula Asia | FJ883047 | AJ004794 | KJ453166 | KJ453242 | FJ883079 | FJ883154 | FJ883094 | KJ453308 | KJ453502 |
| *Hippolais olivetorum* | Africa | FJ883048 | AJ004795 | JQ175054 | KJ453243 | FJ883080 | FJ883155 | FJ883121 | KJ453310 | KJ453503 |
| *Hippolais polyglotta* | Africa Europe | FJ883050 | AJ004797 | KJ453169 | KJ453245 | FJ883081 | FJ883157 | FJ883115 | KJ453312 | KJ453505 |
| *Iduna caligata* | Asia India | FJ883044 | AJ004793 | GQ481975 | KJ453250 | FJ883084 | FJ883151 | FJ883116 | KJ453317 | KJ453512 |
| *Iduna natalensis* | Africa | DQ008523 | | KJ453171 | KJ453252 | FJ883082 | FJ883150 | FJ883114 | KJ453319 | KJ453519 |
| *Iduna opaca* | Africa | FJ883049 | AJ004790 | KJ453173 | KJ453253 | FJ883086 | FJ883156 | FJ883118 | KJ453320 | KJ453513 |
| *Iduna pallida* | Africa Asia | FJ883045 | AJ004791 | KJ453175 | KJ453254 | FJ883085 | FJ883152 | FJ883119 | KJ453321 | KJ453509 |
| *Iduna rama* | Asia | FJ883051 | AJ004792 | KJ453176 | KJ453256 | FJ883087 | FJ883158 | FJ883117 | KJ453323 | KJ453514 |
| *Iduna similis* | Africa | FJ899738 | | KJ453178 | KJ453258 | FJ883083 | FJ883159 | FJ883125 | KJ453326 | KJ453520 |
| *Locustella fluviatilis* | Africa Arabic Peninsula Asia Europe | HQ608847 | | JQ175275 | GQ242093 | HQ706203 | GQ242152 | DQ008527 | GQ242048 | KJ453524 |
| *Locustella lanceolata* | Asia IAA | DQ119525 | | GU571953 | AY382379 | FJ883088 | FJ883160 | FJ883091 | | |
| *Megalurus palustris* | IAA India | FJ883052 | | JF957022 | JN614731 | FJ883089 | FJ883161 | DQ008529 | EU680661 | AY319988 |
| *Nesilla typica* | Madagascar | KJ453193 | | KJ453180 | KJ453260 | KJ453371 | KJ453455 | KJ453412 | EU680665 | KJ453522 |

For the analyses for the Acrocephalidae, we complemented some of the Cytochrome B sequences (see Table S1). We made two independent runs (see main text) and combined their results through LogCombiner (Drummond & Rambaut, 2007). Molecular rates were obtained from the original article.

#### 2. Anatini

Table S2: Genbank numbers for the sequences used in the Acrocephalidae analyses with the distribution attributed to each terminal.

| Species | Locality | CytB | 12S | COI | ND2 | ATP6 |
|---|---|---|---|---|---|---|
| *Aix sponsa* | America | AF059053 | | AY666569 | AF059114 | |
| *Amazonetta brasiliensis* | America | AF059054 | HM063533 | JN801484 | AF059115 | |
| *Anas acuta* | Africa America Asia India | AF059055 | HM063538 | KBBI227-07 | AF059116 | |
| *Anas americana* | America | AF059103 | HM063537 | DQ433309 | AF059163 | |
| *Anas aucklandica* | New Zealand | AF059056 | AF173482 | | AF059117 | AF173490 |
| *Anas bahamensis bahamensis* | America | AF059058 | | FJ027081 | AF059119 | |
| *Anas bernieri* | Madagascar | AF059060 | | | AF059121 | |
| *Anas capensis* | Africa | AF059105 | | | AF059165 | |
| *Anas carolinensis* | America | AF059063 | HM063539 | DQ434281 | AF059123 | |
| *Anas castanea* | Australia | AF059065 | AF173481 | | AF059125 | AF173494 |

| | | | | | | |
|---|---|---|---|---|---|---|
| *Anas chlorotis* | New Zealand | AF059061 | AF173484 | | AF059122 | AF173491 |
| *Anas clypeata* | Africa America Asia India | AF059062 | HM063536 | BROMB439-06 | AF059174 | |
| *Anas crecca* | Africa Asia India | AF059064 | HM063540 | GU571238 | AF059124 | |
| *Anas cyanoptera cyanoptera* | America | AF059066 | AF173690 | BROMB795-07 | AF059126 | |
| *Anas diazi* | America | AF059069 | | | AF059129 | |
| *Anas discors* | America | AF059068 | JF914878 | AY666325 | AF059128 | |
| *Anas erythrorhyncha* | Africa | AF059070 | | | AF059130 | |
| *Anas falcata* | Asia | AF059106 | AY164527 | GQ481322 | AF059166 | |
| *Anas flavirostris flavirostris* | America | AF059071 | | JN801487 | AF059131 | |
| *Anas formosa* | Asia | NC_015482 | NC_015482 | NC_015482 | NC_015482 | NC_015482 |
| *Anas fulvigula* | America | AF059074 | | DQ432723 | AF059134 | |
| *Anas georgica spinicauda* | America | AF059075 | | FJ027093 | AF059135 | |
| *Anas gracilis* | Australia | AF059076 | AF173480 | | AF059136 | AF173493 |
| *Anas hottentota* | Africa Madagascar | AF059077 | AF173692 | | AF059137 | |
| *Anas laysanensis* | Pacific | AF059078 | | JF498830 | AF059138 | |
| *Anas luzonica* | IAA | AF059079 | | | AF059139 | |
| *Anas melleri* | Madagascar | AF059080 | | | AF059140 | |
| *Anas nesiotis* | New Zealand | AF059057 | AF173483 | | AF059118 | AF173489 |
| *Anas penelope* | Africa Asia | AF059107 | AY164518 | GU571239 | AF059167 | |
| *Anas platalea* | America | AF059084 | | FJ027098 | AF059144 | |
| *Anas platyrhynchos* | Cosmopolite | NC_009684 | NC_009684 | NC_009684 | NC_009684 | NC_009684 |
| *Anas poecilorhyncha* | Asia India | AF059083 | AY164517 | JF499090 | AF059143 | |
| *Anas puna* | America | AF059085 | | FJ027101 | AF059145 | |
| *Anas querquedula* | Africa Asia India | AF059086 | AF173691 | GQ481326 | AF059146 | |
| *Anas rhynchotis rhynchotis* | Australia | AF059087 | | BROMB529-07 | AF059147 | |
| *Anas rubripes* | America | AF059088 | | AY666211 | AF059148 | |
| *Anas sibilatrix* | America | AF059108 | | FJ027106 | AF059168 | |
| *Anas smithii* | Africa | AF059089 | | | AF059149 | |
| *Anas sparsa* | Africa | AF059091 | | | AF059151 | |
| *Anas strepera* | America | AF059109 | AF173689 | GQ481327 | AF059169 | |
| *Anas superciliosa* | Australia IAA Pacific | AF059092 | AF173486 | JN801396 | AF059152 | AF173488 |
| *Anas undulata* | Africa | AF059093 | | | AF059153 | |
| *Anas versicolor* | America | AF059094 | | FJ027115 | AF059154 | |
| *Anas zonorhyncha* | Asia IAA | AF059095 | | | AF059155 | |
| *Asarcornis scutulata* | IAA | AF059099 | HM063548 | | AF059159 | |
| *Aythya americana* | America | NC_000877 | NC_000877 | NC_000877 | NC_000877 | NC_000877 |
| *Cairina moschata* | America | NC_010965 | NC_010965 | NC_010965 | NC_010965 | NC_010965 |
| *Callonetta leucophrys* | America | AF059097 | HM063546 | FJ027277 | AF059157 | |
| *Chenonetta jubata* | Australia | AF059100 | HM063545 | BROMB794-07 | AF059160 | |
| *Cyanochen cyanopterus* | Africa | AF059101 | HM063550 | | AF059161 | |
| *Lophonetta specularioides* | America | AF059102 | HM063535 | | AF059162 | |
| *Marmaronetta angustirostris* | Mediterranean | AF059104 | HM063551 | | AF059164 | |
| *Pachyanas (Anas) chathamica* | New Zealand | KF562761 | KF562761 | KF562761 | KF562761 | KF562761 |

| | | | | | |
|---|---|---|---|---|---|
| *Pteronetta hartlaubii* | Africa | AF059110 | HM063549 | | AF059170 |
| *Sarkidiornis melanotos* | Africa IAA India Madagascar | AF059111 | HM063541 | FJ028237 | AF059171 |
| *Speculanas specularis* | America | AF059090 | HM063534 | | AF059150 |
| *Tachyeres pteneres* | America | AF059112 | HM063532 | JN802006 | AF059172 |
| *Tadorna tadorna* | Asia | AF059113 | HM063544 | GU571650 | AF059173 |

The vouchers starting with BROMB are deposited in the Barcode of Life database.

For these analyses, we partitioned each position of the codon separately for all markers, except 12S. A prior to enforce the monophyly of the ingroup and the outgroup (*Aix*, *Cairina* and *Tadorna*) was also included. For the *A. soporata* calibration we placed the mean in real space instead of log space; this means that the mean age is the actual μ of a normal distribution and not the M of the lognormal distribution (the original article seems to have done that although it is not explicit in the Materials and Methods section). We made two independent runs and combined their results through LogCombiner.

#### 3. Palaeognathae

Table S3: Genbank numbers for the sequences used in the Palaeognathae analyses with the distribution attributed to each terminal.

| Species | | Mitochondrion genome | BACH1 | BMP2 | CMOS | DNAH3 | IRBP | NT3 | PNN | PTPN | RAG2 | TRAF6 |
|---|---|---|---|---|---|---|---|---|---|---|---|---|
| *Acanthisitta chloris* | | AY325307 | JX533012 | JX533043 | JX533074 | JX533122 | JX533214 | JX533265 | JX533296 | JX533327 | JX533380 | JX533411 |
| *Aepyornis hildebrandti* | Madagascar | KJ749824 | | | | | | | | | | |
| *Alectura lathami* | | NC_007227 | | | | | | | | | | |
| *Alligator mississippiensis* | | NC_001922 | JX533010 | JX533041 | JX533072 | JX533120 | JX533212 | JX533263 | JX533294 | JX533325 | JX533378 | JX533409 |
| *Anas platyrhynchos* | | NC_009684 | | | | | | | | | | |
| *Anomalopteryx didiformis* | New Zealand | NC_002779 | JX533038 | JX533069 | JX533100 | JX533148 | JX533240 | JX533291 | JX533322 | JX533353 | JX533406 | JX533437 |
| *Anser albifrons* | | NC_004539 | JX533023 | JX533054 | JX533085 | JX533133 | JX533225 | JX533276 | JX533307 | JX533338 | JX533391 | JX533422 |
| *Anseranas semipalmata* | | NC_005933 | JX533024 | JX533055 | JX533086 | JX533134 | JX533226 | JX533277 | JX533308 | JX533339 | JX533392 | JX533423 |
| *Apteryx haastii* | New Zealand | NC_002782 | | | | | | | | | | |
| *Apteryx mantelli* | New Zealand | AY016010 | JX533033 | JX533064 | JX533095 | JX533143 | JX533235 | JX533286 | JX533317 | JX533348 | JX533401 | JX533432 |
| *Apteryx owenii* | New Zealand | NC_013806 | JX533031 | JX533062 | JX533093 | JX533141 | JX533233 | JX533284 | JX533315 | JX533346 | JX533399 | JX533430 |
| *Apus apus* | | NC_008540 | JX533021 | JX533052 | JX533083 | JX533131 | JX533223 | JX533274 | JX533305 | JX533336 | JX533389 | JX533420 |
| *Archilochus colubris* | | NC_010094 | JX533022 | JX533053 | JX533084 | JX533132 | JX533224 | JX533275 | JX533306 | JX533337 | JX533390 | JX533421 |
| *Arenaria interpres* | | NC_003712 | JX533019 | JX533050 | JX533081 | JX533129 | JX533221 | JX533272 | JX533303 | JX533334 | JX533387 | JX533418 |
| *Caiman crocodilus* | | NC_002744 | JX533011 | JX533042 | JX533073 | JX533121 | JX533213 | JX533264 | JX533295 | JX533326 | JX533379 | JX533410 |
| *Casuarius bennetti* | Australia IAA | AY016011 | | | | | | | | | | |
| *Casuarius casuarius* | Australia | AF338713 | JX533036 | JX533067 | JX533098 | JX533146 | JX533238 | JX533289 | JX533320 | JX533351 | JX533404 | JX533435 |
| *Ciconia ciconia* | | NC_002197 | JX533016 | JX533047 | JX533078 | JX533126 | JX533218 | JX533269 | JX533300 | JX533331 | JX533384 | JX533415 |
| *Crypturellus tataupa* | America | AY016012 | | | | | | | | | | |
| *Dinornis giganteus* | New Zealand | NC_002672 | | | | | | | | | | |
| *Dromaius novaehollandiae* | Australia | AF338711 | JX533037 | JX533068 | JX533099 | JX533147 | JX533239 | JX533290 | JX533321 | JX533352 | JX533405 | JX533436 |
| *Emeus crassus* | New Zealand | NC_002673 | | | | | | | | | | |
| *Eudromia elegans* | America | NC_002772 | JX533025 | JX533056 | JX533087 | JX533135 | JX533227 | JX533278 | JX533309 | JX533340 | JX533393 | JX533424 |

| | | | | | | | | | | | | |
|---|---|---|---|---|---|---|---|---|---|---|---|---|
| *Gallus gallus* | | NC_001323 | | | | | | | | | | |
| *Gavia stellata* | | NC_007007 | JX533013 | JX533044 | JX533075 | JX533123 | JX533215 | JX533266 | JX533297 | JX533328 | JX533381 | JX533412 |
| *Haematopus ater* | | NC_003713 | JX533018 | JX533049 | JX533080 | JX533128 | JX533220 | JX533271 | JX533302 | JX533333 | JX533386 | JX533417 |
| *Mullerornis agilis* | Madagascar | KJ749825 | | | | | | | | | | |
| *Pterocnemia pennata* | America | NC_002783 | JX533030 | JX533061 | JX533092 | JX533140 | JX533232 | JX533283 | JX533314 | JX533345 | JX533398 | JX533429 |
| *Pterodroma brevirostris* | | NC_007174 | JX533017 | JX533048 | JX533079 | JX533127 | JX533219 | JX533270 | JX533301 | JX533332 | JX533385 | JX533416 |
| *Pygoscelis adeliae* | | NC_021137 | JX533015 | JX533046 | JX533077 | JX533125 | JX533217 | JX533268 | JX533299 | JX533330 | JX533383 | JX533414 |
| *Rhea americana* | America | NC_000846 | JX533029 | JX533060 | JX533091 | JX533139 | JX533231 | JX533282 | JX533313 | JX533344 | JX533397 | JX533428 |
| *Struthio camelus* | Africa | NC_002785 | JX533028 | JX533059 | JX533090 | JX533138 | JX533230 | JX533281 | JX533312 | JX533343 | JX533396 | JX533427 |
| *Taeniopygia guttata* | | NC_007897 | | | | | | | | | | |
| *Tinamus major* | America | AF338707 | JX533026 | JX533057 | JX533088 | JX533136 | JX533228 | JX533279 | JX533310 | JX533341 | JX533394 | JX533425 |

In this analysis, the complete dataset and the mitogenome had a difference in topology: the mitogenome datset retrieved the same groups as the original article, but with all markers the rheas were retrieved as a sister-group to the *Aepyornis-Apteryx-Casuarius-Dromaius* clade. These two topologies, however, did not interfere with the obtained dates or with the RASP results and our further discussion.

#### 4. Psittacoidea

Table S4: Genbank numbers for the sequences used in the Psittacoidea analyses with the distribution attributed to each terminal.

| Species | Locality | CytB | CMOS | RAG1 | ZENK |
|---|---|---|---|---|---|
| *Ara macao* | America | MH400248 | JF807951 | JF807979 | JF807965 |
| *Agapornis canus* | Madagascar | AF001332 | GQ505083 | GQ505191 | GQ505138 |
| *Agapornis fischeri* | Africa | AF346358 | GQ505084 | GQ505192 | GQ505139 |
| *Agapornis lilianae* | Africa | AF001331 | GQ505087 | | |
| *Agapornis nigrigenis* | Africa | AF001328 | GQ505085 | GQ505193 | GQ505140 |
| *Agapornis roseicollis* | Africa | AF001330 | GQ505086 | GQ505194 | GQ505141 |
| *Alisterus chloropterus* | Australia IAA | | GQ505091 | GQ505199 | GQ505145 |
| *Alisterus scapularis* | Australia | FJ499014 | GQ505090 | GQ505198 | GQ505144 |
| *Aprosmictus jonquillaceus* | Australia | AB177959 | GQ505092 | GQ505200 | GQ505146 |
| *Amazona aestiva* | America | NC_033336 | JF807952 | JF807980 | JF807966 |
| *Amazona dufresniana* | America | AY283454 | JF807953 | JF807981 | JF807967 |
| *Amazona pretrei* | America | | JF807954 | JF807982 | JF807968 |
| *Barnardius zonarius* | Australia | JQ066207 | GQ505095 | GQ505203 | GQ505149 |
| *Bolbopsittacus lunulatus* | IAA | | JF807955 | JF807983 | JF807969 |
| *Cacatua galerita fitzroyi* | Australia IAA | AF313755 | GQ505120 | GQ505231 | GQ505173 |
| *Cacatua moluccensis* | IAA | AB177980 | GQ505121 | GQ505232 | GQ505174 |
| *Calyptorhynchus funereus* | Australia | JF414306 | GQ505118 | GQ505229 | |
| *Calyptorhynchus latirostris* | Australia | JF414302 | GQ505119 | GQ505230 | GQ505172 |
| *Charmosyna pulchella* | Australia IAA | JX442429 | GQ505126 | GQ505237 | GQ505179 |
| *Coracopsis nigra* | Comoros Madagascar Seychelles | GQ996494 | GQ505114 | GQ505224 | |

| | | | | | |
|---|---|---|---|---|---|
| *Coracopsis vasa* | Comoros Madagascar | AF346355 | GQ505113 | GQ505223 | GQ505167 |
| *Cyanoramphus auriceps* | New Zealand | JX442418 | GQ505104 | GQ505213 | GQ505158 |
| *Cyanoramphus novaezelandiae* | New Zealand | AF346380 | GQ505103 | GQ505212 | GQ505157 |
| *Cyclopsitta diophthalma* | Australia IAA | | GQ505130 | | |
| *Deroptyus accipitrinus* | America | DQ150992 | JF807956 | JF807984 | JF807970 |
| *Eclectus roratus* | Australia IAA | AB177965 | GQ505135 | GQ505244 | GQ505187 |
| *Eos cyanogenia* | IAA | AF346330 | GQ505122 | GQ505233 | GQ505175 |
| *Eunymphicus cornutus* | Pacific | AF242516 | GQ505106 | GQ505215 | GQ505159 |
| *Eunymphicus uvaeensis* | Pacific | AF242514 | GQ505107 | GQ505216 | GQ505160 |
| *Guarouba guarouba* | America | DQ150990 | JF807957 | JF807985 | JF807971 |
| *Lathamus discolor* | Australia | JX442416 | GQ505102 | GQ505211 | GQ505156 |
| *Loriculus catamene* | IAA | | GQ505088 | GQ505195 | GQ505142 |
| *Loriculus galgulus* | IAA | AB177967 | GQ505089 | GQ505196 | |
| *Loriculus philippensis* | IAA | | | GQ505197 | GQ505143 |
| *Lorius garrulus* | IAA | AB177951 | GQ505125 | GQ505236 | GQ505178 |
| *Mascarinus mascarinus* | Reunion | GQ996499 | | | |
| *Melopsittacus undulatus* | Australia | EF450826 | | GQ505222 | GQ505166 |
| *Micropsitta finschii tristrami* | Pacific | U89176 | GQ505128 | GQ505240 | GQ505182 |
| *Micropsitta pusio* | Pacific | | GQ505129 | GQ505241 | GQ505183 |
| *Neophema chrysogaster* | Australia | NC_019804 | GQ505110 | GQ505219 | GQ505163 |
| *Neophema chrysostoma* | Australia | JX442425 | GQ505111 | GQ505220 | GQ505164 |
| *Neophema pulchella* | Australia | JX442424 | GQ505109 | GQ505218 | GQ505162 |
| *Neophema splendida* | Australia | JQ066212 | GQ505108 | GQ505217 | GQ505161 |
| *Neopsephotos bourkii* | Australia | JQ066213 | GQ505112 | GQ505221 | GQ505165 |
| *Nestor notabilis* | New Zealand | KX369037 | JF807958 | GQ505238 | GQ505180 |
| *Northiella haematogaster* | Australia | JX442432 | JF807959 | JF807986 | JF807972 |
| *Pionus menstruus* | America | EF517605 | JF807960 | JF807987 | JF807973 |
| *Platycercus caledonicus* | Australia | JQ066208 | GQ505097 | GQ505205 | GQ505151 |
| *Platycercus eximius* | Australia | DQ467901 | | GQ505206 | GQ505152 |
| *Platycercus flaveolus* | Australia | JX442434 | GQ505099 | GQ505208 | |
| *Platycercus venustus* | Australia | JX442413 | GQ505098 | GQ505207 | GQ505153 |
| *Poicephalus gulielmi* | Africa | AY283498 | JF807961 | JF807988 | JF807974 |
| *Poicephalus meyeri* | Africa | GQ996511 | | JF807989 | JF807975 |
| *Poicephalus rufiventris* | Africa | MG736916 | GQ505116 | GQ505226 | GQ505169 |
| *Poicephalus senegalus* | Africa | AF346359 | | GQ505227 | GQ505170 |
| *Polytelis alexandrae* | Australia | JQ066206 | GQ505093 | GQ505201 | GQ505147 |
| *Polytelis anthopeplus* | Australia | AF346386 | GQ505094 | GQ505202 | GQ505148 |
| *Prioniturus discurus* | IAA | JQ066197 | GQ505133 | | GQ505186 |
| *Prioniturus luconensis* | IAA | NC_027846 | GQ505134 | | |
| *Prioniturus montanus* | IAA | JQ066185 | JF807962 | JF807990 | JF807976 |
| *Prosopeia tabuensis* | Pacific | EU627173 | GQ505105 | GQ505214 | |
| *Psephotus chrysopterygius* | Australia | JX442436 | JF807963 | JF807991 | JF807977 |
| *Psephotus dissimilis* | Australia | JX442415 | GQ505101 | GQ505210 | GQ505155 |

| | | | | | |
|---|---|---|---|---|---|
| *Psephotus varius* | Australia | JX442414 | GQ505100 | GQ505209 | GQ505154 |
| *Psittacella brehmii* | Australia IAA | JQ066220 | JF807964 | JF807992 | JF807978 |
| *Psittacula alexandri alexandri* | IAA | AB177970 | | | |
| *Psittacula alexandri fasciata* | Asia | GQ996507 | | | |
| *Psittacula calthrapae* | India | GQ996512 | | | |
| *Psittacula columboides* | India | AY220108 | | | |
| *Psittacula cyanocephala* | India | GQ996508 | | KJ456123 | |
| *Psittacula derbiana* | Asia | AF346388 | U88424 | | |
| *Psittacula echo* | Mauritius | AY220113 | | | |
| *Psittacula eques* | Reunion | LN614517 | | | |
| *Psittacula eupatria* | Asia / India / IAA | AY220115 | GQ505137 | GQ505246 | GQ505189 |
| *Psittacula exsul* | Rodrigues | LN614516 | | | |
| *Psittacula finschii* | Asia | GQ996510 | | | |
| *Psittacula himalayana* | Asia | AY220102 | | KJ456124 | |
| *Psittacula krameri borealis* | Asia | AY220116 | | | |
| *Psittacula krameri krameri* | Africa | AY220117 | | | |
| *Psittacula krameri manillensis* | India | GQ996517 | | | |
| *Psittacula krameri parvirostris* | Africa | GQ996497 | | | |
| *Psittacula longicauda* | IAA | GQ996509 | | | |
| *Psittacula roseata* | India | AY220107 | | KJ456126 | |
| *Psittacula wardi* | Seychelles | GQ996500 | | | |
| *Psittaculirostris desmarestii* | IAA | AB177960 | GQ505131 | GQ505242 | GQ505184 |
| *Psittaculirostris edwardsii* | Australia IAA | AB177971 | GQ505132 | GQ505243 | GQ505185 |
| *Psittacus erithacus erithacus* | Africa | AY082082 | GQ505115 | GQ505225 | GQ505168 |
| *Psitteuteles goldiei* | Australia IAA | KM372512 | GQ505124 | GQ505235 | GQ505177 |
| *Psittinus cyanurus* | IAA | JQ066222 | | GQ505247 | GQ505190 |
| *Psittrichas fulgidus* | IAA | AF346391 | GQ505127 | GQ505239 | GQ505181 |
| *Purpureicephalus spurius* | Australia | JX442410 | GQ505096 | GQ505204 | GQ505150 |
| *Tanygnathus megalorhynchus* | IAA | GQ996516 | GQ505136 | GQ505245 | GQ505188 |
| *Trichoglossus johnstoniae* | IAA | | GQ505123 | GQ505234 | GQ505176 |
| *Triclaria malachitacea* | America | AY669442 | GQ505117 | GQ505228 | GQ505171 |
| *Anas* | | KX534428 | AF478185 | AF143729 | EU738887 |
| *Anser* | | EU585613 | AY994065 | DQ137227 | EU738899 |
| *Larus* | | AB208758 | U88419 | AY228799 | EU738965 |
| *Coracias caudata* | | U89184 | AY056916 | AF143737 | EU738931 |
| *Todus angustirostris* | | AF441626 | AF441636 | DQ111790 | EU738886 |
| *Gavia* | | HQ864537 | U88423 | DQ137228 | EU738953 |
| *Gallus* | | EU839454 | AY056925 | AF143730 | EU738891 |
| *Falco* | | KP863030 | AY447974 | AY461399 | AF490155 |
| *Acanthisitta chloris* | | AY325307 | AY056903 | AY056975 | EU738893 |
| *Corvus corone* | | JQ864491 | AY056918 | AY056989 | EF568306 |
| *Picathartes gymnocephalus* | | KJ909200 | AY056950 | AY057019 | EF568314 |
| *Pipra coronata* | | FJ899382 | AY056951 | AY057020 | AF492518 |

| | | | | |
|---|---|---|---|---|
| *Pitta* | KJ456408 | AY056952 | AY057021 | EF568299 |
| *Scopus umbretta* | KX534433 | AF339323 | DQ881830 | EU739024 |
| *Pelecanus* | MH041272 | AF339325 | DQ881819 | EU738992 |
| *Phalacrocorax carbo* | NC_027267 | AF339332 | DQ881821 | EU738997 |
| *Phoenicopterus* | KJ400319 | AF339336 | DQ881823 | EU739001 |
| *Podiceps* | NC_008140 | AF339334 | DQ881825 | EU739009 |
| *Puffinus* | AJ004215 | U88421 | DQ881827 | AF490146 |
| *Eudyptes pachyrhynchus (Spheniscidiae)* | DQ137210 | U88420 | DQ137231 | |
| *Eudyptula minor (Spheniscidiae)* | MF370525 | | | EU738944 |
| *Struthio camelus* | NC_002785 | U88429 | AF143727 | EU738886 |

In this analysis, we added some terminal taxa and sequences. By checking multiple runs, we found out that the *Coracopsis* species were acting as wildcards, changing their position across the tree. Most of the time, this genus was a sister-clade of *Mascarinus*. We checked the literature and concluded that it was a good approximation to enforce the monophyly of *Coracopsis*-*Mascarinus*-*Psittrichas* clade.

#### 5. Pycnonotidae

Table S5: Genbank numbers for the sequences used in the Pycnonotidae analyses with the distribution attributed to each terminal.

| Species | Location | 12S | 16S | ATP6 | COI | CYTB | ND2 | ND3 | FIB-I5 | FIB-I7 | GAPDH-I11 | MB-I2 | ODC-I6/I7 | RAG1 | TGFB2-I5 |
|---|---|---|---|---|---|---|---|---|---|---|---|---|---|---|---|
| *Alophoixus bres* | IAA | AF386490 | AF391227 | KT312509 | | JN827010 | DQ402228 | DQ402289 | | DQ402319 | KT311996 | | KT312713 | | GU112563 |
| *Alophoixus finschii* | IAA | | | | | KY404180 | KY454707 | KY454726 | | | | KY454741 | | | KY454738 |
| *Alophoixus flaveolus* | Asia | AF386478 | AF391215 | KT312498 | JQ173973 | KJ456190 | KJ455318 | KT312363 | | | KT311974 | KJ454749 | KT312691 | KJ455972 | |
| *Alophoixus frater* | IAA | | | | | | GU112679 | GU112725 | | GU112633 | | | | | GU112587 |
| *Alophoixus ochraceus* | IAA | AF386482 | AF391219 | KT312419 | | KY404181 | DQ402229 | DQ402290 | | DQ402322 | KT311798 | | KT312551 | | GU112564 |
| *Alophoixus pallidus* | Asia | AF386471 | AF391209 | KT312428 | | DQ008507 | GQ242078 | GQ242112 | EF626743 | GU112600 | KT311828 | EF625281 | KT312565 | | GU112531 |
| *Alophoixus phaeocephalus* | IAA | AF386491 | AF391228 | KT312520 | | KY404182 | DQ402230 | DQ402291 | | DQ402323 | KT312018 | | KT312735 | | GU112565 |
| *Andropadus curvirostris* | Africa | | | | JQ174018 | | | | | | | | | | |
| *Andropadus gracilis* | Africa | | | | JQ174019 | | | | | | | | | | |
| *Andropadus importunus* | Africa | | | | FJ473230 | FJ487856 | AF003469 | GQ242115 | EF626713 | GQ242059 | | EF625252 | EF625302 | | |
| *Andropadus latirostris* | Africa | | | | JQ174020 | | | | | | | | | | |
| *Arizelocichla chlorigula* | Africa | | | | | AH005509 | AF003457 | | EF626703 | | | EF625243 | EF625292 | | |
| *Arizelocichla fusciceps* | Africa | | | | | AH005510 | AF003458 | | EF626700 | | | EF625240 | EF625289 | | |
| *Arizelocichla kakamegae* | Africa | | | | | AH005504 | KY454708 | KY454727 | | | | KY454742 | | | |
| *Arizelocichla kikuyensis* | Africa | | | | | AH005516 | AF003463 | | EF626701 | | | EF625241 | EF625290 | | |
| *Arizelocichla masukensis* | Africa | | | | | | GQ242085 | GQ242120 | EF626698 | GQ242064 | | JX236344 | EF625287 | JX236416 | |
| *Arizelocichla milanjensis* | Africa | | | | | | KY454709 | | | | | KY454743 | | | |
| *Arizelocichla montana* | Africa | | | | | AH005507 | AF003455 | | | | | | | | |
| *Arizelocichla neumanni* | Africa | | | | | | AF003456 | | EF626702 | | | EF625242 | EF625291 | | |
| *Arizelocichla nigriceps* | Africa | | | | | AH005511 | AF003459 | | | | | | | | |
| *Arizelocichla striifacies* | Africa | | | | | AH005513 | KY454710 | KY454728 | EF626704 | | | KY454744 | EF625293 | | |
| *Arizelocichla tephrolaema* | Africa | | | | | AF282785 | GQ242086 | GQ242121 | GQ242044 | GQ242065 | | GQ242109 | GQ242147 | | |
| *Atimastillas flavicollis* | Africa | | | | | JX236375 | DQ402205 | DQ402266 | EF626721 | DQ402318 | JX236287 | JX236346 | EF625310 | | |
| *Baeopogon clamans* | Africa | | | | | | DQ402203 | DQ402264 | EF626716 | | | EF625256 | EF625305 | | |
| *Baeopogon indicator* | Africa | AF386465 | AF391205 | | | | DQ402204 | DQ402265 | EF626717 | DQ402317 | | EF625255 | EF625306 | | |

| | | | | | | | | | | | | | | | |
|---|---|---|---|---|---|---|---|---|---|---|---|---|---|---|---|
| *Bernieria madagascariensis* | Madagascar | | | | | AF199387 | DQ402194 | DQ402255 | | DQ402338 | HQ333100 | HQ333071 | HQ333086 | JX236419 | |
| *Bleda canicapillus* | Africa | | | | | | DQ402201 | DQ402262 | GQ242043 | DQ402315 | | GQ242108 | GQ242146 | | GU112528 |
| *Bleda eximius* | Africa | | | | JQ174174 | | DQ402202 | DQ402263 | | DQ402316 | | | | | |
| *Bleda notatus* | Africa | AF386474 | AF391203 | | | | KY454711 | KY454729 | EF626739 | | | KY454745 | EF625328 | | |
| *Bleda syndactylus* | Africa | AF386466 | AF391204 | | JQ174177 | KC355099 | AY136592 | AY590757 | EF626738 | GQ242063 | | EF625276 | EF625327 | AY319976 | |
| *Calyptocichla serinus* | Africa | | | | | | DQ402199 | DQ402260 | EF626715 | DQ402314 | | EF625254 | EF625304 | | GU112566 |
| *Cerasophila thompsoni* | Asia | | | | | | KY454712 | KY454730 | | | | KY454746 | | | |
| *Chlorocichla falkensteini* | Africa | | | | | | KY454713 | KY454731 | | | | KY454747 | | | |
| *Chlorocichla flaviventris* | Africa | | | | | AY228053 | GQ242087 | AY590758 | EF626720 | GQ242066 | | AY228290 | EF625309 | AY228009 | |
| *Chlorocichla laetissima* | Africa | | | | HQ998101 | | | | | | | | | | |
| *Chlorocichla prigoginei* | Africa | | | | HQ998159 | | | | | | | | | | |
| *Chlorocichla simplex* | Africa | | | | | | KY454714 | KY454732 | | | | KY454748 | | | |
| *Criniger barbatus* | Africa | AF386487 | AF391224 | | JQ174571 | | DQ402208 | DQ402269 | EF626740 | | | EF625278 | EF625329 | | |
| *Criniger calurus* | Africa | AF386483 | AF391220 | | JQ174574 | | DQ402206 | DQ402267 | | | | DQ125947 | | | GU112529 |
| *Criniger chloronotus* | Africa | | | JX259157 | | | DQ402207 | DQ402268 | | DQ402321 | | JX259184 | | | |
| *Criniger ndussumensis* | Africa | AF386472 | AF391210 | | | | DQ402209 | DQ402270 | EF626741 | DQ402324 | | EF625279 | EF625330 | | |
| *Criniger olivaceus* | Africa | AF386488 | AF391225 | | | | | | | | | | | | |
| *Eurillas ansorgei* | Africa | | | | | | DQ402195 | DQ402256 | EF626708 | DQ402313 | | EF625246 | EF625297 | | |
| *Eurillas curvirostris* | Africa | | | | | | AH005522 | DQ402259 | EF626706 | DQ402359 | | EF625247 | EF625295 | | |
| *Eurillas gracilis* | Africa | | | | | | DQ402196 | DQ402257 | EF626709 | DQ402357 | | EF625245 | EF625298 | | |
| *Eurillas latirostris* | Africa | AF386467 | AF096477 | | | DQ008508 | GQ242089 | GQ242124 | EF626710 | GQ242068 | | | EF625299 | | |
| *Eurillas virens* | Africa | AF386477 | AF391214 | | HQ998164 | AH005519 | DQ402197 | DQ402258 | EF626711 | DQ402358 | | EF625250 | EF625300 | | |
| *Hemixos castanonotus* | Asia | | | | | | GU112647 | GU112693 | | GU112601 | | | | | GU112532 |
| *Hemixos cinereus* | IAA | | | | | | DQ402224 | DQ402285 | GQ242038 | | | GQ242104 | GQ242141 | | GU112567 |
| *Hemixos flavala* | Asia | | | | | KJ456298 | GU112648 | GU112694 | | GU112602 | | | | | GU112533 |
| *Hirundo rustica* | Cosmopolite | AB042349 | AB042382 | GU460221 | FJ027653 | DQ119526 | HF548562 | DQ402247 | EF626748 | DQ402309 | HQ333105 | AY064258 | EF441240 | AY064271 | |
| *Hypsipetes amaurotis* | Japan | EU167068 | | | AB765881 | AB159161 | DQ402222 | DQ402283 | GQ242037 | DQ402325 | | GQ242103 | GQ242140 | | GU112570 |
| *Hypsipetes borbonicus* | Reunion | | | AY590702 | | | | AY590733 | | | | | | | |
| *Hypsipetes crassirostris* | Seychelles | | | AY590703 | | | | AY590741 | | | | | | | |

| | | | | | | | | | | | | | | | |
|---|---|---|---|---|---|---|---|---|---|---|---|---|---|---|---|
| *Hypsipetes everetti* | IAA | | | | | | GU112651 | GU112697 | | GU112605 | | | | | GU112536 |
| *Hypsipetes guimarasensis* | IAA | | | | | | GU112658 | GU112704 | | GU112612 | | | | | GU112543 |
| *Hypsipetes leucocephalus* | Asia | AF386481 | | | | KJ456308 | GU112649 | GU112695 | GQ242040 | GU112603 | | GQ242105 | GQ242143 | | GU112534 |
| *Hypsipetes madagascariensis* | Madagascar Comoros | AB042346 | AB042379 | AY590704 | JQ175126 | | DQ402225 | DQ402286 | | DQ402328 | | | | | GU112568 |
| *Hypsipetes mindorensis* | IAA | | | | | | GU112655 | GU112701 | | GU112609 | | | | | GU112540 |
| *Hypsipetes moheliensis* | Comoros | | | AY590710 | | | | AY590737 | | | | | | | |
| *Hypsipetes olivaceus* | Mauritius | | | AY590709 | | | | AY590735 | | | | | | | |
| *Hypsipetes parvirostris* | Comoros | | | AY590711 | | | | AY590739 | | | | | | | |
| *Hypsipetes philippinus* | IAA | AF386486 | AF391223 | AY590713 | | | GU112654 | GU112700 | EF626742 | GU112608 | | EF625280 | EF625331 | | GU112539 |
| *Hypsipetes rufigularis* | IAA | | | | | | GU112660 | GU112706 | | GU112614 | | | | | GU112545 |
| *Hypsipetes siquijorensis* | IAA | | | | | | GU112661 | GU112707 | | GU112615 | | | | | GU112546 |
| *Iole charlottae* | IAA | | | | | | GU112690 | GU112736 | GQ242036 | DQ402354 | | GQ242102 | GQ242139 | | GU112598 |
| *Iole palawanensis* | IAA | | | | | | GU112688 | GU112734 | | GU112642 | | | | | GU112596 |
| *Iole propinqua* | Asia IAA | AF386475 | AF391212 | | JQ175164 | | GQ369691 | | | | | GQ369646 | | | |
| *Iole viridescens* | Asia | | | | | | KU601856 | KU601803 | | KU601909 | | | | | |
| *Ixonotus guttatus* | Africa | AF386470 | AF391208 | | | | GQ242088 | GQ242123 | EF626718 | GQ242067 | | EF625257 | EF625307 | | |
| *Ixos malaccensis* | IAA | | | | | KY404183 | DQ402227 | DQ402288 | | DQ402329 | | | | | GU112569 |
| *Ixos mcclellandii* | Asia | AF386468 | AF391206 | | | JX398905 | GQ242079 | GQ242113 | GQ242039 | GU112607 | KJ455060 | DQ008558 | GQ242142 | KJ456057 | GU112538 |
| *Ixos philippinus* | IAA | | | | EU541456 | | | | | | | | | EF568257 | |
| *Ixos virescens* | IAA | | | | | | KY454715 | KY454733 | | | | KY454749 | | | |
| *Malia grata* | IAA | | | | | JX398905 | JX398921 | JN826851 | | | | | | | |
| *Neolestes torquatus* | Africa | | | | HQ998140 | | GQ242083 | GQ242117 | GQ242041 | GQ242061 | | GQ242106 | GQ242144 | | |
| *Nicator chloris* | Africa | | | | JQ175554 | JX236396 | DQ402188 | DQ402249 | EU680666 | | | EU680603 | EU680745 | AY319991 | |
| *Phyllastrephus albigula* | Africa | | | | | | HQ716722 | | HQ716832 | | | | | | |
| *Phyllastrephus albigularis* | Africa | AF386476 | AF391213 | | | | DQ402210 | DQ402271 | EF626734 | DQ402330 | | EF625272 | EF625323 | | |
| *Phyllastrephus baumanni* | Africa | | | | | | KY454716 | | | | | | | | |
| *Phyllastrephus cerviniventris* | Africa | | | | | JX236400 | | | EF626726 | | | JX236363 | EF625315 | JX236447 | |
| *Phyllastrephus debilis* | Africa | | | | | AF199383 | DQ402213 | DQ402274 | EF626737 | DQ402335 | HQ716929 | EF625275 | EF625326 | | |
| *Phyllastrephus fischeri* | Africa | | | | | | DQ402216 | DQ402277 | EF626730 | DQ402339 | | EF625266 | EF625319 | | |

| | | | | | | | | | | | | | | | |
|---|---|---|---|---|---|---|---|---|---|---|---|---|---|---|---|
| *Phyllastrephus flavostriatus* | Africa | | | | | AH005523 | DQ402214 | DQ402275 | EF626735 | DQ402336 | | EF625273 | EF625324 | | |
| *Phyllastrephus fulviventris* | Africa | | | | | | KY454717 | | | | | | | | |
| *Phyllastrephus hypochloris* | Africa | | | | | | DQ402215 | DQ402276 | EF626727 | DQ402337 | | EF625265 | EF625316 | | |
| *Phyllastrephus icterinus* | Africa | AF386469 | AF391207 | | JQ175790 | AF199382 | DQ402211 | DQ402272 | EF626731 | DQ402331 | | EF625269 | EF625320 | | GU112530 |
| *Phyllastrephus lorenzi* | Africa | | | | HQ998127 | | | | EF626732 | | | EF625270 | EF625321 | | |
| *Phyllastrephus placidus* | Africa | | | | | | DQ402212 | DQ402273 | EF626729 | DQ402332 | | EF625268 | EF625318 | | |
| *Phyllastrephus poensis* | Africa | | | | | | GQ242084 | GQ242118 | GQ242042 | GQ242062 | | GQ242107 | GQ242145 | | |
| *Phyllastrephus scandens* | Africa | | | | | AF199381 | DQ402218 | DQ402279 | EF626723 | | | EF625261 | EF625312 | | |
| *Phyllastrephus strepitans* | Africa | | | | | | | | EF626725 | | | EF625263 | EF625314 | | |
| *Phyllastrephus terrestris* | Africa | | | | | | DQ402217 | DQ402278 | EF626724 | | | EF625262 | EF625313 | | |
| *Phyllastrephus xavieri* | Africa | | | | | | DQ402219 | AY590761 | EF626733 | DQ402340 | | EF625271 | EF625322 | | |
| *Pycnonotus atriceps* | IAA | AF386484 | AF391221 | | | KY404184 | DQ402231 | DQ402292 | GQ242029 | DQ402341 | | GQ242096 | GQ242132 | | GU112571 |
| *Pycnonotus aurigaster* | Asia IAA | | | | | | KY454718 | KY454734 | | | | KY454750 | | | |
| *Pycnonotus barbatus* | Africa | AF386479 | AF391216 | | FJ473231 | FJ487857 | DQ402232 | DQ402293 | EF626746 | DQ402342 | FJ357922 | EF625284 | EF625335 | AY057027 | GU112572 |
| *Pycnonotus bimaculatus* | IAA | | | | | | KP943466 | | | | | | | | |
| *Pycnonotus blanfordi* | Asia IAA | | | | JQ176058 | | KY454719 | KY454735 | | KP943348 | | KY454751 | | | KP943473 |
| *Pycnonotus brunneus* | IAA | | AY574891 | | FJ473091 | FJ487694 | DQ402233 | DQ402294 | | DQ402343 | | | | | GU112573 |
| *Pycnonotus cafer* | India | KF289822 | KF289828 | | JF498896 | KJ456440 | KJ455616 | | | | KJ455154 | KJ454895 | KJ455904 | KJ456130 | |
| *Pycnonotus capensis* | Africa | | | | | | | | EF626745 | | | EF625283 | EF625334 | | |
| *Pycnonotus cinereifrons* | IAA | | | | | | GU112684 | GU112730 | | GU112638 | | | | | GU112592 |
| *Pycnonotus cyaniventris* | IAA | | | | | KY404185 | DQ402234 | DQ402295 | | DQ402344 | | | | | GU112574 |
| *Pycnonotus dispar* | IAA | | | | | | KP943427 | | | | | | | | |
| *Pycnonotus erythropthalmos* | IAA | | | | | FJ487794 | DQ402235 | DQ402296 | | DQ402345 | | | | | GU112575 |
| *Pycnonotus eutilotus* | IAA | | AY574900 | | | KY404186 | DQ402236 | DQ402297 | GQ242030 | DQ402346 | | GQ242097 | GQ242133 | | GU112576 |
| *Pycnonotus finlaysoni* | IAA | AF386473 | AF391211 | | FJ473154 | FJ487764 | GQ242076 | GQ242110 | GQ242033 | GQ242053 | | GQ242100 | GQ242136 | | |
| *Pycnonotus flavescens* | Asia | | | | | KY404187 | GU112669 | GU112715 | | GU112623 | | | | | GU112554 |
| *Pycnonotus flaviventris* | Asia India | KJ186975 | KJ186975 | KJ186975 | KJ186975 | KJ186975 | KJ186975 | KJ186975 | GQ242032 | | | GQ242099 | GQ242135 | | |
| *Pycnonotus goiavier* | IAA | | | | FJ473057 | FJ487665 | DQ402237 | DQ402298 | | DQ402347 | | | | | KY454739 |
| *Pycnonotus gularis* | India | | | | | | KP943372 | | | | | | | | |

| | | | | | | | | | | | | | | | |
|---|---|---|---|---|---|---|---|---|---|---|---|---|---|---|---|
| *Pycnonotus jocosus* | Asia IAA India | AF386480 | AF391217 | | GU170351 | KJ456441 | GQ242077 | GQ242111 | GQ242034 | GU112624 | KJ455155 | DQ008557 | GQ242137 | KJ456131 | GU112555 |
| *Pycnonotus leucogenys* | Asia India | | | | HQ168045 | | DQ402241 | DQ402302 | | DQ402351 | KJ455156 | KJ454896 | KJ455905 | EF568258 | GU112578 |
| *Pycnonotus leucogrammicus* | IAA | | | | | | KP943465 | | | | | | | | |
| *Pycnonotus leucotis* | Arabic Peninsula India | | | | | | KP943428 | | | | | | | | |
| *Pycnonotus luteolus* | India | | | | | | KP943367 | | | | | | | | |
| *Pycnonotus melanicterus* | India | | | | | | KY454720 | | | | | | | KJ456132 | |
| *Pycnonotus melanoleucos* | IAA | | | | | | DQ402242 | DQ402303 | | DQ402352 | | | | | GU112579 |
| *Pycnonotus montis* | IAA | | | | | KY404188 | DQ402243 | DQ402304 | | DQ402353 | KJ455157 | | | | |
| *Pycnonotus nigricans* | Africa | | | | | | DQ402238 | DQ402299 | | DQ402348 | | | | | GU112580 |
| *Pycnonotus plumosus* | IAA | | AY574884 | | FJ473241 | FJ487748 | DQ402239 | DQ402300 | JN826138 | DQ402349 | | | | | JN826387 |
| *Pycnonotus priocephalus* | India | | | | | | KY454721 | | | | | | | | |
| *Pycnonotus simplex* | IAA | | | | FJ473143 | FJ487754 | GU112671 | GU112717 | | GU112625 | | | | | GU112556 |
| *Pycnonotus sinensis* | Asia | GU475148 | GU475148 | GU475148 | GU475148 | GU475148 | GU475148 | GU475148 | | GU112626 | | | | | GU112557 |
| *Pycnonotus squamatus* | IAA | | | | | KY404189 | KY454722 | GU112737 | | GU112645 | | | | | KY454740 |
| *Pycnonotus striatus* | Asia | | | | | KJ456444 | KJ455620 | | | | KJ455158 | KJ454898 | KJ455907 | KJ456133 | |
| *Pycnonotus taivanus* | Taiwan | FJ378536 | FJ378536 | FJ378536 | FJ378536 | FJ378536 | FJ378536 | FJ378536 | | | | | | | |
| *Pycnonotus tricolor* | Africa | | | | | | KY454723 | KY454736 | | | | KY454752 | | | |
| *Pycnonotus urostictus* | IAA | | | | EU541463 | | KJ931003 | AY590762 | EF626744 | GU112629 | | EF625282 | EF625333 | | GU112560 |
| *Pycnonotus xanthopygos* | Arabic Peninsula | | | | | | KP943432 | | | | | | | | |
| *Pycnonotus xanthorrhous* | Asia | AF386485 | AF391222 | | | | GU112677 | GU112723 | | GU112631 | | | | | GU112562 |
| *Pycnonotus zeylanicus* | IAA | | | | | | DQ402240 | DQ402301 | | DQ402350 | | | | | GU112582 |
| *Setornis criniger* | IAA | | | | | KY404190 | DQ402221 | DQ402282 | | DQ402355 | | | | | GU112583 |
| *Spizixos canifrons* | Asia | | | | | | KY454724 | KY454737 | | | | | | | |
| *Spizixos semitorques* | Asia | | | | FJ661100 | JF509584 | DQ402244 | DQ402305 | GQ242031 | DQ402356 | | GQ242098 | GQ242134 | | GU112584 |
| *Stelgidillas gracilirostris* | Africa | | | | | | GQ242082 | GQ242116 | EF626705 | GQ242060 | | EF625249 | EF625294 | | |
| *Thapsinillas affinis* | IAA | | | | | | KY454725 | | | | | | | | |
| *Thescelocichla leucopleura* | Africa | | | | | | DQ402200 | DQ402261 | EF626722 | DQ402312 | | EF625260 | EF625311 | | |
| *Tricholestes criniger* | IAA | AF386489 | AF391226 | | | KY404191 | DQ402223 | DQ402284 | GQ242035 | DQ402326 | | GQ242101 | GQ242138 | | GU112585 |

The calibration for this analysis was based on a Passeriformes dating analysis (Ericson, Klopfstein, Irestedt, Nguyen, & Nylander, 2014). Thus, a prior was set on the root with a lognormal distribution (M = 3.631 and S = 0.13), so that the median would be equal to 37.8 m.y.a. and the 95% confidence interval would be between 29 and 48 m.y.a. These dates represent the estimated origin date of the sylvioidea, the less inclusive group that includes the Pycnonotidae and the outgroup (Alström, Ericson, Olsson, & Sundberg, 2006).

#### 6. Atyidae

Table S6: Genbank numbers for the sequences used in the Atyidae analyses with the distribution attributed to each terminal.

| Species | Locality | 16S | 28S | H3 |
|---|---|---|---|---|
| Antecaridina lauensis | Australia IAA Pacific | EU123851 | | FN995457 |
| Antecaridina sp | Australia IAA Pacific | FN995353 | | FN995458 |
| Antecaridina sp | Australia IAA Pacific | EU123853 | | FN995459 |
| Atya gabonensis | Africa America | EF489961 | FN995546 | FN995460 |
| Atya gabonensis | Africa America | EF489989 | FN995547 | FN995461 |
| Atya margaritacea | America | EF489983 | FN995548 | FN995462 |
| Atya ortmannioides | America | EF489993 | FN995549 | FN995463 |
| Atya scabra | America | EF489985 | FN995550 | FN995465 |
| Atyaephyra desmarestii | Europe | FN995354 | FN995551 | FN995466 |
| Atyella brevirostris | Africa | | FN995552 | FN995468 |
| Atyoida bisulcata | Pacific | EF489995 | FN995553 | FN995469 |
| Atyoida pilipes | Pacific | DQ681277 | FN995554 | FN995470 |
| Atyoida pilipes | Pacific | DQ681279 | FN995555 | FN995471 |
| Atyopsis moluccensis | IAA | DQ681280 | FN995557 | FN995473 |
| Atyopsis spinipes | IAA Pacific | DQ681282 | | |
| Australatya striolata | Australia | AY795035 | FN995558 | FN995474 |
| Caridella paski | Africa | FN995355 | FN995559 | FN995475 |
| Caridina cantonensis | Asia | DQ478487 | FN995560 | FN995476 |
| Caridina celebensis | IAA Pacific | FN995356 | FN995561 | FN995477 |
| Caridina cf africana | Africa | DQ478483 | FN995562 | FN995478 |
| Caridina endehensis | IAA | FN995357 | FN995563 | FN995479 |
| Caridina ensifera | IAA | AM747734 | FN995564 | FN995480 |
| Caridina gracilipes | Asia IAA | FN995358 | FN995565 | FN995481 |
| Caridina gracilirostris | IAA India Madagascar | EU873516 | FN995566 | FN995482 |
| Caridina serrata | Asia | DQ478512 | | FN995483 |
| Caridina serratirostris | Comoros IAA Madagascar Pacific | DQ478513 | FN995567 | FN995484 |
| Caridina serratirostris | Comoros IAA Madagascar Pacific | DQ478514 | FN995568 | FN995485 |
| Caridina serratirostris | Comoros IAA Madagascar Pacific | DQ478516 | FN995569 | FN995486 |
| Caridina sp | Australia | DQ478534 | FN995570 | EU123809 |
| Caridina sp2 | Asia | FN995359 | FN995571 | |
| Caridina steineri | Madagascar | DQ681274 | | |

| | | | | |
|---|---|---|---|---|
| Caridina sumatrensis | IAA | FN995360 | FN995572 | FN995487 |
| Caridina thomasi | IAA | EU873523 | FN995573 | FN995488 |
| Caridina togoensis | Africa | FN995361 | FN995574 | FN995489 |
| Caridina trifasciata | Asia | FN995362 | FN995575 | FN995490 |
| Caridina typus ARC | Australia IAA India Mascarene Pacific Seychelles | KY069381 | KY069755 | |
| Caridina typus SUL | IAA | KY069384 | KY069757 | |
| Caridina typus TAL | IAA | KY069360 | KY069741 | |
| Caridina villadolidi | IAA | KY436221 | | |
| Caridinides wilkinsi | Australia | DQ681273 | FN995577 | FN995492 |
| Caridinopsis chevalieri | Africa | | FN995578 | |
| Dugastella marocana | Africa | FJ594357 | FN995579 | FN995494 |
| Dugastella marocana | Africa | FN995363 | FN995580 | FN995495 |
| Dugastella valentina | Europe | FN995364 | FN995581 | FN995496 |
| Dugastella valentina | Europe | FN995365 | FN995582 | FN995497 |
| Edoneus marulas | IAA | FN995366 | FN995583 | FN995498 |
| Edoneus marulas | IAA | FN995367 | FN995584 | FN995499 |
| Gallocaris inermis | Europe | DQ641599 | | |
| Halocaridina rubra | Pacific | FN995368 | FN995585 | FN995500 |
| Halocaridinides trigonophthalma | Pacific | FN995369 | FN995586 | FN995501 |
| Jonga serrei | America | EF490003 | FN995587 | FN995502 |
| Lancaris singhalensis | India | FN995370 | FN995588 | FN995503 |
| Limnocaridina parvula | Africa | FN995371 | FN995589 | FN995504 |
| Limnocaridina similis | Africa | FN995372 | FN995590 | FN995505 |
| Limnocaridina tanganyikae | Africa | FN995373 | FN995591 | FN995506 |
| Marosina longirostris | IAA | FN995374 | FN995592 | FN995507 |
| Marosina longirostris | IAA | FN995375 | FN995593 | FN995508 |
| Micratya poeyi | America | EF489991 | FN995595 | FN995510 |
| Micratya sp | America | EF489994 | FN995596 | FN995511 |
| Neocaridina cf zhangjiajiensis | Asia | | FN995597 | FN995512 |
| Neocaridina denticulata | Asia | DQ681268 | FN995598 | FN995513 |
| Neocaridina denticulata | Asia | FN995377 | FN995599 | FN995514 |
| Neocaridina palmata | Asia | | FN995601 | FN995516 |
| Neocaridina saccam | Asia | DQ681270 | | |
| Palaemonias alabamae | America | FN995378 | FN995602 | FN995517 |
| Palaemonias alabamae | America | FN995379 | FN995603 | FN995518 |
| Palaemonias alabamae | America | FN995380 | FN995604 | FN995519 |
| Palaemonias alabamae | America | FN995381 | FN995605 | FN995520 |
| Palaemonias sp | America | FN995382 | FN995606 | FN995521 |
| Palaemonias sp2 | America | FN995383 | FN995607 | FN995522 |
| Paracaridina sp | Asia | FN995384 | FN995608 | FN995523 |
| Paratya australiensis | Australia | EF490007 | FN995610 | FN995525 |
| Paratya curvirostris | New Zealand | AY661475 | FN995611 | FN995526 |
| Parisia gracilis | Australia | EU123843 | FN995612 | EU123810 |

| | | | | |
|---|---|---|---|---|
| Parisia unguis | Australia | DQ681288 | | FN995527 |
| Potimirim glabra | America | EF489999 | FN995613 | FN995528 |
| Potimirim potimirim | America | FN995386 | FN995614 | FN995529 |
| Potimirim sp | America | FN995387 | FN995615 | FN995530 |
| Potimirim sp2 | America | EF490000 | FN995616 | FN995531 |
| Pycneus morsitans | Australia | EU123849 | FN995617 | EU123849 |
| Pycnisia raptor | Australia | DQ681271 | FN995618 | FN995532 |
| Sinodina sp | Asia | FN995388 | FN995619 | FN995533 |
| Stygiocaris lancifera | Australia | EU123826 | FN995620 | EU123804 |
| Stygiocaris sp | Australia | EU123840 | FN995621 | EU123807 |
| Stygiocaris stylifera | Australia | EU123836 | FN995622 | EU123806 |
| Syncaris pacifica | America | FN995389 | FN995623 | FN995534 |
| Syncaris pacifica | America | FN995390 | FN995624 | FN995535 |
| Troglocaris anophtalmus | Europe | DQ641587 | DQ641614 | |
| Troglocaris bosnica | Europe | DQ641588 | DQ641615 | |
| Troglocaris kutaissiana | Europe | DQ320026 | | |
| Troglocaris planinensis | Europe | DQ320024 | | |
| Troglocaris prasence | Europe | DQ641592 | DQ641618 | |
| Troglocaris pretneri | Europe | DQ641590 | DQ641616 | |
| Troglocaris sp | Europe | DQ641593 | DQ641619 | |
| Typhlatya miravetensis | Europe | DQ641602 | DQ641626 | |
| Typhlatya mitchelli | America | FN995393 | FN995627 | FN995538 |
| Typhlatya rogersi | America | FN995394 | FN995628 | FN995539 |
| Typhlatya sp | America | FN995395 | FN995629 | FN995540 |
| Typhlatya sp2 | America | FN995396 | FN995630 | FN995541 |
| Xiphocaris elongata | America | EF490010 | FN995631 | FN995542 |
| Xiphocaris elongata | America | EF490011 | FN995632 | FN995543 |

In the alignments, the three clades from Bernardes et al. (2017) were treated as separated terminals. Two calibration points were used from the three alternatives in the original article (von Rintelen et al., 2012): *Atyoida roxoi* was assigned to stem of Atyidae with a lognormal distribution (M = 1.25 and S = 0.68) set to encompass the Aptian age (offset = 112 m.y.a.); the secondary calibration from Porter, Pérez-Losada, & Crandall (2005) was assigned to the root with a lognormal distribution (M=3.95 and S = 0.30; offset = 217 m.y.a.). The calibration based in the Palaemonidae fossils was not used because we decided to reduce the dataset to simplify the analyses.

#### 7. Caecilia

The sequences used in the Caecilia analyses were downloaded directly from the original article's dryad repository (https://datadryad.org/resource/doi:10.5061/dryad.jm453), separated from other amphibian groups, and re-aligned. Proopiomelanocortin and Rhodopsin genes were removed from the analyses because they represented no/little data for the caecilians specifically. We used several points of calibration. First, we used calibration points for Tetrapoda (lognormal distribution; M = 1.8 and S = 0.44; offset = 337 m.y.a.) and for Batrachia (exponential distribution; mean = 8.0 and offset = 260 m.y.a.) based on Benton and Donoghue (2007), Pyron (2014), Benton et al. (2015), and Feng et al. (2017). We placed a calibration in the Caecilia most recent ancestor node (lognormal distribution; M = 1.1 and S = 1.63; offset = 100 m.y.a.) based on Wiens (2011). In the original article, these dates were obtained with a dated amphibian tree and not only were these analyses robust but they were also in agreement with several other studies we analysed (e.g. (Feng et al., 2017). Calibration using the fossil from Early Cretaceous *Eocaecilia micropodia* (Jenkins & Walsh, 1993) was tested by assigning it either to the node of the Lissamphibia ancestor with a lognormal distribution or to the node of the Caecilia ancestor (M = 1.6 and S = 0.87; offset = 174 m.y.a.). *Eocaecilia* is stem-group caecilian (Maddin, Jenkins Jr, & Anderson, 2012) and should not be used to date the crown-group. Our tests showed that adding *Eocaecilia* in the Lissamphibia node made no difference to the final result. However, to treat *Eocaecilia* as a crown-group caecilian places the divergence between Indian and Seychellois taxa in a period that is congruent with the geological dates of the India-Seychelles separation.

#### 8. Neobatrachia

Table S7: Genbank numbers for the sequences used in the Neobatrachia analyses with the distribution attributed to each terminal.

| Species | Locality | CILP | DISP2 | DMXL1 | DSEL | EVPL | EXOC8 | HYP | MSH6 | PPL | RAG1 | RAG2 | SALL1 | WFIKKN2 | ZFPM2 | CXCR4 |
|---|---|---|---|---|---|---|---|---|---|---|---|---|---|---|---|---|
| *Acris crepitans* | Seychelles | KX200136 | KX201145 | KX201295 | KX201817 | KX202098 | KX202244 | KX204391 | KX207070 | KX208566 | KX208710 | KX208856 | KX209700 | KX211266 | KX211700 | KX200424 |
| *Adenomus kelaartii* | India | | | | | | | | | | | EF107284 | | | | EF107447 |
| *Afrixalus dorsalis* | Africa | | | | | | | | | | | DQ347236 | | | | EF107456 |
| *Agalychnis callidryas* | Seychelles | KX200097 | KX201101 | KX201246 | KX201770 | KX202055 | KX202199 | KX204345 | KX207026 | KX208518 | KX208666 | KX208812 | KX209657 | KX211218 | KX211654 | KX200375 |
| *Agalychnis lemur* | Seychelles | KX200085 | KX201089 | KX201232 | KX201756 | KX202041 | KX202184 | KX204333 | KX207015 | KX208503 | KX208652 | KX208799 | KX209643 | KX211203 | KX211638 | KX200360 |
| *Agalychnis saltator* | Seychelles | | | | | | | | | | | EF174315 | | | | GQ365983 |
| *Aglyptodactylus madagascariensis* | Madagascar | KX200146 | KX201156 | KX201306 | KX201828 | KX202109 | KX202255 | KX204402 | KX207080 | KX208576 | KX208721 | KX208867 | KX209711 | KX211277 | KX211711 | KX200434 |
| *Allobates femoralis* | Seychelles | | KX201120 | KX201270 | KX201792 | KX202073 | KX202221 | KX204367 | KX207047 | KX208541 | KX208686 | KX208833 | KX209675 | KX211240 | KX211675 | KX200398 |
| *Allophryne ruthveni* | Seychelles | | | | | | | | | | | EU663432 | | | | KF534373 |
| *Alsodes gargola* | Seychelles | KX200128 | KX201134 | KX201284 | KX201806 | KX202086 | KX202235 | KX204381 | KX207061 | KX208555 | KX208700 | KX208845 | KX209689 | KX211254 | KX211688 | KX200412 |
| *Alytes obstetricans* | America | KX200147 | KX201157 | KX201307 | KX201829 | KX202110 | KX202256 | KX204403 | KX207081 | KX208577 | KX208722 | | KX209712 | KX211278 | KX211712 | KX200435 |
| *Amazophrynella minuta* | Seychelles | | | | | | | | | | DQ158346 | | | | KX211639 | DQ306496 |
| *Amietia lubrica* | Africa | KX200201 | KX201212 | KX201362 | KX201886 | KX202164 | KX202312 | KX204460 | KX207133 | KX208633 | KX208779 | KX208918 | KX209770 | KX211334 | KX211769 | KX200492 |
| *Amolops loloensis* | Asia | KX200118 | KX201124 | KX201274 | KX201796 | KX202077 | KX202225 | KX204371 | KX207051 | KX208545 | KX208690 | KX208836 | KX209679 | KX211244 | | KX200402 |
| *Amolops ricketti* | Asia | | KX201159 | KX201309 | KX201831 | | KX202258 | KX204405 | KX207083 | KX208579 | KX208724 | KX208869 | KX209714 | KX211280 | KX211714 | KX200437 |
| *Anaxyrus canorus* | Seychelles | | KX201093 | | KX201760 | KX202045 | KX202188 | KX204337 | KX207018 | KX208507 | KX208656 | KX208803 | KX209647 | KX211207 | KX211643 | KX200364 |
| *Anaxyrus punctatus* | Seychelles | KX200104 | KX201109 | KX201258 | KX201781 | KX202061 | KX202210 | KX204355 | KX207035 | KX208529 | KX208674 | | KX209664 | KX211229 | KX211663 | KX200386 |
| *Andrias davidianus* | | KX200159 | KX201170 | KX201319 | KX201842 | KX202121 | | KX204416 | KX207093 | KX208590 | KX208734 | KX208880 | KX209725 | KX211291 | KX211725 | KX200447 |
| *Anodonthyla boulengerii* | Madagascar | | | KX201252 | KX201776 | | KX202205 | | KX207031 | KX208524 | | | | | | |
| *Aplastodiscus perviridis* | Seychelles | KX200126 | KX201132 | KX201282 | KX201804 | KX202084 | KX202233 | KX204379 | KX207059 | KX208553 | KX208698 | KX208843 | KX209687 | KX211252 | KX211686 | KX200410 |
| *Arthroleptis poecilonotus* | Africa | KX200148 | KX201158 | KX201308 | KX201830 | KX202111 | KX202257 | KX204404 | KX207082 | KX208578 | KX208723 | KX208868 | KX209713 | KX211279 | KX211713 | KX200436 |
| *Arthroleptis variabilis* | Africa | | | | | | | | | | | EF396073 | | | | AY364180 |
| *Ascaphus truei* | Seychelles | KX200149 | KX201160 | KX201310 | KX201832 | KX202112 | | KX204406 | KX207084 | KX208580 | KX208725 | KX208870 | KX209715 | KX211281 | KX211715 | KX200438 |
| *Astylosternus diadematus* | Africa | KX200145 | KX201155 | KX201305 | KX201827 | KX202108 | KX202254 | KX204401 | KX207079 | | KX208720 | KX208866 | KX209710 | KX211276 | KX211710 | KX200433 |
| *Atelognathus reverberii* | Seychelles | KX200139 | KX201149 | KX201299 | KX201821 | KX202102 | KX202248 | KX204395 | KX207074 | KX208570 | KX208714 | KX208860 | KX209704 | KX211270 | KX211704 | KX200428 |
| *Atelopus peruensis* | Seychelles | | | | | | | | | | | DQ158345 | | | | DQ306495 |
| *Aubria subsigillata* | Africa | KX200138 | KX201147 | KX201297 | KX201819 | KX202100 | KX202246 | KX204393 | KX207072 | KX208568 | KX208712 | KX208858 | KX209702 | KX211268 | KX211702 | KX200426 |
| *Babina chapaensis* | Asia | | | | | | | | | | | EU076752 | | | | KR264317 |
| *Babina pleuraden* | Asia | | | | | | | | | | | KR264384 | | | | KR264303 |
| *Barbourula busuangensis* | IAA | KX200143 | KX201153 | KX201303 | KX201825 | KX202106 | KX202252 | KX204399 | KX207077 | KX208574 | KX208718 | KX208864 | KX209708 | KX211274 | KX211708 | KX200431 |
| *Barycholos pulcher* | Seychelles | KX200094 | KX201099 | KX201241 | KX201766 | KX202051 | KX202194 | KX204343 | KX207023 | KX208513 | KX208662 | KX208808 | KX209653 | KX211213 | KX211649 | KX200370 |
| *Batrachophrynus macrostomus* | Seychelles | KX200122 | KX201128 | KX201278 | KX201800 | KX202081 | KX202229 | KX204375 | KX207055 | KX208549 | KX208694 | | KX209683 | KX211248 | KX211682 | KX200406 |
| *Batrachyla leptopus* | Seychelles | KX200134 | KX201142 | KX201292 | KX201814 | KX202095 | KX202242 | KX204389 | KX207067 | KX208563 | KX208707 | | KX209697 | KX211263 | KX211697 | KX200421 |
| *Batrachyla taeniata* | Seychelles | KX200141 | KX201151 | KX201301 | KX201823 | KX202104 | KX202250 | KX204397 | | KX208572 | KX208716 | KX208862 | KX209706 | KX211272 | KX211706 | KX200429 |
| *Blommersia wittei* | Madagascar | | | | | | | | | | | AY323774 | | | | AB612028 |
| *Bombina fortinuptialis* | Asia America | KX200116 | KX201122 | KX201272 | KX201794 | KX202075 | KX202223 | KX204369 | KX207049 | KX208543 | KX208688 | KX208835 | KX209677 | KX211242 | KX211677 | KX200400 |

| | | | | | | | | | | | | | | | | |
|---|---|---|---|---|---|---|---|---|---|---|---|---|---|---|---|---|
| *Bombina orientalis* | Asia America | KX200154 | KX201165 | KX201315 | KX201837 | KX202117 | KX202263 | KX204411 | KX207089 | KX208585 | KX208730 | KX208875 | KX209720 | KX211286 | KX211720 | KX200442 |
| *Boophis madagascariensis* | Comoros Madagascar | KX200095 | | KX201242 | KX201767 | | KX202195 | | | KX208514 | | KX208809 | | KX211214 | KX211650 | KX200371 |
| *Boophis xerophilus* | Comoros Madagascar | | | | | | | | | | | AY364209 | | | | AY364179 |
| *Brachycephalus ephippium* | Seychelles | | | | | | | | | | | GQ345275 | | | | GQ345180 |
| *Brachytarsophrys feae* | Asia | KX200150 | KX201161 | KX201311 | KX201833 | KX202113 | KX202259 | KX204407 | KX207085 | KX208581 | KX208726 | KX208871 | KX209716 | KX211282 | KX211716 | |
| *Breviceps macrops* | Africa | KX200152 | KX201163 | KX201313 | KX201835 | KX202115 | KX202261 | KX204409 | KX207087 | KX208583 | KX208728 | KX208873 | KX209718 | KX211284 | KX211718 | KX200440 |
| *Breviceps mossambicus* | Africa | | | | | | | | | | | EF396076 | | | | EF017965 |
| *Buergeria buergeri* | Asia | | | | | | | | | | | AB612031 | | | | AB612035 |
| *Buergeria oxycephala* | Asia | KX200090 | KX201095 | KX201237 | KX201762 | KX202047 | KX202190 | KX204339 | KX207020 | KX208509 | KX208658 | KX208805 | KX209649 | KX211209 | KX211645 | KX200366 |
| *Bufo bufo* | America | | | | | | | | | | | AY583336 | | | | DQ306504 |
| *Bufo gargarizans* | Asia | KX200151 | KX201162 | KX201312 | KX201834 | KX202114 | KX202260 | KX204408 | KX207086 | KX208582 | KX208727 | KX208872 | KX209717 | KX211283 | KX211717 | KX200439 |
| *Bufo japonicus* | Asia | | | | | | | | | | | AB612057 | | | | AB612061 |
| *Bufotes viridis* | Asia India America | | | | | | | | | | | KJ609682 | | | | FJ882714 |
| *Calluella guttulata* | IAA | | | | | | | | | | | AB611861 | | | | EF017975 |
| *Callulina kreffti* | Africa | | KX201143 | KX201293 | KX201815 | KX202096 | | | KX207068 | KX208564 | KX208708 | KX208854 | KX209698 | KX211264 | KX211698 | KX200422 |
| *Callulina laphami* | Africa | | | | | | | | | | | KF990044 | | | | KF954751 |
| *Callulops wilhelmanus* | IAA | | KX201102 | KX201248 | KX201772 | | KX202201 | KX204347 | KX207027 | KX208520 | KX208667 | KX208814 | | KX211220 | KX211655 | KX200377 |
| *Calyptocephalella gayi* | Seychelles | KX200113 | KX201118 | KX201268 | KX201790 | KX202071 | KX202219 | KX204365 | KX207045 | KX208539 | KX208684 | KX208831 | KX209673 | KX211238 | KX211673 | KX200396 |
| *Centrolene bacatum* | Seychelles | | | | | | | | | | | EU663437 | | | | KF534379 |
| *Centrolene daidaleum* | Seychelles | | | | | | | | | | | EU663465 | | | | KF534381 |
| *Ceratophrys cornuta* | Seychelles | KX200155 | KX201166 | KX201316 | KX201838 | KX202118 | KX202264 | KX204412 | | KX208586 | KX208731 | KX208876 | KX209721 | KX211287 | KX211721 | KX200443 |
| *Ceratophrys ornata* | Seychelles | | | | | | | | | | | AY364218 | | | | AY364188 |
| *Ceuthomantis smaragdinus* | Seychelles | | | | | | | | | | | GQ345287 | | | | GQ345192 |
| *Chalcorana macrops* | IAA | | | | | | | | | | | KR264416 | | | | KR264339 |
| *Chaperina fusca* | IAA | | | | | | | | | | | AB611865 | | | | AB611870 |
| *Chiasmocleis ventrimaculata* | Seychelles | KX200089 | KX201094 | KX201236 | KX201761 | KX202046 | KX202189 | KX204338 | KX207019 | KX208508 | KX208657 | KX208804 | KX209648 | KX211208 | KX211644 | KX200365 |
| *Chimerella mariaelenae* | Seychelles | | | | | | | | | | | EU663449 | | | | KF534391 |
| *Cochranella granulosa* | Seychelles | | | | | | | | | | | EF107297 | | | | EF107460 |
| *Conraua crassipes* | Africa | KX200156 | KX201167 | KX201317 | KX201839 | KX202119 | KX202265 | KX204413 | KX207090 | KX208587 | KX208732 | KX208877 | KX209722 | KX211288 | KX211722 | KX200444 |
| *Cophixalus cheesmanae* | Australia IAA | KX200142 | KX201152 | KX201302 | KX201824 | KX202105 | KX202251 | KX204398 | KX207076 | KX208573 | KX208717 | KX208863 | KX209707 | KX211273 | KX211707 | KX200430 |
| *Cophixalus cryptotympanum* | Australia IAA | | | | | | | | | | | AB611873 | | | | AB611878 |
| *Cophixalus sp.* | Australia IAA | KX200099 | KX201104 | KX201250 | KX201774 | KX202057 | KX202203 | KX204349 | KX207029 | KX208522 | KX208669 | KX208816 | KX209659 | KX211222 | KX211657 | KX200379 |
| *Cophixalus sp. B* | Australia IAA | | | | | | | | | | | DQ347276 | | | | EF017967 |
| *Cornufer pelewensis* | Australia IAA | KX200197 | KX201207 | KX201357 | KX201881 | KX202160 | KX202307 | KX204455 | KX207129 | KX208628 | KX208774 | KX208915 | KX209765 | KX211329 | KX211764 | KX200487 |
| *Craugastor augusti* | Seychelles | | KX201177 | KX201326 | KX201849 | KX202128 | KX202274 | KX204423 | KX207099 | KX208597 | KX208741 | KX208887 | KX209732 | KX211298 | KX211732 | KX200454 |
| *Craugastor fitzingeri* | Seychelles | KX200120 | KX201126 | KX201276 | KX201798 | KX202079 | KX202227 | KX204373 | KX207053 | KX208547 | KX208692 | KX208838 | KX209681 | KX211246 | KX211680 | KX200404 |
| *Craugastor podiciferus* | Seychelles | | | | | | | | | | | GQ345277 | | | | GQ345182 |
| *Crinia signifera* | Australia | KX200158 | KX201169 | | KX201841 | | KX202267 | KX204415 | KX207092 | KX208589 | KX208733 | KX208879 | KX209724 | KX211290 | KX211724 | KX200446 |
| *Cruziohyla calcarifer* | Seychelles | | | | | | | | | | | EF174317 | | | | GQ365984 |
| *Cryptobatrachus boulengeri* | Seychelles | KX200157 | KX201168 | KX201318 | KX201840 | KX202120 | KX202266 | KX204414 | KX207091 | KX208588 | | KX208878 | KX209723 | KX211289 | KX211723 | KX200445 |
| *Ctenophryne geayi* | Seychelles | | | | | | | | | | | AB611881 | | | | AB611886 |

| | | | | | | | | | | | | | | | | |
|---|---|---|---|---|---|---|---|---|---|---|---|---|---|---|---|---|
| *Cyclorana maini* | Australia | | | | | | | | | | | EF107311 | | | | EF107475 |
| *Dendrobates auratus* | Seychelles | | | | | | | | | | | AY364214 | | | | AY364184 |
| *Dendropsophus parviceps* | Seychelles | KX200110 | KX201115 | KX201265 | KX201788 | KX202068 | KX202216 | KX204362 | KX207042 | KX208536 | KX208681 | KX208828 | KX209670 | KX211235 | KX211670 | KX200393 |
| *Diasporus diastema* | Seychelles | | | | | | | | | | | GQ345279 | | | | GQ345184 |
| *Discoglossus pictus* | America | KX200093 | KX201098 | KX201240 | KX201765 | KX202050 | KX202193 | KX204342 | | KX208512 | KX208661 | | KX209652 | KX211212 | KX211648 | KX200369 |
| *Duttaphrynus melanostictus* | Asia IAA India | KX200153 | KX201164 | KX201314 | KX201836 | KX202116 | KX202262 | KX204410 | KX207088 | KX208584 | KX208729 | KX208874 | KX209719 | KX211285 | KX211719 | KX200441 |
| *Duttaphrynus stomaticus* | Asia IAA India | | | | | | | | | | | KJ609680 | | | | FJ882681 |
| *Dyscophus antongilii* | America | KX200160 | KX201171 | KX201320 | KX201843 | KX202122 | KX202268 | KX204417 | | KX208591 | KX208735 | KX208881 | KX209726 | KX211292 | KX211726 | KX200448 |
| *Elachistocleis ovalis* | Seychelles | KX200161 | KX201172 | KX201321 | KX201844 | KX202123 | KX202269 | KX204418 | KX207094 | KX208592 | KX208736 | KX208882 | KX209727 | KX211293 | KX211727 | KX200449 |
| *Eleutherodactylus coqui* | Seychelles | | | | | | | | | | | EF107341 | | | | EF107500 |
| *Eleutherodactylus marnockii* | Seychelles | | | | | | | | | | | EF107300 | | | | EF107463 |
| *Eleutherodactylus planirostris* | Seychelles | KX200130 | KX201137 | KX201288 | KX201809 | KX202090 | KX202237 | KX204384 | KX207063 | KX208558 | | KX208849 | KX209692 | KX211258 | KX211692 | KX200416 |
| *Epidalea calamita* | America | | | | | | | | | | | EU497610 | | | | FJ882709 |
| *Epipedobates tricolor* | Seychelles | | | | | | | | | | | EF107295 | | | | EF107458 |
| *Espadarana prosoblepon* | Seychelles | | | | | | | | | | | AY364223 | | | | AY364193 |
| *Euphlyctis cyanophlyctis* | Asia India America | | | | | | | | | | | AB488957 | | | | AB488919 |
| *Eupsophus calcaratus* | Seychelles | KX200105 | KX201110 | KX201259 | KX201782 | KX202062 | KX202211 | KX204356 | KX207036 | KX208530 | KX208675 | KX208822 | KX209665 | KX211230 | KX211664 | KX200387 |
| *Fejervarya granosa* | Asia IAA India | | | | | | | | | | | AB488947 | | | | AB488913 |
| *Fejervarya limnocharis* | Asia IAA India | KX200163 | KX201174 | KX201323 | KX201846 | KX202125 | KX202271 | KX204420 | KX207096 | KX208594 | KX208738 | KX208884 | KX209729 | KX211295 | KX211729 | KX200451 |
| *Fejervarya multistriata* | Asia IAA India | | | | | | | | | | | AB500221 | | | | AB500243 |
| *Flectonotus fitzgeraldi* | Seychelles | | | | | | | | | | | DQ679274 | | | | GQ345177 |
| *Gastrophryne olivacea* | Seychelles | KX200164 | KX201175 | KX201324 | KX201847 | KX202126 | KX202272 | KX204421 | KX207097 | KX208595 | KX208739 | KX208885 | KX209730 | KX211296 | KX211730 | KX200452 |
| *Gastrophrynoides immaculatus* | IAA | | | | | | | | | | | AB611901 | | | | AB611906 |
| *Gastrotheca pseustes* | Seychelles | KX200165 | KX201176 | KX201325 | KX201848 | KX202127 | KX202273 | KX204422 | KX207098 | KX208596 | KX208740 | KX208886 | KX209731 | KX211297 | KX211731 | KX200453 |
| *Gastrotheca weinlandii* | Seychelles | | | KX201244 | KX201769 | KX202053 | KX202197 | | | KX208516 | KX208664 | | KX209655 | KX211216 | KX211652 | KX200373 |
| *Heleophryne purcelli* | Africa | KX200169 | KX201181 | KX201330 | KX201853 | KX202132 | KX202278 | KX204427 | KX207103 | | KX208745 | KX208891 | KX209736 | KX211302 | KX211736 | KX200458 |
| *Hemiphractus bubalus* | Seychelles | | | | | | | | | | | DQ679303 | | | | GQ345179 |
| *Hemisus marmoratus* | Africa | KX200168 | KX201180 | KX201329 | KX201852 | KX202131 | KX202277 | KX204426 | KX207102 | KX208600 | KX208744 | KX208890 | KX209735 | KX211301 | KX211735 | KX200457 |
| *Hoplobatrachus tigerinus* | Africa Asia India | KX200207 | KX201218 | KX201366 | KX201892 | KX202170 | KX202318 | KX204465 | KX207139 | KX208639 | KX208785 | KX208924 | KX209776 | KX211340 | KX211775 | KX200498 |
| *Hoplophryne rogersi* | Africa | | | | | | | | | | | EF396089 | | | | EF017980 |
| *Hyalinobatrachium aureoguttatum* | Seychelles | | | | | | | | | | | EU663491 | | | | KF534403 |
| *Hyalinobatrachium colymbiphyllum* | Seychelles | | | | | | | | | | | EU663498 | | | | KF534408 |
| *Hyalinobatrachium ibama* | Seychelles | | | | | | | | | | | EU663507 | | | | KF534413 |
| *Hydrophylax leptoglossa* | India | | | | | | | | | | | KR264388 | | | | KR264308 |
| *Hyla arenicolor* | Asia America Seychelles | | | | | | | | | | | AY364220 | | | | AY364190 |
| *Hyla chinensis* | Asia America Seychelles | KX200167 | KX201179 | KX201328 | KX201851 | KX202130 | KX202276 | KX204425 | KX207101 | KX208599 | KX208743 | KX208889 | KX209734 | KX211300 | KX211734 | KX200456 |
| *Hyla cinerea* | Asia America Seychelles | | | | | | | | | | | AY323766 | | | | DQ306493 |
| *Hylarana sp.* | India IAA | KX200119 | KX201125 | KX201275 | KX201797 | KX202078 | KX202226 | KX204372 | KX207052 | KX208546 | KX208691 | KX208837 | KX209680 | KX211245 | KX211679 | |
| *Hylodes nasus* | Seychelles | | | | | | | | | | | KJ961597 | | | | GQ345194 |
| *Hylophorbus rufescens* | Australia | | | | | | | | | | | EF018047 | | | | EF017977 |
| *Hyloscirtus lindae* | Seychelles | KX200109 | KX201114 | KX201264 | KX201787 | KX202067 | KX202215 | KX204361 | KX207041 | KX208535 | KX208680 | KX208827 | KX209669 | KX211234 | KX211669 | KX200392 |

| | | | | | | | | | | | | | | | | |
|---|---|---|---|---|---|---|---|---|---|---|---|---|---|---|---|---|
| *Hyloxalus jacobuspetersi* | Seychelles | | KX201135 | KX201285 | KX201807 | KX202087 | | KX204382 | | KX208556 | KX208701 | KX208846 | KX209690 | KX211255 | KX211689 | KX200413 |
| *Hymenochirus boettgeri* | Africa | KX200170 | KX201182 | KX201331 | KX201854 | KX202133 | KX202279 | KX204428 | KX207104 | KX208601 | KX208746 | KX208892 | KX209737 | KX211303 | KX211737 | KX200459 |
| *Hyperolius bolifambae* | Africa | KX200166 | KX201178 | KX201327 | KX201850 | KX202129 | KX202275 | KX204424 | KX207100 | KX208598 | KX208742 | KX208888 | KX209733 | KX211299 | KX211733 | KX200455 |
| *Hyperolius viridiflavus* | Africa | | | | | | | | | | | AY323769 | | | | AB612019 |
| *Hypodactylus brunneus* | Seychelles | KX200106 | KX201111 | KX201260 | KX201783 | KX202063 | KX202212 | KX204357 | KX207037 | KX208531 | KX208676 | KX208823 | KX209666 | KX211231 | KX211665 | KX200388 |
| *Hypsiboas fasciatus* | Seychelles | KX200098 | KX201103 | KX201249 | KX201773 | KX202056 | KX202202 | KX204348 | KX207028 | KX208521 | KX208668 | KX208815 | KX209658 | KX211221 | KX211656 | KX200378 |
| *Ikakogi tayrona* | Seychelles | | | | | | | | | | | EU663455 | | | | KF534427 |
| *Incilius luetkenii* | Seychelles | | | | | | | | | | | KJ609668 | | | | DQ306565 |
| *Incilius nebulifer* | Seychelles | KX200108 | KX201113 | KX201262 | KX201785 | KX202065 | KX202214 | KX204359 | KX207039 | KX208533 | KX208678 | KX208825 | KX209668 | KX211233 | KX211667 | KX200390 |
| *Indirana sp. A* | India | | | | | | | | | | | DQ347204 | | | | EF017984 |
| *Indirana sp. B* | India | | | | | | | | | | | DQ347215 | | | | EF017985 |
| *Ingerophrynus divergens* | Asia IAA | | | | | | | | | | | EU712818 | | | | FJ882701 |
| *Ingerophrynus galeatus* | Asia IAA | | | | | | | | | | | DQ158374 | | | | DQ306506 |
| *Insuetophrynus acarpicus* | Seychelles | KX200103 | KX201108 | KX201256 | KX201779 | KX202060 | KX202208 | KX204354 | KX207034 | KX208527 | KX208673 | KX208820 | KX209663 | KX211227 | KX211662 | KX200384 |
| *Ischnocnema guentheri* | Seychelles | | | | | | | | | | | KC468335 | | | | GQ345181 |
| *Kalophrynus interlineatus* | Asia IAA | | | | | | | | | | | AB611909 | | | | AB611914 |
| *Kalophrynus pleurostigma* | Asia IAA | KX200171 | KX201183 | KX201332 | KX201855 | KX202134 | KX202280 | KX204429 | KX207105 | KX208602 | KX208747 | | KX209738 | KX211304 | KX211738 | KX200460 |
| *Kaloula conjuncta* | Asia IAA | | | KX201263 | KX201786 | KX202066 | | KX204360 | KX207040 | KX208534 | KX208679 | KX208826 | | | KX211668 | KX200391 |
| *Kaloula pulchra* | Asia IAA | KX200172 | KX201184 | KX201333 | KX201856 | KX202135 | KX202281 | KX204430 | KX207106 | KX208603 | KX208748 | KX208893 | KX209739 | KX211305 | KX211739 | KX200461 |
| *Kurixalus odontotarsus* | Asia IAA | KX200195 | KX201205 | KX201355 | KX201879 | KX202158 | KX202305 | KX204453 | KX207127 | KX208626 | KX208772 | | KX209763 | KX211327 | KX211762 | KX200485 |
| *Lankanectes corrugatus* | India | | | | | | | | | | | AY571653 | | | | AY948773 |
| *Leiopelma hochstetteri* | Europe | KX200132 | KX201139 | KX201290 | KX201811 | KX202092 | KX202239 | KX204386 | KX207064 | KX208560 | KX208704 | KX208851 | KX209694 | KX211260 | KX211694 | KX200418 |
| *Lepidobatrachus laevis* | Seychelles | | | | | | | | | | | DQ679270 | | | | KP295530 |
| *Lepidobatrachus sp.* | Seychelles | KX200111 | KX201116 | KX201266 | | KX202069 | KX202217 | KX204363 | KX207043 | KX208537 | KX208682 | KX208829 | KX209671 | KX211236 | KX211671 | KX200394 |
| *Leptobrachium chapaense* | Asia IAA | KX200174 | KX201186 | KX201335 | KX201858 | KX202137 | KX202283 | KX204432 | KX207108 | KX208605 | KX208750 | KX208895 | KX209741 | KX211307 | KX211741 | KX200463 |
| *Leptodactylodon ovatus* | Africa | KX200176 | KX201188 | KX201337 | KX201860 | KX202139 | KX202285 | KX204434 | KX207110 | KX208606 | KX208752 | KX208897 | KX209743 | KX211309 | KX211743 | KX200465 |
| *Leptodactylus albilabris* | Seychelles | KX200140 | KX201150 | KX201300 | KX201822 | KX202103 | KX202249 | KX204396 | KX207075 | KX208571 | KX208715 | KX208861 | KX209705 | KX211271 | KX211705 | |
| *Leptodactylus latrans* | Seychelles | | | | | | | | | | | DQ158343 | | | | DQ306492 |
| *Leptodactylus melanonotus* | Seychelles | | | | | | | | | | | AY364224 | | | | AY364194 |
| *Leptolalax alpinus* | Asia IAA | KX200173 | KX201185 | KX201334 | KX201857 | KX202136 | KX202282 | KX204431 | KX207107 | KX208604 | KX208749 | KX208894 | KX209740 | KX211306 | KX211740 | KX200462 |
| *Leptopelis kivuensis* | Africa | | | | | | | | | | | KM047088 | | | | AY364181 |
| *Leptopelis parkeri* | Africa | | | KX201338 | KX201861 | KX202140 | KX202286 | | KX207111 | KX208607 | KX208753 | | KX209744 | KX211310 | | KX200466 |
| *Leptophryne borbonica* | IAA | | | | | | | | | | | EF107287 | | | | EF107450 |
| *Limnodynastes salmini* | Australia | KX200177 | KX201189 | KX201339 | KX201862 | KX202141 | KX202287 | KX204435 | KX207112 | KX208608 | KX208754 | | KX209745 | KX211311 | KX211744 | KX200467 |
| *Limnonectes fujianensis* | Asia IAA | KX200175 | KX201187 | KX201336 | KX201859 | KX202138 | KX202284 | KX204433 | KX207109 | | KX208751 | KX208896 | KX209742 | KX211308 | KX211742 | KX200464 |
| *Limnonectes laticeps* | Asia IAA | | | | | | | | | | | AB488960 | | | | AB277320 |
| *Limnonectes magnus* | Asia IAA | | | | | | | | | | | JF744593 | | | | EF017994 |
| *Liophryne schlaginhaufeni* | IAA | | | KX201247 | KX201771 | | KX202200 | KX204346 | | KX208519 | | KX208813 | | KX211219 | | KX200376 |
| *Lithodytes lineatus* | Seychelles | | KX201140 | | KX201812 | KX202093 | KX202240 | KX204387 | KX207065 | KX208561 | KX208705 | KX208852 | KX209695 | KX211261 | KX211695 | KX200419 |
| *Litoria caerulea* | Australia IAA | | | | | | | | | | | AY323767 | | | | GQ365988 |
| *Liurana xizangensis* | Asia | KX200213 | KX201224 | KX201372 | KX201898 | KX202176 | KX202324 | KX204471 | KX207145 | KX208645 | KX208791 | KX208930 | KX209782 | KX211346 | KX211781 | KX200504 |

| | | | | | | | | | | | | | | | | |
|---|---|---|---|---|---|---|---|---|---|---|---|---|---|---|---|---|
| *Mantella madagascariensis* | Madagascar | | | | | | | | | | | AB612043 | | | | AB612047 |
| *Mantophryne lateralis* | IAA | | | KX201245 | | KX202054 | KX202198 | | KX207025 | KX208517 | KX208665 | KX208811 | KX209656 | KX211217 | KX211653 | KX200374 |
| *Megophrys nasuta* | IAA | | | | | | | | | | | AB612063 | | | | AB612068 |
| *Melanobatrachus indicus* | India | | | | | | | | | | | EF018053 | | | | EF017983 |
| *Melanophryniscus stelzneri* | Seychelles | KX200086 | KX201090 | KX201233 | KX201757 | KX202042 | KX202185 | KX204334 | | KX208504 | KX208653 | KX208800 | KX209644 | KX211204 | KX211640 | KX200361 |
| *Meristogenys kinabaluensis* | IAA | | | | | | | | | | | EF088250 | | | | AY364176 |
| *Mertensophryne micranotis* | Africa | | | | | | | | | | | EF107330 | | | | EF107491 |
| *Metaphrynella pollicaris* | IAA | | | | | | | | | | | AB611927 | | | | AB611932 |
| *Metaphrynella sundana* | IAA | | | | | | | | | | | AB611935 | | | | EF017973 |
| *Micrixalus sp.* | India | | | | | | | | | | | DQ347216 | | | | EF017986 |
| *Microhyla annectens* | Asia India IAA | | | | | | | | | | | AB611940 | | | | AB611945 |
| *Microhyla heymonsi* | Asia India IAA | KX200179 | KX201191 | KX201341 | KX201864 | | KX202289 | KX204437 | KX207114 | KX208610 | KX208756 | KX208899 | KX209747 | KX211313 | KX211746 | KX200469 |
| *Microhyla marmorata* | Asia India IAA | | | | | | | | | | | AB611948 | | | | AB611953 |
| *Microhyla okinavensis* | Asia India IAA | | | | | | | | | | | AB611956 | | | | AB611960 |
| *Micryletta inornata* | Asia | | | | | | | | | | | AB611962 | | | | AB611966 |
| *Mixophyes coggeri* | Australia IAA | KX200178 | KX201190 | KX201340 | KX201863 | KX202142 | KX202288 | KX204436 | KX207113 | KX208609 | KX208755 | KX208898 | KX209746 | KX211312 | KX211745 | KX200468 |
| *Myobatrachus gouldii* | Australia | | | | | | | | | | | AY364226 | | | | AY364196 |
| *Nannophryne cophotis* | Seychelles | | | | | | | | | | | DQ158369 | | | | DQ306540 |
| *Nannophryne variegata* | Seychelles | | | | | | | | | | | DQ158410 | | | | DQ306515 |
| *Nasikabatrachus sahyadrensis* | India | | | | | | | | | | | AY364225 | | | | AY364195 |
| *Nectophrynoides tornieri* | Africa | | | | | | | | | | | EF107329 | | | | EF107490 |
| *Nyctibates corrugatus* | Africa | KX200181 | | KX201342 | KX201866 | KX202144 | KX202291 | KX204439 | KX207116 | KX208612 | KX208758 | KX208901 | KX209749 | KX211315 | KX211748 | KX200471 |
| *Nyctibatrachus sp.* | India | | | | | | | | | | | AY948913 | | | | |
| *Nyctimystes kubori* | IAA | KX200182 | KX201193 | KX201343 | KX201867 | KX202145 | KX202292 | KX204440 | KX207117 | KX208613 | KX208759 | KX208902 | KX209750 | KX211316 | KX211749 | KX200472 |
| *Nyctimystes pulcher* | IAA | | | | | | | | | | | AY948941 | | | | GQ365993 |
| *Nymphargus grandisonae* | Seychelles | | | | | | | | | | | EU663443 | | | | KF534431 |
| *Occidozyga laevis* | Asia IAA | | | | | | | | | | | DQ347254 | | | | EF017995 |
| *Occidozyga lima* | Asia IAA | | | | | | | | | | | AB488961 | | | | AB488920 |
| *Odontobatrachus natator* | Africa | KX200188 | KX201199 | KX201348 | KX201873 | KX202151 | KX202298 | KX204446 | KX207122 | KX208619 | KX208765 | KX208908 | KX209756 | KX211322 | KX211755 | KX200478 |
| *Odontophrynus occidentalis* | Seychelles | KX200117 | KX201123 | KX201273 | KX201795 | KX202076 | KX202224 | KX204370 | KX207050 | KX208544 | KX208689 | | KX209678 | KX211243 | KX211678 | KX200401 |
| *Odorrana hosii* | Asia | | | | | | | | | | | EF088259 | | | | KR264331 |
| *Odorrana schmackeri* | Asia | KX200206 | KX201217 | | KX201891 | KX202169 | KX202317 | KX204464 | KX207138 | KX208638 | KX208784 | KX208923 | KX209775 | KX211339 | KX211774 | KX200497 |
| *Ophryophryne microstoma* | Asia IAA | KX200184 | KX201195 | | KX201869 | KX202147 | KX202294 | KX204442 | KX207119 | KX208615 | KX208761 | KX208904 | KX209752 | KX211318 | KX211751 | KX200474 |
| *Oreolalax jingdongensis* | Asia | KX200183 | KX201194 | KX201344 | KX201868 | KX202146 | KX202293 | KX204441 | KX207118 | KX208614 | KX208760 | KX208903 | KX209751 | KX211317 | KX211750 | KX200473 |
| *Oreophryne sp.* | IAA | | KX201148 | KX201298 | KX201820 | KX202101 | KX202247 | KX204394 | KX207073 | KX208569 | KX208713 | KX208859 | KX209703 | KX211269 | KX211703 | KX200427 |
| *Oreophryne sp. B* | IAA | | | | | | | | | | | EF018046 | | | | EF017976 |
| *Osteocephalus taurinus* | Seychelles | KX200185 | KX201196 | KX201345 | KX201870 | KX202148 | KX202295 | KX204443 | KX207120 | KX208616 | KX208762 | KX208905 | KX209753 | KX211319 | KX211752 | KX200475 |
| *Otophryne pyburni* | Seychelles | KX200101 | KX201106 | KX201253 | KX201777 | | KX202206 | KX204351 | KX207032 | KX208525 | KX208671 | KX208818 | KX209661 | KX211224 | KX211659 | KX200381 |
| *Papurana latouchii* | Australia | KX200205 | KX201216 | | KX201890 | KX202168 | KX202316 | KX204463 | KX207137 | KX208637 | KX208783 | KX208922 | KX209774 | KX211338 | KX211773 | KX200496 |
| *Paradoxophyla palmata* | Madagascar | KX200112 | KX201117 | KX201267 | KX201789 | KX202070 | KX202218 | KX204364 | KX207044 | KX208538 | KX208683 | KX208830 | KX209672 | KX211237 | KX211672 | KX200395 |
| *Pelobates cultripes* | America | | | | | | | | | | | AY323758 | | | | AY364171 |

| | | | | | | | | | | | | | | | | |
|---|---|---|---|---|---|---|---|---|---|---|---|---|---|---|---|---|
| *Pelobates syriacus* | America | KX200124 | KX201130 | KX201280 | KX201802 | KX202082 | KX202231 | KX204377 | KX207057 | KX208551 | KX208696 | KX208841 | KX209685 | KX211250 | KX211684 | KX200408 |
| *Pelodytes ibericus* | America | KX200191 | KX201202 | KX201351 | KX201876 | KX202154 | KX202301 | KX204449 | KX207124 | KX208622 | KX208768 | KX208911 | KX209759 | KX211324 | KX211758 | KX200481 |
| *Pelophylax nigromaculatus** | Asia America | | | | | | | | | | | | | | | Dryad |
| *Peltophryne lemur* | Seychelles | | | | | | | | | | | DQ158386 | | | | DQ306513 |
| *Peltophryne longinasus* | Seychelles | | | | | | | | | | | JF342384 | | | | JF342426 |
| *Peltophryne peltocephala* | Seychelles | KX200133 | KX201141 | KX201291 | KX201813 | KX202094 | KX202241 | KX204388 | KX207066 | KX208562 | KX208706 | KX208853 | KX209696 | KX211262 | KX211696 | KX200420 |
| *Petropedetes euskircheni* | Africa | KX200189 | KX201200 | KX201349 | KX201874 | KX202152 | KX202299 | KX204447 | | KX208620 | KX208766 | KX208909 | KX209757 | | KX211756 | KX200479 |
| *Petropedetes parkeri* | Africa | | | | | | | | | | | DQ019505 | | | | AY364183 |
| *Phlyctimantis boulengeri* | Africa | KX200186 | KX201197 | KX201346 | KX201871 | KX202149 | KX202296 | KX204444 | | KX208617 | KX208763 | KX208906 | KX209754 | KX211320 | KX211753 | KX200476 |
| *Phrynella pulchra* | IAA | | | | | | | | | | | AB611969 | | | | AB611974 |
| *Phrynobatrachus africanus* | Africa | | | | | | | | | | | KF693602 | | | | EF017996 |
| *Phrynobatrachus krefftii* | Africa | | | | | | | | | | | GU457687 | | | | EF017997 |
| *Phrynobatrachus natalensis* | Africa | KX200190 | KX201201 | KX201350 | KX201875 | KX202153 | KX202300 | KX204448 | KX207123 | KX208621 | KX208767 | KX208910 | KX209758 | KX211323 | KX211757 | KX200480 |
| *Phrynoidis aspera* | Asia IAA | | | | | | | | | | | DQ158356 | | | | DQ306503 |
| *Phrynoidis juxtaspera* | Asia IAA | | | | | | | | | | | DQ158385 | | | | DQ306542 |
| *Phrynomantis microps* | Africa | KX200194 | KX201204 | KX201354 | KX201878 | KX202157 | KX202304 | KX204452 | KX207126 | KX208625 | KX208771 | KX208913 | KX209762 | KX211326 | KX211761 | KX200484 |
| *Phrynopus bracki* | Seychelles | | | | | | | | | | | GQ345281 | | | | GQ345186 |
| *Phyllobates vittatus* | Seychelles | | | | | | | | | | | EF107296 | | | | EF107459 |
| *Phyllomedusa hypochondrialis* | Seychelles | | | | | | | | | | | AY948929 | | | | GQ366014 |
| *Phyllomedusa tomopterna* | Seychelles | | KX201210 | KX201360 | KX201884 | KX202163 | KX202310 | KX204458 | | KX208631 | KX208777 | | KX209768 | KX211332 | KX211767 | KX200490 |
| *Physalaemus cuvieri* | Seychelles | KX200100 | KX201105 | KX201251 | KX201775 | KX202058 | KX202204 | KX204350 | KX207030 | KX208523 | KX208670 | KX208817 | KX209660 | KX211223 | KX211658 | KX200380 |
| *Physalaemus pustulosus* | Seychelles | KX200088 | KX201092 | KX201235 | KX201759 | KX202044 | KX202187 | KX204336 | KX207017 | KX208506 | KX208655 | KX208802 | KX209646 | KX211206 | KX211642 | KX200363 |
| *Pipa parva* | Seychelles | KX200131 | KX201138 | KX201289 | KX201810 | KX202091 | KX202238 | KX204385 | | KX208559 | KX208703 | KX208850 | KX209693 | KX211259 | KX211693 | KX200417 |
| *Pipa pipa* | Seychelles | KX200144 | KX201154 | KX201304 | KX201826 | KX202107 | KX202253 | KX204400 | KX207078 | KX208575 | KX208719 | KX208865 | KX209709 | KX211275 | KX211709 | KX200432 |
| *Platymantis hazelae* | IAA | | | | | | | | | | | DQ347248 | | | | EF017993 |
| *Platypelis tuberifera* | Madagascar | | | KX201254 | | | | KX204352 | | | | | | KX211225 | KX211660 | KX200382 |
| *Plethodontohyla inguinalis* | Madagascar | | | | | | | | | | | AB611985 | | | | AB611990 |
| *Pleurodema somuncurensis* | Seychelles | KX200091 | KX201096 | KX201238 | KX201763 | KX202048 | KX202191 | KX204340 | KX207021 | KX208510 | KX208659 | KX208806 | KX209650 | KX211210 | KX211646 | KX200367 |
| *Pleurodema thaul* | Seychelles | KX200199 | KX201209 | KX201359 | KX201883 | KX202162 | KX202309 | KX204457 | KX207131 | KX208630 | KX208776 | | KX209767 | KX211331 | KX211766 | KX200489 |
| *Polypedates megacephalus* | Asia IAA India | KX200192 | KX201203 | KX201352 | KX201877 | KX202155 | KX202302 | KX204450 | KX207125 | KX208623 | KX208769 | KX208912 | KX209760 | KX211325 | KX211759 | KX200482 |
| *Pristimantis thymelensis* | Seychelles | KX200162 | KX201173 | KX201322 | KX201845 | KX202124 | KX202270 | KX204419 | KX207095 | KX208593 | KX208737 | KX208883 | KX209728 | KX211294 | KX211728 | KX200450 |
| *Probreviceps durirostris* | Africa | | | | | | | | | | | KF990045 | | | | KF954752 |
| *Proceratophrys boiei* | Seychelles | KX200125 | KX201131 | KX201281 | KX201803 | KX202083 | KX202232 | KX204378 | KX207058 | KX208552 | KX208697 | KX208842 | KX209686 | KX211251 | KX211685 | KX200409 |
| *Pseudhymenochirus merlini* | Africa | KX200193 | | KX201353 | | KX202156 | KX202303 | KX204451 | | KX208624 | KX208770 | | KX209761 | | KX211760 | KX200483 |
| *Pseudis paradoxa* | Seychelles | KX200127 | KX201133 | KX201283 | KX201805 | KX202085 | KX202234 | KX204380 | KX207060 | KX208554 | KX208699 | KX208844 | KX209688 | KX211253 | KX211687 | KX200411 |
| *Pseudophilautus wynaadensis* | India | | | | | | | | | | | GQ204568 | | | | AY364169 |
| *Ptychadena cooperi* | Africa Madagascar New Zealand | | | | | | | | | | | KF380597 | | | | KF380104 |
| *Ptychadena mascareniensis* | Africa Madagascar New Zealand | | | | | | | | | | | AY571658 | | | | EF017992 |
| *Ptychadena oxyrhynchus* | Africa Madagascar New Zealand | KX200196 | KX201206 | KX201356 | KX201880 | KX202159 | KX202306 | KX204454 | KX207128 | KX208627 | KX208773 | KX208914 | KX209764 | KX211328 | KX211763 | KX200486 |
| *Pyxicephalus edulis* | Africa | | | | | | | | | | | KF991338 | | | | EF107494 |
| *Quasipaa spinosa* | Asia | KX200198 | KX201208 | KX201358 | KX201882 | KX202161 | KX202308 | KX204456 | KX207130 | KX208629 | KX208775 | KX208916 | KX209766 | KX211330 | KX211765 | KX200488 |

| | | | | | | | | | | | | | | | | |
|---|---|---|---|---|---|---|---|---|---|---|---|---|---|---|---|---|
| *Rana amurensis* | Asia America | KX200200 | KX201211 | KX201361 | KX201885 | | KX202311 | KX204459 | KX207132 | KX208632 | KX208778 | KX208917 | KX209769 | KX211333 | KX211768 | KX200491 |
| *Rana berlandieri* | Seychelles | KX200102 | KX201107 | KX201255 | KX201778 | KX202059 | KX202207 | KX204353 | KX207033 | KX208526 | KX208672 | KX208819 | KX209662 | KX211226 | KX211661 | KX200383 |
| *Rana catesbeiana* | Seychelles | | | | | | | | | | | AB612037 | | | | AB612041 |
| *Rana chensinensis* | Asia | KX200202 | KX201213 | KX201363 | KX201887 | KX202165 | KX202313 | KX204461 | KX207134 | KX208634 | KX208780 | KX208919 | KX209771 | KX211335 | KX211770 | KX200493 |
| *Rana draytonii* | Seychelles | KX200137 | KX201146 | KX201296 | KX201818 | KX202099 | KX202245 | KX204392 | KX207071 | KX208567 | KX208711 | KX208857 | KX209701 | KX211267 | KX211701 | KX200425 |
| *Rana japonica* | Asia | | | | | | | | | | | AB728272 | | | | KR264277 |
| *Rana pipiens* | Seychelles | | | | | | | | | | | JN227250 | | | | JN227141 |
| *Rana temporaria* | America | | | | | | | | | | | AY323776 | | | | EF017988 |
| *Rana virgatipes* | Seychelles | KX200135 | KX201144 | KX201294 | KX201816 | KX202097 | KX202243 | KX204390 | KX207069 | KX208565 | KX208709 | KX208855 | KX209699 | KX211265 | KX211699 | KX200423 |
| *Ranitomeya imitator* | Seychelles | KX200083 | KX201087 | KX201230 | KX201754 | KX202039 | KX202182 | KX204332 | | KX208501 | KX208650 | KX208797 | KX209641 | KX211201 | KX211636 | KX200358 |
| *Rentapia hosii* | IAA | KX200084 | KX201088 | KX201231 | KX201755 | KX202040 | KX202183 | | KX207014 | KX208502 | KX208651 | KX208798 | KX209642 | KX211202 | KX211637 | KX200359 |
| *Rhacophorus dennysi* | Asia IAA India | KX200187 | KX201198 | KX201347 | KX201872 | KX202150 | KX202297 | KX204445 | KX207121 | KX208618 | KX208764 | KX208907 | KX209755 | KX211321 | KX211754 | KX200477 |
| *Rhaebo glaberrimus* | Seychelles | | | | | | | | | | | DQ158377 | | | | DQ306548 |
| *Rhaebo nasicus* | Seychelles | | | | | | | | | | | DQ158396 | | | | DQ306512 |
| *Rhinella marina* | Seychelles | KX200107 | KX201112 | KX201261 | KX201784 | KX202064 | KX202213 | KX204358 | KX207038 | KX208532 | KX208677 | KX208824 | KX209667 | KX211232 | KX211666 | KX200389 |
| *Rhinella ocellata* | Seychelles | | | | | | | | | | | JN867519 | | | | DQ306538 |
| *Rhinoderma darwinii* | Seychelles | KX200092 | KX201097 | KX201239 | KX201764 | KX202049 | KX202192 | KX204341 | KX207022 | KX208511 | KX208660 | KX208807 | KX209651 | KX211211 | KX211647 | KX200368 |
| *Rhinophrynus dorsalis* | Seychelles | KX200203 | KX201214 | KX201364 | KX201888 | KX202166 | KX202314 | | KX207135 | KX208635 | KX208781 | KX208920 | KX209772 | KX211336 | KX211771 | KX200494 |
| *Rulyrana adiazeta* | Seychelles | | | | | | | | | | | EU663460 | | | | KF534442 |
| *Rulyrana flavopunctata* | Seychelles | | | | | | | | | | | EU663467 | | | | KF534443 |
| *Sachatamia ilex* | Seychelles | | | | | | | | | | | EU663446 | | | | KF534447 |
| *Sanguirana luzonensis* | IAA | | | | | | | | | | | EU076755 | | | | KR264350 |
| *Scaphiophryne boribory* | Madagascar | KX200115 | KX201121 | KX201271 | KX201793 | KX202074 | KX202222 | KX204368 | KX207048 | KX208542 | KX208687 | KX208834 | KX209676 | KX211241 | KX211676 | KX200399 |
| *Scaphiophryne madagascariensis* | Madagascar | | | | | | | | | | | AB612001 | | | | AB612006 |
| *Scaphiophryne marmorata* | Madagascar | KX200114 | KX201119 | KX201269 | KX201791 | KX202072 | KX202220 | KX204366 | KX207046 | KX208540 | KX208685 | KX208832 | KX209674 | KX211239 | KX211674 | KX200397 |
| *Scaphiopus couchii* | Seychelles | KX200208 | KX201219 | KX201367 | KX201893 | KX202171 | KX202319 | KX204466 | KX207140 | KX208640 | KX208786 | KX208925 | KX209777 | KX211341 | KX211776 | KX200499 |
| *Scaphiopus holbrookii* | Seychelles | | | | | | | | | | | AB612071 | | | | AB612076 |
| *Schismaderma carens* | Africa | KX200096 | KX201100 | KX201243 | KX201768 | KX202052 | KX202196 | KX204344 | KX207024 | KX208515 | KX208663 | KX208810 | KX209654 | KX211215 | KX211651 | KX200372 |
| *Scinax ruber* | Seychelles | KX200087 | KX201091 | KX201234 | KX201758 | KX202043 | KX202186 | KX204335 | KX207016 | KX208505 | KX208654 | KX208801 | KX209645 | KX211205 | KX211641 | KX200362 |
| *Sclerophrys brauni* | Africa | | | | | | | | | | | DQ158361 | | | | DQ306514 |
| *Sclerophrys maculata* | Africa | | | | | | | | | | | KJ609678 | | | | DQ306533 |
| *Scotobleps gabonicus* | Africa | KX200209 | KX201220 | KX201368 | KX201894 | KX202172 | KX202320 | KX204467 | KX207141 | KX208641 | KX208787 | KX208926 | KX209778 | KX211342 | KX211777 | KX200500 |
| *Scutiger gongshanensis* | Asia | KX200210 | KX201221 | KX201369 | KX201895 | KX202173 | KX202321 | KX204468 | KX207142 | KX208642 | KX208788 | KX208927 | KX209779 | KX211343 | KX211778 | KX200501 |
| *Sooglossus thomasseti* | New Zealand | KX200123 | KX201129 | KX201279 | KX201801 | | KX202230 | KX204376 | KX207056 | KX208550 | KX208695 | KX208840 | KX209684 | KX211249 | KX211683 | KX200407 |
| *Spea intermontana* | Seychelles | KX200121 | KX201127 | KX201277 | KX201799 | KX202080 | KX202228 | KX204374 | KX207054 | KX208548 | KX208693 | KX208839 | KX209682 | KX211247 | KX211681 | KX200405 |
| *Spea multiplicata* | Seychelles | KX200212 | KX201223 | KX201371 | KX201897 | KX202175 | KX202323 | KX204470 | KX207144 | KX208644 | KX208790 | KX208929 | KX209781 | KX211345 | KX211780 | KX200503 |
| *Spelaeophryne methneri* | Africa | | | | | | | | | | | KF990042 | | | | EF107453 |
| *Staurois latopalmatus* | IAA | | | | | | | | | | | AB612049 | | | | EF017987 |
| *Stefania ginesi* | Seychelles | | | | | | | | | | | DQ679308 | | | | GQ345178 |
| *Stereocyclops incrassatus* | Seychelles | | | KX201287 | | KX202089 | | | | | | KX208848 | | KX211257 | KX211691 | KX200415 |
| *Strabomantis biporcatus* | Seychelles | | | | | | | | | | | GQ345283 | | | | GQ345188 |

| | | | | | | | | | | | | | | | | |
|---|---|---|---|---|---|---|---|---|---|---|---|---|---|---|---|---|
| *Strabomantis sulcatus* | Seychelles | KX200082 | KX201086 | KX201229 | KX201753 | KX202038 | KX202181 | KX204331 | KX207013 | KX208500 | KX208649 | KX208796 | KX209640 | KX211200 | KX211635 | KX200357 |
| *Strongylopus grayii* | Africa | KX200211 | KX201222 | KX201370 | KX201896 | KX202174 | KX202322 | KX204469 | KX207143 | KX208643 | KX208789 | KX208928 | KX209780 | KX211344 | KX211779 | KX200502 |
| *Stumpffia pygmaea* | Madagascar | | | KX201257 | KX201780 | | KX202209 | | | KX208528 | | KX208821 | | KX211228 | | KX200385 |
| *Sylvirana guentheri* | Asia | KX200204 | KX201215 | KX201365 | KX201889 | KX202167 | KX202315 | KX204462 | KX207136 | KX208636 | KX208782 | KX208921 | KX209773 | KX211337 | KX211772 | KX200495 |
| *Synapturanus sp.* | Seychelles | | | | | | | | | | | EF018051 | | | | EF017981 |
| *Telmatobius vellardi* | Seychelles | KX200215 | KX201226 | KX201374 | KX201900 | KX202178 | KX202326 | KX204473 | KX207147 | KX208646 | KX208793 | KX208932 | KX209784 | KX211348 | KX211783 | KX200506 |
| *Teratohyla spinosa* | Seychelles | | | | | | | | | | | EU663483 | | | | KF534452 |
| *Thoropa taophora* | Seychelles | | | | | | | | | | | GQ345288 | | | | GQ345193 |
| *Trachycephalus typhonius* | Seychelles | | | | | | | | | | | EU034147 | | | | AY364185 |
| *Trichobatrachus robustus* | Africa | KX200214 | KX201225 | KX201373 | KX201899 | KX202177 | KX202325 | KX204472 | KX207146 | | KX208792 | KX208931 | KX209783 | KX211347 | KX211782 | KX200505 |
| *Uperodon montanus* | India | | | | | | | | | | | AB611993 | | | | AB611998 |
| *Uperodon systoma* | India | | | | | | | | | | | EF018049 | | | | EF017979 |
| *Uperodon variegatus* | India | | | | | | | | | | | EF018052 | | | | EF017982 |
| *Uperoleia laevigata* | Australia IAA | | | | | | | | | | | KJ874616 | | | | EF107474 |
| *Vitreorana helenae* | Seychelles | | | | | | | | | | | EU663471 | | | | KF534456 |
| *Xenophrys omeimontis* | Asia IAA | KX200180 | KX201192 | | KX201865 | KX202143 | KX202290 | KX204438 | KX207115 | KX208611 | KX208757 | KX208900 | KX209748 | KX211314 | KX211747 | KX200470 |
| *Xenopus epitropicalis* | Africa | KX200216 | KX201227 | KX201375 | KX201901 | KX202179 | KX202327 | KX204474 | KX207148 | KX208647 | KX208794 | KX208933 | KX209785 | KX211349 | KX211784 | KX200507 |
| *Xenopus kobeli* | Africa | KX200217 | KX201228 | | KX201902 | KX202180 | KX202328 | KX204475 | KX207149 | KX208648 | KX208795 | KX208934 | KX209786 | KX211350 | | KX200508 |
| *Xenorhina obesa* | IAA | | | | | | | | | | | EF018048 | | | | EF017978 |
| *Xenorhina sp.* | IAA | KX200129 | KX201136 | KX201286 | KX201808 | KX202088 | KX202236 | KX204383 | KX207062 | KX208557 | KX208702 | KX208847 | KX209691 | KX211256 | KX211690 | KX200414 |

* See original article for data on these taxa and for data on the outgroups

The data used in these analyses were taken from Feng et al. (2017), who used multiple sources for a large amount of data. In order to reduce the dataset, we selected the 20 most parsimony informative markers and, from those, we eliminated the ones with the least complete dataset, which reduced our selection to 14 markers: CILP, DISP2, DMXL1, DSEL, EVPL, EXOC8, HYP, MSH6, PPL, RAG1, RAG2, SALL1, WFIKKN2, ZFPM2. Since this was inbetween a genus-level and a family-level phylogeny, we attributed distributions according to the range of the entire genus. In addition, we extrapolated the dates from the original articles with soft lower boundaries, because we reduced the amount of data used, even though other articles have retrieved the same topology (e.g. Li et al., 2013) with congruent dates.

#### 9. Emballonurinae

Table S8: Genbank numbers for the sequences used in the Emballonuridae analyses with the distribution attributed to each terminal.

| Species | Locality | CytB | RAG2 |
|---|---|---|---|
| *Emballonura tiavato* | Madagascar | DQ178278 | |
| *Emballonura tiavato* | Madagascar | DQ178275 | |
| *Emballonura tiavato* | Madagascar | DQ178280 | HQ693737 |
| *Emballonura tiavato* | Madagascar | DQ178277 | |
| *Emballonura tiavato* | Madagascar | DQ178252 | |
| *Emballonura tiavato* | Madagascar | DQ178255 | HQ693738 |
| *Emballonura tiavato* | Madagascar | DQ178253 | |
| *Emballonura tiavato* | Madagascar | DQ178250 | |
| *Emballonura tiavato* | Madagascar | DQ178273 | |
| *Emballonura tiavato* | Madagascar | DQ178262 | |
| *Emballonura tiavato* | Madagascar | DQ178264 | |
| *Emballonura tiavato* | Madagascar | DQ178265 | |
| *Emballonura tiavato* | Madagascar | DQ178249 | |
| *Emballonura tiavato* | Madagascar | DQ178272 | |
| *Emballonura atrata* | Madagascar | HQ693729 | |
| *Emballonura atrata* | Madagascar | HQ693732 | |
| *Emballonura atrata* | Madagascar | HQ693733 | |
| *Emballonura atrata* | Madagascar | HQ693730 | |
| *Emballonura atrata* | Madagascar | HQ693726 | |
| *Emballonura atrata* | Madagascar | HQ693734 | HQ693745 |
| *Emballonura atrata* | Madagascar | HQ693727 | HQ693739 |
| *Emballonura atrata* | Madagascar | HQ693728 | |
| *Emballonura atrata* | Madagascar | HQ693725 | |

| Species | Locality | CytB | RAG2 |
|---|---|---|---|
| *Emballonura atrata* | Madagascar | HQ693731 | |
| *Emballonura monticola* | IAA | HQ693714 | HQ693747 |
| *Emballonura monticola* | IAA | HQ693715 | HQ693746 |
| *Emballonura semicaudata* | IAA | HQ693708 | HQ693751 |
| *Emballonura semicaudata* | IAA | HQ693709 | HQ693751 |
| *Emballonura alecto* | IAA | HQ693710 | HQ693752 |
| *Emballonura alecto* | IAA | HQ693711 | HQ693748 |
| *Emballonura dianae* | Australia | HQ693716 | HQ693750 |
| *Emballonura dianae* | Australia | HQ693717 | HQ693750 |
| *Emballonura raffrayana* | Australia | EF584224 | |
| *Emballonura beccarii* | Australia | EF584222 | |
| *Emballonura beccarii* | Australia | EF635537 | |
| *Emballonura serii* | Australia | EF635544 | |
| *Emballonura serii* | Australia | EF635543 | |
| *Mosia nigrescens* | Australia | EF635559 | |
| *Coleura afra* | Africa | HQ693720 | HQ693743 |
| *Coleura seychellensis* | Seychelles | HQ693735 | |
| *Peropteryx kappleri* | America | HQ693719 | HQ693741 |
| *Saccopteryx leptura* | America | HQ693718 | HQ693740 |
| *Cormura brevirostris* | America | EF584158 | HQ693742 |
| *Taphozous nudiventris* | Iran | HQ693712 | HQ693749 |
| *Nycteris hispida* | Africa | HQ693722 | HQ693736 |

#### 10. Strepsirrhini

The sequences used in the Strepsirrhini analyses were downloaded directly from the original article's dryad repository (https://datadryad.org/resource/doi:10.5061/dryad.51f00) and re-aligned. All calibration points were set to lognormal with parameters that we set to retrieve the dates from the original fossil descriptions: *Aegyptopithecus* (M = 1.0 and S = 0.6; offset = 29.9 m.y.a. (Seiffert, 2006; Simons, 1965)); *Branisella* (M = 1.0 and S = 0.59; offset = 26.4 m.y.a. (Hoffstetter, 1969; Takai, Anaya, Shigehara, & Setoguchi, 2000)); *Djebelemur* (M = 1.0 and S = 0.86; offset = 41.3 m.y.a. (Godinot, 2006; Hartenberger & Marandat, 1992)); *Nycticeboides* (M = 1.0 and S = 0.83; offset = 6.4 m.y.a. (MacPhee & Jacobs, 1986)); *Plesiopithecus* (M = 1.0 and S = 0.84; offset = 33.9 m.y.a. (E. L. Simons, 1992)); *Saharagalago* (M = 1.0 and S = 1.07; offset = 33.9 m.y.a. (Seiffert, Simons, & Attia, 2003)); and *Wadilemur* (M = 1.0 and S = 0.84; offset = 33.9 m.y.a. (Seiffert, Simons, Ryan, & Attia, 2005; Simons, 1997)). The dates we obtained are older than in Herrera and Dávalos (2016), but the topology was identical except for the position of

*Phaner furcifer*, which in our analyses was in agreement with Masters, Silvestro, Génin, and DelPero (2013).

#### 11. Pteropodidae

Table S9: Genbank numbers for the sequences used in the Pteropodidae analyses with the distribution attributed to each terminal.

| Species | Locality | CytB | 12S | 16S | RAG1 | RAG2 | vWF | BRCA1 |
|---|---|---|---|---|---|---|---|---|
| Acerodon celebensis | IAA | GQ410231 | JN398167 | AF293641 | EU617946 | EU617896 | EU617928 | JN398239 |
| Aethalops alecto | IAA | GQ410218 | GQ410312 | GQ410335 | GQ410263 | GQ410240 | GQ410286 | JN398260 |
| Aethalops alecto | IAA | GQ410219 | GQ410313 | GQ410336 | GQ410264 | GQ410241 | GQ410287 | |
| Alionycteris paucidentata | IAA | GQ410221 | GQ410315 | GQ410338 | GQ410266 | GQ410243 | GQ410289 | |
| Alionycteris paucidentata | IAA | GQ410222 | GQ410316 | GQ410339 | GQ410267 | GQ410244 | GQ410290 | JN398248 |
| Aproteles bulmerae | Australia | | U93066 | AF293645 | | | | |
| Balionycteris maculata | IAA | GQ410227 | GQ410321 | GQ410344 | GQ410272 | GQ410249 | GQ410295 | JN398257 |
| Boneia bidens | IAA | FJ218481 | FJ218475 | FJ218475 | FJ218468 | FJ218464 | FJ218471 | |
| Casinycteris argynnis | Africa | JN398197 | JN398168 | JN398168 | JN398284 | JN398301 | JN398268 | JN398264 |
| Chironax melanocephalus | IAA | GQ410220 | GQ410314 | | GQ410265 | GQ410242 | GQ410288 | JN398256 |
| Cynopterus brachyotis | IAA | AB046321 | U93068 | U93068 | | | | |
| Cynopterus brachyotis | IAA | GQ410210 | GQ410303 | GQ410327 | GQ410256 | GQ410233 | GQ410279 | JN398258 |
| Cynopterus sphinx | IAA | DQ445703 | GQ410302 | GQ410326 | EU617947 | EU617897 | DQ445697 | KT875821 |
| Desmalopex leucopterus | IAA | JN398198 | JN398169 | JN398169 | EU617966 | EU617915 | EU617929 | JN398252 |
| Dobsonia inermis | Australia | DQ445704 | FJ218476 | FJ218476 | EU617948 | EU617898 | DQ445686 | JN398234 |
| Dobsonia minor | Australia | DQ445705 | FJ218477 | FJ218477 | FJ218467 | FJ218463 | DQ445701 | |
| Dobsonia moluccensis | Australia | FJ218484 | JN398196 | JN398196 | EU617949 | EU617899 | EU617930 | JN398220 |
| Dobsonia praedatrix | Australia | JN398199 | JN398170 | JN398170 | JN398285 | JN398302 | JN398269 | JN398222 |
| Dyacopterus spadiceus | IAA | GQ410230 | GQ410324 | GQ410347 | GQ410275 | GQ410252 | GQ410298 | JN398259 |
| Eidolon helvum | Africa | JN398200 | JN398171 | JN398171 | EU617950 | EU617900 | EU617931 | JN398240 |
| Eonycteris robusta | IAA | JN398201 | JN398172 | JN398172 | JN398286 | JN398303 | JN398270 | JN398251 |
| Eonycteris spelaea | IAA | AB062476 | U93059 | | | | | |
| Eonycteris spelaea | IAA | FJ218482 | FJ529123 | FJ529123 | EU617951 | EU617901 | DQ445684 | JN398254 |
| Epomophorus wahlbergi | Africa | DQ445706 | JN398174 | JN398174 | EU617953 | EU617903 | DQ445691 | JN398266 |
| Epomops franqueti | Africa | DQ445707 | KT875876 | KT875876 | EU617952 | EU617902 | DQ445692 | |
| Epomops franqueti | Africa | JN398202 | JN398175 | JN398175 | JN398287 | JN398304 | JN398271 | JN398233 |
| Haplonycteris fischeri | IAA | GQ410225 | GQ410319 | GQ410342 | GQ410270 | GQ410247 | GQ410293 | JN398245 |
| Haplonycteris fischeri | IAA | GQ410226 | GQ410320 | GQ410343 | GQ410271 | GQ410248 | GQ410294 | |
| Harpyionycteris celebensis | IAA | JN398203 | FJ218473 | FJ218473 | JN398288 | | | JN398225 |
| Harpyionycteris whiteheadi | IAA | DQ445708 | FJ218474 | FJ218474 | EU617954 | EU617904 | DQ445690 | JN398246 |
| Hypsignathus monstrosus | Africa | JN398204 | JN398176 | JN398176 | JN398289 | JN398305 | JN398272 | JN398235 |
| Latidens salimalii | India | GQ410217 | GQ410311 | FJ009216 | | | | |
| Lissonycteris angolensis | Africa | JN398205 | JN398177 | JN398177 | JN398290 | JN398306 | JN398273 | JN398243 |
| Macroglossus minimus | Australia | JN398206 | JN398178 | JN398178 | EU617955 | EU617905 | DQ445693 | JN398229 |
| Macroglossus sobrinus | IAA | | JN398179 | JN398179 | JN398291 | JN398307 | JN398274 | JN398236 |

| | | | | | | | | |
|---|---|---|---|---|---|---|---|---|
| Megaerops ecaudatus | IAA | GQ410214 | GQ410308 | GQ410332 | GQ410260 | GQ410237 | GQ410283 | JN398262 |
| Megaerops kusnotoi | IAA | GQ410215 | GQ410309 | GQ410333 | GQ410261 | GQ410238 | GQ410284 | JN398255 |
| Megaerops wetmorei | IAA | GQ410212 | GQ410306 | GQ410330 | GQ410258 | GQ410235 | GQ410281 | |
| Megaerops wetmorei | IAA | GQ410213 | GQ410307 | GQ410331 | GQ410259 | GQ410232 | GQ410282 | |
| Megaloglossus woermanni | Africa | DQ445710 | JN398180 | JN398180 | EU617956 | EU617906 | DQ445702 | JN398231 |
| Melonycteris fardoulisi | Australia | FJ218478 | JN398181 | JN398181 | EU617957 | EU617907 | DQ445699 | JN398237 |
| Melonycteris melanops | Australia | JN398207 | JN398182 | JN398182 | FJ218465 | FJ218461 | FJ218469 | JN398223 |
| Micropteropus pusillus | Africa | JN398208 | JN398183 | JN398183 | JN398292 | JN398308 | JN398275 | JN398241 |
| Myonycteris torquata | Africa | FJ218483 | JN398184 | JN398184 | EU617958 | EU617908 | DQ445700 | JN398232 |
| Nanonycteris veldkampii | Africa | JN398209 | JN398185 | JN398185 | JN398293 | JN398309 | JN398276 | JN398253 |
| Nyctimene cephalotes | IAA | JN398210 | JN398186 | JN398186 | JN398294 | | JN398277 | JN398226 |
| Nyctimene robinsoni | Australia | AF144066 | GQ410325 | GQ410348 | GQ410276 | GQ410253 | GQ410299 | JN398221 |
| Nyctimene vizcaccia | Australia | DQ445711 | JN398187 | JN398187 | EU617959 | EU617904 | DQ445698 | JN398238 |
| Otopteropus cartilagonodus | IAA | GQ410223 | GQ410317 | GQ410340 | GQ410268 | GQ410245 | GQ410291 | |
| Otopteropus cartilagonodus | IAA | GQ410224 | GQ410318 | GQ410341 | GQ410269 | GQ410246 | GQ410292 | JN398249 |
| Penthetor lucasi | IAA | GQ410216 | GQ410310 | GQ410334 | GQ410262 | GQ410239 | GQ410285 | JN398261 |
| Ptenochirus jagori | IAA | FJ218480 | GQ410304 | GQ410328 | EU617960 | EU617910 | DQ445696 | JN398250 |
| Ptenochirus jagori | IAA | GQ410211 | GQ410305 | GQ410329 | GQ410257 | GQ410234 | GQ410280 | |
| Pteralopex atrata | Australia | | U93069 | AF293643 | | | | |
| Pteralopex acrodonta | Pacific | FJ561376 | FJ588908 | | | | | |
| Pteropus admiralitatum | Australia | KJ532441 | U93072 | | | | | |
| Pteropus aldabrensis | Seychelles | FJ561394 | FJ588896 | | | | | |
| Pteropus conspicillatus | Australia | FJ561378 | FJ588879 | | EU617963 | | | |
| Pteropus conspicillatus | Australia | FJ561379 | FJ588880 | | | | | |
| Pteropus conspicillatus | Australia | FJ561380 | FJ588881 | | | | | |
| Pteropus dasymallus | IAA | AB042770 | AB042770 | AB042770 | | | | |
| Pteropus giganteus | India | FJ561381 | FJ588882 | | JN414929 | AY011964 | | |
| Pteropus giganteus | India | JN398211 | AY012138 | AY011170 | EU617964 | EU617913 | EU617935 | JN398242 |
| Pteropus hypomelanus | IAA | FJ561382 | FJ588883 | AF044623 | EU617965 | EU617914 | AF203777 | |
| Pteropus hypomelanus | IAA | FJ561383 | FJ588884 | AF069537 | | AY141025 | DQ445687 | |
| Pteropus livingstonii | Comoros | FJ561384 | FJ588885 | | | | | |
| Pteropus niger | MI | FJ561385 | FJ588886 | | | | | |
| Pteropus poliocephalus | Australia | FJ561386 | FJ588887 | | EU617971 | EU617920 | EU617940 | |
| Pteropus poliocephalus | Australia | FJ561387 | FJ588888 | | | | | |
| Pteropus pumilus | IAA | FJ561388 | FJ588889 | | EU617972 | EU617921 | EU617941 | |
| Pteropus pumilus | IAA | FJ561389 | FJ588890 | | | | | |
| Pteropus pumilus | IAA | FJ561390 | FJ588891 | | | | | |
| Pteropus rodricensis | MI | FJ561391 | FJ588892 | AF044624 | | | | |
| Pteropus rodricensis | MI | FJ561392 | FJ588893 | | | | | |
| Pteropus rufus | Madagascar | FJ561393 | FJ588894 | | | | | |
| Pteropus scapulatus | Australia | AF321050 | AF321050 | AF321050 | EU617974 | | | |
| Pteropus scapulatus | Australia | FJ561377 | FJ588895 | | | | | |
| Pteropus seychellensis comoroensis | Comoros. | FJ561395 | FJ588897 | | | | | |

| | | | | | | | | |
|---|---|---|---|---|---|---|---|---|
| Pteropus seychellensis comoroensis | Comoros | FJ561396 | FJ588898 | | | | | |
| Pteropus seychellensis comoroensis | Comoros | FJ561397 | FJ588899 | | | | | |
| Pteropus seychellensis comoroensis | Comoros | FJ561398 | FJ588900 | | | | | |
| Pteropus seychellensis seychellensis | Seychelles | FJ561399 | FJ588901 | | | | | |
| Pteropus seychellensis seychellensis | Seychelles | FJ561400 | FJ588902 | | | | | |
| Pteropus speciosus | IAA | AB062474 | | | | | | |
| Pteropus tonganus | Pacific | JN398213 | JN398188 | JN398188 | EU617976 | EU617924 | DQ445695 | JN398267 |
| Pteropus vampyrus | IAA | EF584230 | KJ532340 | | EU617977 | EU617925 | EU617944 | |
| Pteropus vampyrus | IAA | FJ561402 | FJ588904 | NC_026542 | | | | |
| Pteropus vampyrus | IAA | FJ561403 | FJ588905 | KP214033 | | | | |
| Pteropus vampyrus | IAA | JN398212 | JN398189 | JN398189 | JN398295 | JN398310 | JN398278 | JN398228 |
| Pteropus voeltzkowi | Africa | FJ561404 | FJ588906 | | | | | |
| Pteropus voeltzkowi | Africa | FJ561405 | FJ588907 | | | | | |
| Rousettus aegyptiacus | Africa | DQ445713 | AB205183 | AB205183 | EU617979 | | DQ445688 | KT875815 |
| Rousettus aegyptiacus | Africa | DQ445714 | AB205183 | AB205183 | KT875845 | | DQ445694 | |
| Rousettus amplexicaudatus | IAA | AB046329 | U93070 | AF203742 | AF447512 | AF447529 | AY057836 | AF447500 |
| Rousettus leschenaultii | India | JN398218 | JN398190 | JN398190 | JN398300 | JN398313 | JN398283 | JN398219 |
| Rousettus madagascariensis | Madagascar | JN398214 | JN398191 | JN398191 | JN398296 | JN398311 | JN398279 | JN398224 |
| Scotonycteris zenkeri | Africa | JN398216 | JN398192 | JN398192 | JN398297 | JN398312 | JN398280 | JN398265 |
| Sphaerias blanfordi | Asia | GQ410228 | GQ410322 | GQ410345 | GQ410273 | GQ410250 | GQ410296 | |
| Sphaerias blanfordi | Asia | GQ410229 | GQ410323 | GQ410346 | GQ410274 | GQ410251 | GQ410297 | JN398230 |
| Stenonycteris lanosus | Africa | JN398215 | JN398193 | JN398193 | JN398298 | JN400925 | JN398281 | JN398244 |
| Styloctenium mindorensis | IAA | JN398217 | JN398194 | JN398194 | JN398299 | | JN398282 | JN398227 |
| Syconycteris australis | Australia | FJ218479 | JN398195 | JN398195 | FJ218466 | FJ218462 | FJ218470 | JN398247 |
| Syconycteris australis | Australia | GQ410232 | U93060 | AF293650 | GQ410278 | GQ410255 | GQ410301 | |
| Thoopterus nigrescens | IAA | KC747699 | U93067 | AF293646 | | | | |
| Megaderma lyra | IAA | DQ888678 | AF069538 | AF069538 | AF203757 | AF203767 | U31616 | AF203749 |
| Megaderma lyra | India | DQ888678 | AF069538 | AF069538 | AF203757 | AF203767 | U31616 | AF203749 |
| Rhinopoma hardwickii | India | AY629005 | AF263232 | AF263232 | AF447518 | AF447535 | AF447551 | AF447504 |
| Rhinopoma hardwickii | Africa | AY629006 | AF263232 | AF263232 | AF447518 | AF447535 | AF447551 | AF447504 |
| Hipposideros commersoni | Madagascar | KR606333 | AY395856 | AY395856 | AF203760 | AF203770 | AF203778 | AF203752 |
| Artibeus jamaicensis | America | AF061340 | AF061340 | AF061340 | AY834655 | AY834663 | AY834737 | AY834647 |
| Rhinolophus creaghi | IAA | EF108163 | | | AF447511 | AF447528 | AF447546 | AF447499 |

Since we did not have a solid calibration point for this family without including a considerable number of microchiropteran terminals, we used the dates from Agnarsson, Zambrana-Torrelio, Flores-Saldana, and May-Collado (2011): a root prior with lognormal distribution (M = 4.075 and S = 0.05) and a Pteropodidae-Rhinolophoidea prior with lognormal distribution (M = 3.989 and S = 0.09).

#### 12. Boidae

Table S10: Genbank numbers for the sequences used in the Boidae analyses with the distribution attributed to each terminal.

| Species | Locality | CytB | CMOS | NT3 | BDNF | RAG1 | ODC |
|---|---|---|---|---|---|---|---|
| *Acrantophis dumerili* | Madagascar MI | U69735 | AY099963 | AY988049 | AY988032 | AY988066 | DQ465533 |
| *Acrochordus granulatus* | IAA | AF217841 | AF471124 | EU390905 | EU402621 | EU402831 | |
| *Acrochordus javanicus* | IAA Australia | KX694897 | HM234058 | AY988053 | AY988036 | AY988070 | DQ465538 |
| *Anilius scytale* | America | U69738 | AY099965 | AY988055 | AY988038 | AY988072 | DQ465540 |
| *Aspidites melanocephala* | Australia | U69741 | DQ465557 | DQ465558 | DQ465559 | DQ465560 | DQ465552 |
| *Boa constrictor* | America | U69740 | AF471115 | AY988047 | AY988030 | AY988064 | DQ465531 |
| *Calabaria reinhardtii* | Africa | AY099985 | AY099978 | AY988058 | AY988041 | AY988075 | DQ465542 |
| *Candoia carinata* | IAA | AY099984 | AY099961 | AY988048 | AY988031 | AY988065 | DQ465532 |
| *Candoia aspera* | IAA Australia | U69751 | | | | | |
| *Charina bottae* | America | AY099986 | AY099971 | AY988059 | AY988042 | AY988076 | DQ465543 |
| *Corallus caninus* | America | U69763 | AY987964 | AY988044 | AY988027 | AY988061 | DQ465528 |
| *Cylindrophis ruffus* | IAA | AF471032 | AF471133 | AY988054 | AY988037 | AY988071 | DQ465539 |
| *Epicrates cenchria* | America | U69777 | AY099966 | AY988045 | AY988028 | AY988062 | DQ465529 |
| *Epicrates striatus* | America | U69799 | DQ465553 | DQ465554 | DQ465555 | DQ465556 | DQ465551 |
| *Gongylophis colubrinus* | Africa | U69812 | DQ465568 | DQ465569 | DQ465570 | DQ465571 | DQ465550 |
| *Gongylophis conicus* | India | GQ225658 | DQ469787 | AY988057 | AY988040 | AY988074 | DQ465547 |
| *Eryx elegans* | Asia | U69818 | | | | | |
| *Eryx jayakari* | Arabic Peninsula | | DQ465565 | | DQ465566 | DQ465567 | DQ465548 |
| *Eryx johnii* | India | U69823 | AY099975 | DQ465575 | DQ465576 | DQ465577 | DQ465546 |
| *Eunectes notaeus* | America | U69810 | AY099964 | AY988046 | AY988029 | AY988063 | DQ465530 |
| *Exiliboa placata* | America | AY099989 | AY099973 | AY988051 | AY988034 | AY988068 | DQ465535 |
| *Lichanura trivirgata* | America | U69844 | AY099974 | DQ465578 | DQ465579 | DQ465580 | DQ465536 |
| *Loxocemus bicolor* | America | AY099993 | AY099969 | DQ465572 | DQ465573 | AY444061 | DQ465545 |
| *Morelia spilota* | Australia | U69851 | AF544723 | AY988052 | AY988035 | AY988069 | DQ465537 |
| *Ramphotyphlops braminus* | America | AY099990 | AY099980 | AY988060 | AY988043 | AY988077 | DQ465544 |
| *Sanzinia madagascariensis* | Madagascar | U69866 | AY099982 | AY988050 | AY988033 | AY988067 | DQ465534 |
| *Tropidophis haetianus* | America | U69868 | AY099962 | AY988056 | AY988039 | AY988073 | DQ465541 |
| *Xenopeltis unicolor* | IAA | AY121369 | DQ465561 | DQ465562 | DQ465563 | DQ465564 | DQ465549 |

Some of the calibration points used by the original article have been updated by recent discoveries. We set the following calibration points with lognormal distribution: *Coniophis* (M = 1.0 and S = 1.59; offset = 83.0 m.y.a. (Gómez, Báez, & Rougier, 2008; Marsh, 1892)); *Charina prebottae* (M = 1.0 and S = 0.8; offset = 13.6 m.y.a. (Brattstrom, 1958)); *Cheilophis* (M = 1.0 and S = 0.715; offset = 55.0 m.y.a. (Miller, 1955; Rage, 1984)); *Dunnophis* (M = 0.8 and S = 0.585; offset = 59.0 m.y.a. (Estes, 1976; Szyndlar, Smith, & Rage, 2008)); *Helagras* (M = 1.2 and S = 0.5; offset = 63.3 m.y.a. (Sullivan & Lucas, 1986)); *Dinilysia* (M = 1.25 and S = 1.42; offset = 89.0 m.y.a. (Caldwell & Albino, 2003)); *Pseudoepicrates* (M = 1.0 and S = 0.94; offset = 15.97 m.y.a. (Onary & Hsiou, 2018)); and Nigerophiidae (M = 1.5 and S = 1.5; offset = 66.0 m.y.a. (Laduke, Krause, Scanlon, & Kley, 2010)).

#### 13. Chamaeleoninae

The sequences used in the Chamaeleoninae analyses were downloaded directly from the original article's dryad repository (https://datadryad.org/resource/doi:10.5061/dryad.11350). All calibration points were identical to those of the original article.

#### 14. Gekkonidae

Table S11: Genbank numbers for the sequences used in the Gekkonidae analyses with the distribution attributed to each terminal.

| Species | Location | 12S | CMOS | CytB | ND2 | PDC | RAG1 | RAG2 |
|---|---|---|---|---|---|---|---|---|
| *Afroedura karroica* | Africa | | JQ945523 | | JX041302 | JQ945345 | JQ945277 | JQ945415 |
| *Afroedura multiporis* | Africa | | | | EU054232 | EU054184 | EU054208 | |
| *Afroedura pondolia* | Africa | | | | EU054231 | | EU054207 | |
| *Afrogecko porphyreus* | Africa | DQ852696 | AY172919 | EF490750 | EF490776 | EF490697 | EF490723 | JQ945418 |
| *Afrogekko plumicaudus* | Africa | | JQ945525 | | JX041304 | JQ945347 | JQ945279 | JQ945417 |
| *Afrogecko swartbergensis* | Africa | | JQ945527 | | JX041305 | JQ945348 | JQ945280 | JQ945419 |
| *Agamura persica* | Asia | DQ852726 | DQ852728 | EU589176 | JX041306 | JQ945349 | JQ945281 | JQ945420 |
| *Ailuronyx seychellensis* | Seychelles | AY221267 | AY221331 | FJ830052 | | | KY037922 | |
| *Ailuronyx tachyscopaeus* | Seychelles | | JQ945529 | | JX041307 | JQ945350 | JQ945282 | JQ945421 |
| *Ailuronyx trachygaster* | Seychelles | | JQ945530 | | JX041308 | JQ945351 | JQ945283 | JQ945422 |
| *Alsophylax pipiens* | Asia | | JQ945531 | | JX041309 | JQ945352 | JQ945284 | JQ945423 |
| *Blaesodactylus antongilensis* | Madagascar | | JQ945534 | | EU054254 | EU054206 | EU054230 | JQ945426 |
| *Blaesodactylus boivini* | Madagascar | EU596607 | | | EU054252 | EU054204 | EU054228 | |
| *Blaesodactylus sakalava* | Madagascar | | | | EU054251 | EU054203 | EU054227 | |
| *Bunopus crassicauda* | Arabic Peninsula | EU589154 | | EU589177 | | | | |
| *Bunopus tuberculatus* | Arabic Peninsula | EU589160 | AF148706 | EU589181 | HQ443541 | JQ945355 | JQ945287 | JQ945427 |
| *Calodactylodes aureus* | India | DQ852697 | AY172921 | | | | | |
| *Calodactylodes illingworthorum* | India | | JQ945536 | | JX041318 | JQ945356 | JQ945288 | JQ945428 |
| *Chondrodactylus angulifer* | Africa | DQ275403 | JQ945537 | AY123394 | JX041320 | JQ945357 | JQ073242 | JQ945429 |
| *Chondrodactylus bibronii* | Africa | | EU293690 | | | EU293712 | EU293645 | EU293735 |
| *Chondrodactylus fitzsimonsi* | Africa | DQ275404 | | AF449125 | JN393945 | | DQ275448 | |
| *Chondrodactylus turneri* | Africa | DQ852708 | AY172938 | AF449124 | KY224249 | KM073612 | KM073525 | |
| *Christinus alexanderi* | Australia | | | | KF666813 | | KF666897 | KF666856 |
| *Christinus guentheri* | Australia | | | | KF666800 | | KF666887 | KF666846 |
| *Christinus marmoratus* | Australia | DQ852698 | FJ855461 | | JX041322 | JQ945358 | FJ855440 | JQ945430 |
| *Cnemaspis africana* | Africa | | JQ945539 | | JX041323 | JQ945359 | JQ945291 | JQ945431 |
| *Cnemaspis dickersonae* | Africa | | JQ945540 | | JX041324 | JQ945360 | JQ945292 | JQ945432 |
| *Cnemaspis kandiana* | India | | JQ945541 | | JX041325 | JQ945361 | JQ945293 | JQ945433 |
| *Cnemaspis kendallii* | IAA | | AY172923 | | JX041326 | JQ945362 | JQ945294 | JQ945434 |
| *Cnemaspis limi* | IAA | | EF534935 | | JX041327 | EF534851 | EF534809 | EF534977 |
| *Cnemaspis podihuna* | India | | JQ945543 | | JX041328 | JQ945363 | JQ945295 | JQ945435 |
| *Cnemaspis tropidogaster* | India | DQ852716 | DQ852729 | | | | | |

| | | | | | | | | |
|---|---|---|---|---|---|---|---|---|
| *Cnemaspis uzungwae* | Africa | | JQ945544 | | JX041329 | JQ945364 | JQ945296 | JQ945436 |
| *Colopus kochii* | Africa | | JQ945545 | AY123398 | JX041336 | JQ945365 | DQ275418 | JQ945437 |
| *Colopus wahlbergii* | Africa | DQ275375 | JQ945546 | AY123395 | JX041337 | JQ945366 | DQ275419 | JQ945438 |
| *Crossobamon orientalis* | Asia | DQ852715 | DQ852730 | HM921200 | HQ443529 | JQ945368 | JQ945299 | JQ945440 |
| *Cryptactites peringueyi* | Africa | DQ852718 | DQ852731 | | JX041339 | JQ945369 | JQ945300 | JQ945441 |
| *Cyrtodactylus angularis* | Australia IAA India Asia | | JQ945549 | | HQ401212 | JQ945370 | JQ945301 | JQ945442 |
| *Cyrtodactylus ayeyarwadyensis* | Australia IAA India Asia | | JQ945550 | EU268380 | EU268348 | EU268317 | | JQ945443 |
| *Cyrtodactylus consobrinus* | Australia IAA India Asia | | | EU268381 | EU268349 | EU268318 | | |
| *Cyrtodactylus irregularis* | Australia IAA India Asia | | JQ945551 | | JX041341 | JQ945371 | JQ945302 | JQ945444 |
| *Cyrtodactylus jarujini* | Australia IAA India Asia | | JQ945552 | | HQ401213 | JQ945372 | JQ945303 | JQ945445 |
| *Cyrtodactylus loriae* | Australia IAA India Asia | | | EU268382 | EU268350 | EU268319 | | |
| *Cyrtodactylus novaeguineae* | Australia IAA India Asia | | HQ426531 | | HQ401210 | HQ426185 | HQ426274 | HQ426447 |
| *Cyrtodactylus philippinicus* | Australia IAA India Asia | | JQ945553 | | JX041344 | JQ945373 | JQ945304 | JQ945446 |
| *Cyrtopodion agamuroides* | Asia | EU589161 | | EU589185 | | | | |
| *Cyrtopodion caspium* | Asia | EU589163 | JQ945620 | EU589187 | JX041448 | JQ945409 | JQ945340 | JQ945514 |
| *Cyrtopodion longipes* | Asia | EU589170 | JQ945621 | EU589193 | JX041449 | JQ945410 | JQ945341 | JQ945515 |
| *Cyrtopodion russowii* | Asia | | JQ945588 | | JX041384 | JQ945383 | JQ945315 | JQ945481 |
| *Cyrtopodion scabrum* | Asia | EU589172 | HQ426532 | EU589195 | JX041345 | HQ426186 | HQ426275 | HQ426448 |
| *Cyrtopodion spinicaudum* | Asia | | JQ945589 | | JX041385 | JQ945384 | JQ945316 | JQ945482 |
| *Delma butleri* | Australia | | AY134548 | | AY134584 | HQ426187 | KP851280 | HQ426449 |
| *Dixonius siamensis* | IAA | | JQ945557 | | EU054299 | EU054267 | EU054283 | JQ945450 |
| *Dixonius vietnamensis* | IAA | | JQ945558 | | EU054298 | EU054266 | EU054282 | JQ945451 |
| *Ebenavia inunguis* | Africa Comoros Madagascar MI | EU596608 | FJ830144 | FJ830053 | EF536191 | HQ426191 | EF536143 | HQ426453 |
| *Elasmodactylus tetensis* | Africa | DQ275407 | JQ945559 | AY026926 | JX041349 | JQ945376 | DQ275451 | JQ945452 |
| *Elasmodactylus tuberculosus* | Africa | DQ275408 | | AF449134 | | | DQ275452 | |
| *Geckolepis maculata* | Comoros Madagascar | EU596610 | JQ945562 | | EU054235 | EU054187 | EU054211 | JQ945455 |
| *Geckolepis typica* | Madagascar | | | | EU054233 | EU054185 | EU054209 | |
| *Geckonia chazaliae* | Africa | AF363574 | AF363556 | AF364326 | JX041443 | EU293705 | EU293638 | JQ301343 |
| *Gehyra australis* | Australia | | JQ945563 | | GQ257759 | JN019113 | | JQ945456 |
| *Gehyra barea* | IAA | | | | JN393915 | JN393993 | JN393960 | |
| *Gehyra brevipalmata* | IAA | | | | JN393910 | JN393987 | JN393955 | |
| *Gehyra dubia* | Australia | | JQ945565 | | GQ257788 | JN393989 | | JQ945458 |
| *Gehyra marginata* | IAA | | | | JN393931 | JN394009 | JN393975 | |
| *Gehyra mutilata* | IAA | DQ852699 | FJ830146 | FJ830054 | GQ257784 | JN019114 | FJ830237 | FJ830328 |
| *Gehyra nana* | Australia | | JQ945567 | | GQ257760 | JN393998 | | JQ945460 |
| *Gehyra oceanica* | Pacific | | | | GQ257785 | JN394001 | | |
| *Gehyra variegata* | Australia | AF090185 | AF090851 | | AY369026 | JN393994 | | JQ945461 |
| *Gehyra xenopus* | Australia | | | | GQ257790 | JN394010 | | |
| *Gekko athymus* | IAA | | | | JN019075 | JN019107 | JN019139 | |
| *Gekko auriverrucosus* | Asia | | | EU417695 | JN019062 | JN019096 | JN019127 | |
| *Gekko badenii* | IAA | | JQ945569 | | JN019065 | JN019099 | JN019130 | JQ945462 |
| *Gekko chinensis* | Asia | | JQ945571 | EU417676 | JN019058 | JN019092 | JN019123 | JQ945464 |
| *Gekko gecko* | Asia Pacific | HM370130 | EU366455 | NC_007627 | AY282753 | JN019087 | AY662625 | EF534981 |

| | | | | | | | | |
|---|---|---|---|---|---|---|---|---|
| *Gekko grossmanni* | IAA | | | | JN019064 | JN019098 | JN019129 | |
| *Gekko hokouensis* | Asia | AF323511 | | EU417691 | JN019060 | JN019094 | JN019125 | |
| *Gekko japonicus* | Asia | AF318271 | | EU417688 | JN019059 | JN019093 | JN019124 | |
| *Gekko mindorensis* | IAA | | JQ945572 | | FJ487887 | JN019108 | JN019140 | JQ945465 |
| *Gekko monarchus* | IAA | | JQ945573 | | FJ487870 | JN019110 | JN019141 | JQ945466 |
| *Gekko petricolus* | IAA | | | | JN019067 | JN019101 | JN019132 | |
| *Gekko smithii* | Asia IAA | | | | FJ487868 | JN019091 | JN019121 | |
| *Gekko swinhonis* | Asia | AF323518 | | EU417702 | JN019061 | JN019095 | JN019126 | |
| *Gekko vittatus* | IAA | NC_008772 | JQ945575 | NC_008772 | AB178897 | JN019106 | JN019137 | JQ945468 |
| *Goggia lineata* | Africa | AY763261 | AY172930 | | JX041353 | JQ945378 | JQ945310 | JQ945469 |
| *Hemidactylus aaronbaueri* | India | HM595676 | | HM595641 | | HM622367 | | |
| *Hemidactylus angulatus* | Africa | EF202130 | HQ426540 | DQ120237 | EU268367 | EU268336 | HM559686 | |
| *Hemidactylus bouvieri* | Africa | DQ120424 | AY863042 | DQ120249 | | | | EF540746 |
| *Hemidactylus bowringii* | Asia | AF323515 | | EU268405 | EU268374 | EU268343 | | |
| *Hemidactylus brasilianus* | America | | HQ426523 | DQ120257 | EU268351 | EU268320 | | HQ426439 |
| *Hemidactylus depressus* | India | | | HM559594 | HM559626 | HM559659 | HM559692 | |
| *Hemidactylus dracaenacolus* | Arabic Peninsula | DQ120380 | | DQ120209 | | | | |
| *Hemidactylus fasciatus* | Africa | DQ852724 | JQ945577 | EU268403 | EU268371 | EU268340 | JQ945311 | JQ945470 |
| *Hemidactylus flaviviridis* | Africa Arabic Peninsula Asia | DQ120454 | HQ426541 | EU268388 | EU268356 | EU268325 | HM559694 | HQ426458 |
| *Hemidactylus forbesii* | Arabic Peninsula | DQ120339 | | DQ120168 | | | | |
| *Hemidactylus foudaii* | Africa | DQ120385 | | DQ120214 | | | | |
| *Hemidactylus frenatus* | Australia IAA India Asia | NC_012902 | EF534940 | EU116513 | EU268359 | EU268328 | EF534814 | EF534982 |
| *Hemidactylus garnotii* | Australia IAA | DQ120459 | | EU268395 | EU268364 | EU268333 | HM559697 | |
| *Hemidactylus giganteus* | India | HM595692 | | HM595658 | | HM559665 | | |
| *Hemidactylus gracilis* | India | HM595696 | | HM595660 | EU268379 | HM622374 | | |
| *Hemidactylus granti* | Arabic Peninsula | DQ120381 | | DQ120210 | | | | |
| *Hemidactylus greefii* | Africa | DQ120413 | AY863044 | EU268401 | EU268369 | EU268338 | | HQ426459 |
| *Hemidactylus karenorum* | IAA | DQ120464 | | EU268394 | EU268362 | EU268331 | | |
| *Hemidactylus leschenaultii* | India | HM595697 | | HM595662 | | HM559669 | | |
| *Hemidactylus longicephalus* | Africa | DQ120416 | HQ426544 | DQ120246 | HM559637 | HM559670 | HM559703 | HQ426460 |
| *Hemidactylus mabouia* | Africa | HM180323 | HQ426545 | EU268393 | EU268361 | EU268330 | HM559704 | HQ426462 |
| *Hemidactylus macropholis* | Africa | DQ120343 | HQ426547 | DQ120208 | JX041369 | HQ426203 | HQ426292 | HQ426463 |
| *Hemidactylus maculatus* | India | HM595699 | | HM595664 | | HM559674 | | |
| *Hemidactylus modestus* | Africa | DQ120386 | | DQ120215 | | | | |
| *Hemidactylus oxyrhinus* | Arabic Peninsula | DQ120344 | | DQ120173 | | | | |
| *Hemidactylus palaichthus* | America | DQ120434 | HQ426548 | EU268400 | EU268368 | EU268337 | | HQ426464 |
| *Hemidactylus persicus* | Asia | HM595701 | | EU268409 | EU268377 | EU268346 | | |
| *Hemidactylus platycephalus* | Africa Madagascar | DQ120437 | AY863045 | DQ120270 | | | JQ073246 | |
| *Hemidactylus platyurus* | Asia IAA India | | HQ426530 | EU268384 | EU268352 | EU268321 | HM559685 | HQ426446 |
| *Hemidactylus prashadi* | India | HM595702 | | HM595668 | | HM559676 | | |
| *Hemidactylus pumilio* | Arabic Peninsula | DQ120382 | | DQ120211 | | | | |
| *Hemidactylus reticulatus* | India | HM595705 | | EU268410 | | EU268347 | | |
| *Hemidactylus robustus* | Africa Arabic Peninsula | DQ120345 | HQ426549 | EU268408 | EU268376 | EU268345 | EU054271 | HQ426465 |

| | | | | | | | | |
|---|---|---|---|---|---|---|---|---|
| *Hemidactylus sataraensis* | India | HM595708 | | HM595672 | | | | |
| *Hemidactylus triedrus* | India | HM595709 | HQ426550 | HM595675 | | HM559683 | | HQ426466 |
| *Hemiphyllodactylus aurantiacus* | India | | | FJ971012 | JN393933 | JN394011 | JN393977 | |
| *Hemiphyllodactylus typus* | Asia IAA India Pacific | | | FJ971011 | GQ257744 | | | |
| *Hemiphyllodactylus yunnanensis* | Asia | | JQ945579 | FJ971005 | JN393949 | JN394013 | JN393979 | JQ945472 |
| *Heteronotia binoei* | Australia | NC_010292 | JQ945580 | NC_010292 | NC_010292 | EU054270 | EU054285 | JQ945473 |
| *Heteronotia planiceps* | Australia | | JQ945581 | | EU054300 | EU054268 | EU054284 | JQ945474 |
| *Homopholis fasciata* | Africa | DQ852723 | DQ852735 | | EU054250 | EU054202 | EU054226 | JQ945475 |
| *Homopholis mulleri* | Africa | | | | EU054241 | EU054193 | EU054217 | |
| *Homopholis walbergii* | Africa | | | | EU054248 | EU054200 | EU054224 | |
| *Lepidodactylus moestus* | Pacific | | | | JN019079 | JN019111 | JN019143 | |
| *Lepidodactylus novaeguineae* | IAA | | JQ945583 | | JX041378 | JQ945380 | JQ945312 | JQ945476 |
| *Lepidodactylus orientalis* | IAA | | | | JN019080 | JN019112 | JN019144 | |
| *Luperosaurus cumingii* | IAA | | JQ945585 | | JQ437902 | JQ945381 | JQ945313 | JQ945478 |
| *Lygodactylus arnoulti* | Madagascar | | | GU593523 | | | GU593585 | GU593395 |
| *Lygodactylus bradfieldi* | Africa | DQ852705 | AY172935 | | EU423279 | HQ426212 | HQ426301 | GU593360 |
| *Lygodactylus gravis* | Africa | | | GU593500 | | | GU593556 | GU593366 |
| *Lygodactylus gutturalis* | Africa | | | GU593522 | | | GU593583 | GU593393 |
| *Lygodactylus heterurus* | Madagascar | | | GU593546 | | | GU593611 | GU593421 |
| *Lygodactylus klugei* | America | | HQ426555 | | | HQ426209 | HQ426298 | HQ426471 |
| *Lygodactylus lawrencei* | Africa | | | GU593496 | | | GU593552 | GU593362 |
| *Lygodactylus luteopicturatus* | Africa | | FJ830142 | FJ830051 | | | FJ830234 | |
| *Lygodactylus madagascariensis* | Madagascar | EU596612 | | EU596695 | | | | |
| *Lygodactylus miops* | Madagascar | | HQ426556 | | | HQ426210 | HQ426299 | HQ426472 |
| *Lygodactylus mirabilis* | Madagascar | | HQ426557 | GQ910848 | JX041382 | HQ426211 | HQ426300 | GU593413 |
| *Lygodactylus montanus* | Madagascar | | | GU593536 | | | GU593601 | GU593411 |
| *Lygodactylus picturatus* | Africa | | | FJ830051 | | | JQ073235 | FJ830325 |
| *Lygodactylus tolampyae* | Madagascar | | HQ426559 | GU593526 | JX041383 | HQ426213 | HQ426302 | GU593399 |
| *Lygodactylus verticillatus* | Madagascar | | | GU593529 | | | GU593595 | GU593404 |
| *Matoatoa brevipes* | Madagascar | EU596614 | JQ945587 | EF490751 | EF490777 | EF490698 | EF490724 | JQ945480 |
| *Nactus acutus* | IAA | | | | EU054289 | EU054257 | EU054273 | |
| *Nactus cheverti* | Australia | | | | HM997154 | HM997178 | HM997166 | |
| *Nactus eboracensis* | Australia | | | | HM997156 | HM997180 | HM997168 | |
| *Nactus galgajuga* | Australia | | | | HM997158 | HM997182 | HM997170 | |
| *Nactus multicarinatus* | Australia IAA | | | | HM997160 | HM997184 | HM997172 | |
| *Nactus pelagicus* | Pacific | DQ852706 | AY172936 | | HM997163 | HM997187 | EU054275 | JQ945484 |
| *Nactus vankampeni* | IAA | | JQ945591 | | EU054296 | EU054264 | EU054280 | JQ945485 |
| *Narudasia festiva* | Africa | DQ852707 | EF534934 | | JX041387 | EF534850 | EF534808 | EF534976 |
| *Pachydactylus affinis* | Africa | DQ275389 | | AY123414 | | | DQ275433 | |
| *Pachydactylus austeni* | Africa | DQ275371 | JQ945596 | AF449126 | JX041390 | JQ945389 | DQ275415 | |
| *Pachydactylus barnardi* | Africa | DQ275378 | | AY123410 | | | DQ275422 | |
| *Pachydactylus bicolor* | Africa | DQ275394 | | AY123400 | | | DQ275438 | |
| *Pachydactylus capensis* | Africa | DQ275387 | | AF449133 | HQ165962 | HQ165977 | DQ275431 | |

| | | | | | | | | |
|---|---|---|---|---|---|---|---|---|
| *Pachydactylus caraculicus* | Africa | DQ275393 | | AY123399 | | | DQ275437 | |
| *Pachydactylus fasciatus* | Africa | DQ275384 | | AF449128 | HQ165949 | HQ165964 | DQ275428 | |
| *Pachydactylus formosus* | Africa | DQ275377 | | AY123411 | | | DQ275421 | |
| *Pachydactylus gaiasensis* | Africa | DQ275399 | JQ945597 | AY123407 | JX041391 | JQ945390 | DQ275443 | JQ945490 |
| *Pachydactylus geitje* | Africa | DQ275380 | | AF449132 | | | DQ275424 | |
| *Pachydactylus haackei* | Africa | DQ275402 | | AF449121 | | | DQ275446 | |
| *Pachydactylus kladaroderma* | Africa | DQ275401 | JQ945598 | AF449122 | JX041392 | JQ945391 | DQ275445 | JQ945491 |
| *Pachydactylus labialis* | Africa | DQ275379 | | AF449130 | | | DQ275423 | |
| *Pachydactylus maculatus* | Africa | DQ275382 | | AF449127 | | | DQ275426 | |
| *Pachydactylus mariquensis* | Africa | DQ275370 | | DQ278601 | | | DQ275414 | |
| *Pachydactylus mclachlani* | Africa | | | | HQ165951 | HQ165966 | HQ165981 | |
| *Pachydactylus monicae* | Africa | | | DQ349180 | HQ165953 | HQ165968 | HQ165983 | |
| *Pachydactylus namaquensis* | Africa | DQ275400 | | AF449123 | | | DQ275444 | |
| *Pachydactylus oculatus* | Africa | DQ275381 | | AY123402 | | | DQ275425 | |
| *Pachydactylus oreophilus* | Africa | DQ275398 | | AY123406 | | | DQ275442 | |
| *Pachydactylus oshaughnessyi* | Africa | DQ275391 | | AY123413 | | | DQ275435 | |
| *Pachydactylus parascutatus* | Africa | DQ275397 | | AY123405 | | | DQ275441 | |
| *Pachydactylus punctatus* | Africa | DQ275411 | EU293691 | AF449135 | JX041393 | EU293713 | EU293646 | EU293736 |
| *Pachydactylus purcelli* | Africa | | | DQ349161 | HQ165955 | HQ165970 | HQ165985 | |
| *Pachydactylus rangei* | Africa | | JQ945599 | AY123397 | JX041394 | JQ945392 | JQ945324 | JQ945492 |
| *Pachydactylus rugosus* | Africa | DQ275376 | JQ945600 | AF449129 | JX041395 | JQ945393 | DQ275420 | JQ945493 |
| *Pachydactylus scherzi* | Africa | DQ275392 | | AY123401 | | | DQ275436 | |
| *Pachydactylus scutatus* | Africa | DQ275395 | | AY123403 | | | DQ275439 | |
| *Pachydactylus serval* | Africa | DQ275385 | | DQ349168 | HQ165957 | HQ165972 | DQ275429 | |
| *Pachydactylus tigrinus* | Africa | DQ275390 | | AY123412 | | | DQ275434 | |
| *Pachydactylus tsodiloensis* | Africa | DQ275383 | | AY123408 | | | DQ275427 | |
| *Pachydactylus vansoni* | Africa | DQ275388 | | AY123415 | | | DQ275432 | |
| *Pachydactylus vanzyli* | Africa | | JQ945601 | AY123396 | JX041396 | JQ945394 | JQ945326 | JQ945494 |
| *Pachydactylus weberi* | Africa | DQ275386 | JQ945602 | AF449131 | HQ165961 | HQ165976 | DQ275430 | JQ945495 |
| *Paragehyra gabriellae* | Madagascar | | JQ945603 | | JX041399 | JQ945396 | JQ945328 | JQ945496 |
| *Paroedura androyensis* | Madagascar | GU128974 | HQ256721 | EF490748 | EF490774 | EF490695 | EF490721 | |
| *Paroedura bastardi* | Madagascar | GU128980 | HQ256725 | | EF536211 | EF536187 | EF536163 | |
| *Paroedura gracilis* | Madagascar | GU128969 | HQ256726 | | EF536209 | EF536185 | EF536161 | |
| *Paroedura homalorhina* | Madagascar | | | | EF536214 | EF536190 | EF536166 | |
| *Paroedura karstophila* | Madagascar | | | EF490749 | EF490775 | EF490696 | EF490722 | |
| *Paroedura lohatsara* | Madagascar | GU128971 | HQ256731 | | EF536203 | EF536179 | EF536155 | |
| *Paroedura masobe* | Madagascar | GU128977 | HQ426560 | | EF536193 | EF536169 | EF536145 | HQ426478 |
| *Paroedura oviceps* | Madagascar | EU596615 | HQ256733 | | EF536208 | EF536184 | EF536160 | |
| *Paroedura picta* | Madagascar | GU128965 | EU293692 | | EF536198 | EF536174 | EF536150 | EU293737 |
| *Paroedura sanctijohannis* | Comoros | | | | EF536205 | EF536181 | EF536157 | |
| *Paroedura stumpffi* | Madagascar | GU128964 | HQ256740 | | EF536202 | EF536178 | EF536154 | |
| *Paroedura tanjaka* | Madagascar | | | | EF536201 | EF536177 | EF536153 | |
| *Paroedura vazimba* | Madagascar | GU128981 | HQ256742 | | EF536195 | EF536171 | EF536147 | |

| | | | | | | | | |
|---|---|---|---|---|---|---|---|---|
| *Perochirus ateles* | Pacific | | JQ945604 | | JN393946 | JN394015 | JN393984 | JQ945497 |
| *Phelsuma abbotti* | IAA | EU596618 | AY221346 | FJ829977 | EU423290 | | FJ830148 | FJ830239 |
| *Phelsuma andamanense* | IAA | AY221277 | FJ830063 | FJ829982 | | | FJ830155 | FJ830246 |
| *Phelsuma antanosy* | Madagascar | | FJ830064 | FJ829983 | | | FJ830156 | FJ830247 |
| *Phelsuma astriata* | Seychelles | AY221273 | AY221337 | FJ829985 | EU423286 | | FJ830158 | FJ830249 |
| *Phelsuma barbouri* | Madagascar | | FJ830069 | FJ829988 | EU423289 | | FJ830161 | FJ830252 |
| *Phelsuma berghofi* | Madagascar | | FJ830070 | FJ829989 | | | FJ830162 | FJ830253 |
| *Phelsuma borbonica* | MI | AY221286 | AY221350 | FJ829990 | JX041400 | HQ426216 | FJ830163 | HQ426479 |
| *Phelsuma breviceps* | Madagascar | | FJ830072 | FJ829991 | | | FJ830164 | FJ830255 |
| *Phelsuma cepediana* | MI | AY221294 | FJ830073 | FJ829992 | EU423294 | | FJ830165 | FJ830256 |
| *Phelsuma comorensis* | Comoros | DQ911636 | | | | | JQ073229 | FJ830257 |
| *Phelsuma dubia* | Africa | EF424452 | FJ830080 | FJ829996 | EU423285 | | FJ830172 | FJ830263 |
| *Phelsuma edwardnewtoni* | MI | AY221292 | AY221353 | AY221398 | | | | |
| *Phelsuma flavigularis* | Madagascar | EF424466 | FJ830082 | FJ829997 | | | FJ830174 | FJ830265 |
| *Phelsuma gigas* | MI | AY221293 | AY221354 | AY221399 | | | | |
| *Phelsuma guentheri* | MI | AY221310 | FJ830084 | FJ830000 | | | FJ830176 | FJ830267 |
| *Phelsuma guimbeaui* | MI | | AY221372 | FJ830041 | EU423291 | HQ426217 | HQ426306 FJ830221 | HQ426480 |
| *Phelsuma guttata* | Madagascar | | FJ830085 | FJ830001 | EU423284 | | FJ830177 | FJ830268 |
| *Phelsuma hielscheri* | Madagascar | | FJ830086 | FJ830002 | | | FJ830178 | FJ830269 |
| *Phelsuma inexpectata* | MI | | FJ830087 | FJ830003 | JN393939 | JN394016 | FJ830179 | FJ830270 |
| *Phelsuma kely* | Madagascar | | FJ830088 | FJ830004 | | | FJ830180 | FJ830271 |
| *Phelsuma klemmeri* | Madagascar | | FJ830090 | FJ830006 | | | FJ830182 | FJ830273 |
| *Phelsuma laticauda* | Comoros Madagascar | DQ911673 | AY221343 | FJ830012 | EU423281 | JQ945398 | FJ830189 | FJ830280 |
| *Phelsuma lineata* | Madagascar | AY221279 | AY221342 | FJ830021 | EU423283 | | JN654861 | FJ830285 |
| *Phelsuma madagascariensis* | Madagascar | DQ852709 | EF534937 | EF424450 | EU423288 | AB081507 | EF534811 FJ830175 | EF534979 |
| *Phelsuma malamakibo* | Madagascar | | FJ830108 | FJ830023 | | | FJ830200 | FJ830291 |
| *Phelsuma modesta* | Madagascar | EF424465 | FJ830110 | EF424451 | EU423287 | HQ426218 | FJ830202 | HQ426481 |
| *Phelsuma mutabilis* | Madagascar | AY221272 | FJ830114 | FJ830029 | | | FJ830206 | FJ830297 |
| *Phelsuma nigristriata* | Comoros | | FJ830117 | FJ830032 | | | FJ830209 | FJ830300 |
| *Phelsuma ornata* | MI | AY221323 | FJ830118 | FJ830033 | EU423282 | | FJ830210 | FJ830301 |
| *Phelsuma parkeri* | Africa | DQ911693 | FJ830119 | DQ911751 | | | FJ830211 | FJ830302 |
| *Phelsuma pronki* | Madagascar | | FJ830121 | FJ830034 | | | FJ830213 | FJ830304 |
| *Phelsuma pusilla* | Madagascar | | FJ830122 | FJ830035 | | | FJ830214 | FJ830305 |
| *Phelsuma quadriocellata* | Madagascar | AY221282 | FJ830123 | FJ830036 | | | FJ830215 | FJ830306 |
| *Phelsuma ravenala* | Madagascar | | FJ830078 | FJ829994 | | | FJ830170 | FJ830261 |
| *Phelsuma robertmertensi* | Comoros | DQ911643 | FJ830128 | DQ911701 | | | FJ830220 | FJ830311 |
| *Phelsuma seippi* | Madagascar | | FJ830130 | FJ830042 | | | FJ830222 | FJ830313 |
| *Phelsuma serraticauda* | Madagascar | AY221278 | FJ830131 | FJ830043 | EU423296 | | FJ830223 | FJ830314 |
| *Phelsuma standingi* | Madagascar | AY221281 | FJ830133 | FJ830045 | | | FJ830225 GU593609 | FJ830316 |
| *Phelsuma sundbergi* | Seychelles | | | FJ830048 | EU423295 | | FJ830228 | FJ830319 |
| *Phelsuma vanheygeni* | Madagascar | | FJ830137 | FJ830049 | | | FJ830229 | FJ830320 |
| *Pseudogekko smaragdinus* | IAA | | JQ945608 | | JQ437897 | JQ437940 | JQ945332 | JQ945501 |
| *Ptenopus carpi* | Africa | DQ852711 | AY172942 | | JX041422 | JQ945402 | JQ945333 | JQ945502 |

| | | | | | | | | |
|---|---|---|---|---|---|---|---|---|
| *Ptychozoon kuhli* | IAA | | JQ945610 | | JQ437918 | JQ437960 | JQ945334 | JQ945503 |
| *Ptychozoon lionotum* | IAA | | JQ945611 | | JQ437914 | JQ437956 | JQ945335 | JQ945504 |
| *Pygopus nigriceps* | Australia | DQ852693 | AY172943 | | | | | |
| *Rhoptropus afer* | Africa | DQ275409 | JQ945613 | AY026923 | JX041430 | JQ945405 | DQ275453 | JQ945506 |
| *Rhoptropus boultoni* | Africa | DQ852712 | EF534936 | AY026922 | JX041431 | EF534852 | EF534810 | EF534978 |
| *Rhoptropus bradfieldi* | Africa | EU596606 | JQ945614 | AY026924 | JX041432 | JQ945406 | JQ945337 | JQ945507 |
| *Stenodactylus petrii* | Africa Arabic Peninsula | DQ852722 | DQ852738 | | HQ443558 | | JX440679 | |
| *Stenodactylus sthenodactylus* | Africa Arabic Peninsula | | JQ945617 | | HQ443546 | JQ945408 | JQ945339 | JQ945510 |
| *Microgecko helenae* | Asia | | JQ945590 | | JX041386 | JQ945385 | JQ945317 | JQ945483 |
| *Tropiocolotes tripolitanus* | Africa | | JQ945623 | | HQ443537 | JQ945412 | JQ945343 | JQ945517 |
| *Urocotyledon inexpectata* | Africa Seychelles | | JQ945624 | JN225804 | JX041461 | JN225672 | JQ945344 | JQ945518 |
| *Uroplatus alluaudi* | Madagascar | EU596620 | | EF490766 | EF490793 | EF490713 | EF490740 | |
| *Uroplatus ebenaui* | Madagascar | EU596624 | JN038097 | EF490763 | EF490788 | EF490710 | EF490736 | |
| *Uroplatus fimbriatus* | Madagascar | EU596641 | | | EF490792 | EF490712 | EF490739 | |
| *Uroplatus giganteus* | Madagascar | | JQ945625 | EF490765 | EF490791 | EF490711 | EF490738 | JQ945519 |
| *Uroplatus guentheri* | Madagascar | EU596689 | JQ945626 | EF490752 | EF490778 | EF490699 | EF490725 | JQ945520 |
| *Uroplatus henkeli* | Madagascar | EU596650 | HQ426591 | EF490770 | EF490797 | EF490717 | EF490744 | HQ426510 |
| *Uroplatus lineatus* | Madagascar | EU596656 | | EF490754 | EF490780 | EF490701 | EF490727 | |
| *Uroplatus phantasticus* | Madagascar | EU596664 | HQ426592 | EF490773 | EF490800 | EF490720 | EF490747 | HQ426511 |
| *Uroplatus pietschmanni* | Madagascar | EU596687 | | EF490755 | EF490781 | EF490702 | EF490728 | |
| *Uroplatus sikorae* | Madagascar | EU596681 | | EF490758 | EF490785 | EF490706 | EF490731 | |
| *Oedura marmorata* | Australia | | | | AY369015 | EF534819 | EF534779 | EF534945 |
| *Sphaerodactylus elegans* | America | KF017637 | EF534912 | KF017628 | JN393942 | EF534828 | EF534787 | EF534954 |
| *Sphaerodactylus roosevelti* | America | DQ852713 | AY172946 | | JN393943 | EF534825 | EF534785 | EF534951 |
| *Woodworthia maculata* | Australia | | JQ945628 | | GU459852 | GU459651 | GU459449 | JQ945522 |

The calibration points used in this run were taken from Heinicke, Daza, Greenbaum, Jackman, & Bauer (2014): one point for the Diplodactylidae (most recent common ancestor between *Oedura* and *Woodworthia*; exponential distribution; M = 17.0 and offset = 16.0 m.y.a.); one point for the Pygopodidae (most recent common ancestor between *Delma* and *Pygopus*; exponential distribution; M = 10.0 and offset = 20.0 m.y.a.); one point for the Sphaerodactylidae (most recent common ancestor between *Sphaerodactylus elegans* and *S. roosevelti*; exponential distribution; M = 3.0 and offset = 15.0 m.y.a.); and one point for the Gekkonidae (ingroup; lognormal distribution; M = 3.0 and S = 1.0; offset = 100.0 m.y.a.). In this last calibration point, we altered the offset from 97 to 100 m.y.a. based on the original description of *Cretaceogekko burmae* (Arnold & Poinar, 2008).

#### 15. Typhlopoidea

Table S12: Genbank numbers for the sequences used in the Typhlopoidea analyses with the distribution attributed to each terminal

| Species | Location | BDNF | BMP2 | NT3 | AMEL | RAG1 |
|---|---|---|---|---|---|---|
| *Acutotyphlops kunuaensis* | IAA | GU902419 | GU902499 | GU902590 | GU902339 | GU902669 |
| *Acutotyphlops sp* | IAA | GU902459 | GU902552 | GU902629 | GU902379 | GU902704 |

| | | | | | | |
|---|---|---|---|---|---|---|
| *Acutotyphlops subocularis* | IAA | GU902418 | GU902498 | GU902589 | GU902338 | GU902668 |
| *Afrotyphlops angolensis* | Africa | GU902389 | GU902469 | GU902562 | GU902312 | GU902639 |
| *Afrotyphlops bibronii* | Africa | GU902450 | GU902528 | GU902620 | GU902370 | GU902696 |
| *Afrotyphlops congestus* | Africa | GU902448 | GU902526 | GU902618 | GU902368 | GU902694 |
| *Afrotyphlops elegans* | Africa | GU902391 | GU902471 | GU902564 | GU902314 | GU902641 |
| *Afrotyphlops fornasinii* | Africa | GU902447 | | GU902617 | GU902367 | GU902693 |
| *Afrotyphlops lineolatus* | Africa | GU902451 | GU902529 | GU902621 | GU902371 | GU902697 |
| *Afrotyphlops mucruso* | Africa | GU902387 | GU902467 | GU902560 | GU902310 | GU902637 |
| *Afrotyphlops obtusus* | Africa | | GU902548 | GU902625 | GU902375 | GU902700 |
| *Afrotyphlops punctatus* | Africa | GU902395 | GU902475 | GU902567 | GU902318 | GU902645 |
| *Afrotyphlops schlegelii* | Africa | GU902449 | GU902527 | GU902619 | GU902369 | GU902695 |
| *Amerotyphlops brongersmianus* | America | GU902390 | GU902470 | GU902563 | GU902313 | GU902640 |
| *Amerotyphlops reticulatus* | America | GU902396 | GU902476 | GU902568 | GU902319 | GU902646 |
| *Anilios australis* | Australia | GU902409 | GU902489 | GU902580 | GU902331 | GU902659 |
| *Anilios bicolor* | Australia | GU902410 | GU902490 | GU902581 | GU902332 | GU902660 |
| *Anilios bituberculatus* | Australia | GU902403 | GU902483 | GU902574 | GU902325 | GU902653 |
| *Anilios diversus* | Australia | GU902411 | GU902491 | GU902582 | GU902333 | GU902661 |
| *Anilios endoterus* | Australia | GU902399 | GU902479 | GU902570 | | GU902649 |
| *Anilios ganei* | Australia | GU902412 | GU902492 | GU902583 | GU902334 | GU902662 |
| *Anilios grypus* | Australia | GU902413 | GU902493 | GU902584 | | GU902663 |
| *Anilios guentheri* | Australia | GU902404 | GU902484 | GU902575 | GU902326 | GU902654 |
| *Anilios hamatus* | Australia | GU902401 | GU902481 | GU902572 | GU902323 | GU902651 |
| *Anilios howi* | Australia | GU902414 | GU902494 | GU902585 | GU902335 | GU902664 |
| *Anilios kimberleyensis* | Australia | GU902406 | GU902486 | GU902577 | GU902328 | GU902656 |
| *Anilios ligatus* | Australia | GU902405 | GU902485 | GU902576 | GU902327 | GU902655 |
| *Anilios longissimus* | Australia | GU902408 | GU902488 | GU902579 | GU902330 | GU902658 |
| *Anilios pilbarensis* | Australia | GU902400 | GU902480 | GU902571 | GU902322 | GU902650 |
| *Anilios pinguis* | Australia | GU902415 | GU902495 | GU902586 | GU902336 | GU902665 |
| *Anilios splendidus* | Australia | GU902416 | GU902496 | GU902587 | GU902337 | GU902666 |
| *Anilios troglodytes* | Australia | GU902417 | GU902497 | GU902588 | | GU902667 |
| *Anilios unguirostris* | Australia | GU902407 | GU902487 | GU902578 | GU902329 | GU902657 |
| *Anilios waitii* | Australia | GU902402 | GU902482 | GU902573 | GU902324 | GU902652 |
| *Antillotyphlops catapontus* | America | GU902426 | GU902506 | GU902596 | GU902346 | GU902676 |
| *Antillotyphlops dominicanus* | America | GU902428 | GU902508 | GU902598 | GU902348 | |
| *Antillotyphlops geotomus* | America | GU902429 | GU902509 | GU902599 | GU902349 | |
| *Antillotyphlops granti* | America | GU902430 | GU902510 | GU902600 | GU902350 | GU902678 |
| *Antillotyphlops hypomethes* | America | GU902431 | GU902511 | GU902601 | GU902351 | GU902679 |
| *Antillotyphlops monastus* | America | GU902434 | GU902514 | GU902604 | GU902354 | |
| *Antillotyphlops naugus* | America | GU902435 | GU902515 | GU902605 | GU902355 | GU902681 |
| *Antillotyphlops platycephalus* | America | GU902437 | GU902517 | GU902607 | GU902357 | GU902683 |
| *Antillotyphlops richardi* | America | GU902438 | GU902518 | GU902608 | GU902358 | GU902684 |
| *Cubatyphlops anchaurus* | America | GU902423 | GU902503 | GU902593 | GU902343 | GU902673 |
| *Cubatyphlops anousius* | America | GU902445 | GU902524 | GU902615 | GU902365 | GU902691 |

| | | | | | | |
|---|---|---|---|---|---|---|
| *Cubatyphlops arator* | America | GU902424 | GU902504 | GU902594 | GU902344 | GU902674 |
| *Cubatyphlops caymanensis* | America | GU902427 | GU902507 | GU902597 | GU902347 | GU902677 |
| *Cubatyphlops contorhinus* | America | GU902446 | GU902525 | GU902616 | GU902366 | GU902692 |
| *Cubatyphlops notorachius* | America | GU902436 | GU902516 | GU902606 | GU902356 | GU902682 |
| *Gerrhopilus hedraeus* | IAA | GU902392 | GU902472 | GU902565 | GU902315 | GU902642 |
| *Gerrhopilus mirus* | India | GU902394 | GU902474 | GU902566 | GU902317 | GU902644 |
| *Indotyphlops albiceps* | IAA | GU902382 | GU902462 | GU902555 | GU902305 | GU902632 |
| *Indotyphlops braminus* | Africa Asia | GU902383 | GU902463 | GU902556 | GU902306 | GU902633 |
| *Indotyphlops pammeces* | India | GU902458 | GU902551 | GU902628 | GU902378 | GU902703 |
| *Letheobia feae* | Africa | GU902385 | GU902465 | GU902558 | GU902308 | GU902635 |
| *Letheobia newtoni* | Africa | GU902388 | GU902468 | GU902561 | GU902311 | GU902638 |
| *Madatyphlops andasibensis* | Madagascar | GU902453 | GU902545 | GU902622 | GU902373 | GU902698 |
| *Madatyphlops arenarius* | Madagascar | GU902455 | GU902547 | GU902624 | GU902374 | GU902699 |
| *Malayotyphlops luzonensis* | IAA | GU902393 | GU902473 | | GU902316 | GU902643 |
| *Ramphotyphlops acuticaudus* | IAA | GU902381 | GU902461 | GU902554 | GU902304 | GU902631 |
| *Ramphotyphlops lineatus* | IAA | GU902384 | GU902464 | GU902557 | GU902307 | GU902634 |
| *Ramphotyphlops sp* | IAA | GU902420 | GU902500 | | GU902340 | GU902670 |
| *Rhinotyphlops lalandei* | Africa | GU902386 | GU902466 | GU902559 | GU902309 | GU902636 |
| *Rhinotyphlops unitaeniatus* | Africa | GU902452 | GU902530 | | GU902372 | |
| *Sundatyphlops polygrammicus* | IAA | GU902421 | GU902501 | GU902591 | GU902341 | GU902671 |
| *Typhlophis squamosus* | America | GU902398 | GU902478 | | GU902321 | GU902648 |
| *Typhlops agoralionis* | America | GU902422 | GU902502 | GU902592 | GU902342 | GU902672 |
| *Typhlops capitulatus* | America | GU902425 | GU902505 | GU902595 | GU902345 | GU902675 |
| *Typhlops eperopeus* | America | GU902444 | GU902523 | GU902614 | GU902364 | GU902690 |
| *Typhlops jamaicensis* | America | GU902432 | GU902512 | GU902602 | GU902352 | |
| *Typhlops lumbricalis* | America | GU902433 | GU902513 | GU902603 | GU902353 | GU902680 |
| *Typhlops rostellatus* | America | GU902439 | GU902546 | GU902609 | GU902359 | GU902685 |
| *Typhlops schwartzi* | America | GU902440 | GU902519 | GU902610 | GU902360 | GU902686 |
| *Typhlops sp* | America | GU902454 | GU902553 | GU902623 | GU902380 | GU902705 |
| *Typhlops sulcatus* | America | GU902441 | GU902520 | GU902611 | GU902361 | GU902687 |
| *Typhlops sylleptor* | America | GU902442 | GU902521 | GU902612 | GU902362 | GU902688 |
| *Typhlops syntherus* | America | GU902443 | GU902522 | GU902613 | GU902363 | GU902689 |
| *Xenotyphlops grandidieri* | Madagascar | GU902457 | GU902550 | GU902627 | GU902377 | GU902702 |
| *Xenotyphlops sp* | Madagascar | GU902456 | GU902549 | GU902626 | GU902376 | GU902701 |
| *Xerotyphlops vermicularis* | Eurasia | GU902397 | GU902477 | GU902569 | GU902320 | GU902647 |

#### 16. Natricinae

Table S13: Genbank numbers for the sequences used in the Natricinae analyses with the distribution attributed to each terminal.

| Species | Locality | 12S | CMOS | COI | CytB | ND1 | ND2 | ND4 |
|---|---|---|---|---|---|---|---|---|
| *Adelophis foxi* | America | | | | AF420069 | AF420070 | AF420071 | AF420072 |
| *Afronatrix anoscopus* | Africa | | AF471123 | | AF420073 | AF420074 | AF420075 | AF420076 |

| | | | | | | | | |
|---|---|---|---|---|---|---|---|---|
| *Amphiesma craspedogaster* | Asia | | JQ687437 | | JQ687429 | | JQ687459 | JQ687412 |
| *Amphiesma sauteri* | Asia | AF402622 | | | AF402905 | | AF384824 | |
| *Amphiesma stolatum* | IAA | | JQ687450 | | JQ687432 | | JQ687464 | JQ687425 |
| *Atretium yunnanensis* | Asia | | JQ687448 | | GQ281787 | | JQ687463 | JQ687423 |
| *Clonophis kirtlandii* | America | AF402625 | | | AF402908 | | AF384827 | |
| *Coluber constrictor* | America | AY122667 | AY486937 | AY122649 | AY486913 | AY486962 | AY487001 | AY487040 |
| *Enhydris plumbea* | IAA | DQ343650 | EF395934 | DQ343650 | DQ343650 | DQ343650 | DQ343650 | DQ343650 |
| *Imantodes cenchoa* | America | EU728586 | GQ457865 | EU728586 | EU728586 | EU728586 | EU728586 | EU728586 |
| *Lycognathophis seychellensis* | MI | | FJ387220 | | | | | |
| *Macropisthodon rudis* | Asia | | JQ687442 | | GQ281780 | | JQ687458 | JQ687417 |
| *Natriciteres olivacea* | Africa | AF544772 | AF471146 | | AF471058 | | | |
| *Natrix maura* | Africa | AF402623 | | | AY866530 | AY873742 | AY870616 | EU437572 |
| *Natrix natrix* | Eurasia | AY122682 | AF471121 | AY122664 | AF471059 | AY873760 | AY870640 | AY873736 |
| *Natrix tessellata* | Arabic Peninsula | | | | EU119168 | AY873772 | AY870641 | AY873735 |
| *Nerodia cyclopion* | America | AF402626 | | GQ279084 | AF402909 | | AF384828 | |
| *Nerodia erythrogaster* | America | AF402629 | | GQ279023 | GQ285504 | HQ121595 | GQ285402 | AF420084 |
| *Nerodia fasciata* | America | AF402627 | | GQ279080 | AY866529 | AY873738 | AY870612 | AY873705 |
| *Nerodia floridana* | America | AF402628 | | | AF402911 | | AF384830 | |
| *Nerodia harteri* | America | AF402652 | | | AF402935 | | AF384854 | |
| *Nerodia rhombifer* | America | | | GQ279077 | GQ285446 | HQ121995 | GQ285289 | |
| *Nerodia sipedon* | America | AF402630 | | GQ278935 | AF402913 | HQ121602 | DQ915161 | |
| *Nerodia taxispilota* | America | AF402631 | | GQ279082 | AF402914 | HQ121606 | AF384833 | U49322 |
| *Opisthotropis cheni* | Asia | | JQ687441 | | GQ281779 | | JQ687457 | JQ687416 |
| *Opisthotropis guangxiensis* | Asia | | JQ687447 | | GQ281776 | | JQ687462 | JQ687422 |
| *Opisthotropis lateralis* | Asia | | JQ687445 | | GQ281782 | | JQ687461 | JQ687420 |
| *Opisthotropis latouchii* | Asia | | JQ687446 | | GQ281783 | | | JQ687421 |
| *Regina alleni* | America | AF402633 | | | AF402916 | | AF384835 | |
| *Regina grahami* | America | AF402635 | | | AF402918 | HQ121593 | AF384837 | |
| *Regina rigida* | America | AF402636 | AF471120 | | AF471052 | | AF384838 | |
| *Regina septemvittata* | America | AF402634 | | | AF402917 | | AF384836 | |
| *Rhabdophis nuchalis* | Asia | AF402624 | JQ687438 | | AF402907 | | AF384826 | JQ687413 |
| *Rhabdophis subminiatus* | Asia | AF544776 | AF544713 | | GQ281777 | | | U49325 |
| *Rhabdophis tigrinus* | Asia | AF236679 | AF471119 | | AF471051 | | JQ687460 | JQ687419 |
| *Seminatrix pygaea* | America | AF402637 | | | AF402920 | | AF384839 | |
| *Sinonatrix aequifasciata* | Asia | | JQ687440 | | JQ687430 | | JQ687456 | JQ687415 |
| *Sinonatrix annularis* | Asia | AF236677 | AF544712 | | JQ687431 | | | |
| *Sinonatrix percarinata 1* | Asia | | JQ687439 | | GQ281784 | | JQ687455 | JQ687414 |
| *Sinonatrix percarinata 2* | IAA | | JQ687451 | | JQ687433 | | JQ687465 | JQ687426 |
| *Storeria dekayi* | America | AF402639 | AF471154 | EF417389 | AF471050 | EF417436 | EF417460 | EF417365 |
| *Storeria occipitomaculata* | America | AF402638 | | | AF402921 | | AF384840 | U49323 |
| *Thamnophis atratus* | America | | | | AF420085 | AF420086 | AF420087 | AF420088 |
| *Thamnophis brachystoma* | America | | | | AF420089 | AF420090 | HM630351 | AF420092 |
| *Thamnophis butleri* | America | AF402640 | | | AF402923 | AF420093 | HM630350 | AF420095 |

| | | | | | | | | |
|---|---|---|---|---|---|---|---|---|
| *Thamnophis chrysocephalus* | America | | | EF460849 | AF420108 | AF420096 | AF420097 | AF420098 |
| *Thamnophis couchii* | America | AF402653 | | | AF402936 | AF420104 | AF384855 | AF420106 |
| *Thamnophis cyrtopsis* | America | AF402641 | | EF417388 | EF417412 | EF417435 | EF417459 | EF417364 |
| *Thamnophis elegans* | America | AF402642 | | | AF402925 | AF420114 | AF384844 | AF420116 |
| *Thamnophis eques* | America | | | | AF420117 | AF420118 | AF420119 | AF420120 |
| *Thamnophis exsul* | America | | | | AF420125 | AF420126 | AF420127 | AF420128 |
| *Thamnophis fulvus* | America | | | | AF420129 | AF420130 | AF420131 | AF420132 |
| *Thamnophis gigas* | America | | | | AF420133 | AF420210 | AF420209 | AF414099 |
| *Thamnophis godmani* | America | | AF471165 | | AF420135 | AF420136 | AF420137 | AF420138 |
| *Thamnophis hammondii* | America | | | | AF420139 | AF420140 | AF420141 | AF420142 |
| *Thamnophis marcianus* | America | AF402643 | | | AF402926 | AF420144 | AF384845 | AF420146 |
| *Thamnophis melanogaster* | America | | | EF417386 | EF417410 | EF417433 | EF417457 | EF417362 |
| *Thamnophis mendax* | America | | | | AF420151 | | AF420152 | |
| *Thamnophis ordinoides* | America | AF402644 | | | AF402927 | AF420158 | AF384846 | AF420160 |
| *Thamnophis proximus* | America | AF402645 | | | AF402928 | AF420162 | AF384847 | AF420164 |
| *Thamnophis radix* | America | AF402651 | | | AF402934 | AF420170 | HM630344 | AF420172 |
| *Thamnophis rufipunctatus* | America | | | | AF420173 | AF420174 | AF420175 | AF420176 |
| *Thamnophis sauritus* | America | | | | AF420177 | AF420178 | AF420179 | AF420180 |
| *Thamnophis scalaris* | America | | | | AF420181 | AF420182 | AF420183 | AF420184 |
| *Thamnophis scaliger* | America | | | | AF420189 | AF420190 | AF420191 | AF420192 |
| *Thamnophis sirtalis* | America | AF402647 | DQ902094 | | AF402929 | AF420194 | DQ995396 | AY136272 |
| *Thamnophis sumichrasti* | America | | | | AF420197 | AF420198 | AF420199 | AF420200 |
| *Thamnophis valida* | America | | | EF417385 | EF417408 | EF417432 | EF417437 | EF417360 |
| *Trachischium monticola* | India | | JQ687453 | | JQ687435 | | | JQ687428 |
| *Tropidoclonion lineatum* | America | | | | AF420205 | AF420206 | AF420207 | AF420208 |
| *Virginia striatula* | America | AF402650 | | | AF402933 | | AF384852 | |
| *Xenochrophis flavipunctatus* | IAA | AF544780 | AF544714 | | | | FJ416748 | |
| *Xenochrophis piscator* | IAA India | GQ225679 | GQ225669 | GQ225666 | GQ225659 | | | |
| *Xenochrophis punctulatus* | IAA | | AF471106 | | AF471079 | AY486996 | AY487035 | AY487074 |
| *Xenochrophis schnurrenbergeri* | India | GQ225678 | GQ225668 | GQ225663 | GQ225660 | | | |
| *Xenochrophis vittatus* | IAA | EF395871 | EF395920 | | EF395895 | | | |

In these analyses, we used the same calibration points as the original article, but we have made some alterations in the distribution parameters: for instance, in the Natricinae calibration, we set a lognormal distribution with offset of 30 m.y.a., mean of 0.4, and a standard deviation of 0.8. This yielded a 95% confidence interval from 30 to 37 m.y.a., which included the age of the fossil (30-32 m.y.a.) and set a soft constraint in the lower bound of the distribution. The same principle was applied fro the other two points: the *Natrix* calibration point (lognormal distribution; M = 0.4 and S = 0.9; offset = 22.0 m.y.a.) and the *Thamnophis* (lognormal distribution; M = 0.8 and S = 0.72; offset = 16.0 m.y.a.). We combined the logs for three independent analyses to obtain the final result.

#### 17. Testudinidae

Table S14: Genbank numbers for the sequences used in the Testudinidae analyses with the distribution attributed to each terminal.

| Species | Location | 12S | 16S | CytB | CMOS | RAG2 |
|---|---|---|---|---|---|---|
| *Chersina angulata* | Africa | DQ497248 | DQ497269 | DQ497292 | DQ497325 | DQ497361 |
| *Aldabrachelys arnoldii* | Seychelles | AY081779 | AY081780 | DQ497293 | DQ497326 | DQ497362 |
| *Aldabrachelys dussumieri* | Seychelles | DQ497249 | DQ497270 | DQ497294 | DQ497327 | DQ497363 |
| *Aldabrachelys hololissa* | Seychelles | AY081783 | AY081784 | DQ497295 | DQ497328 | DQ497364 |
| *Chelonoidis carbonarius* | America | AB090019 | AF192926 | DQ497296 | DQ497329 | DQ497365 |
| *Chelonoidis chilensis* | America | AF175336 | AF192924 | DQ497297 | DQ497330 | DQ497366 |
| *Chelonoidis denticulatus* | America | DQ497250 | AF192927 | DQ497298 | DQ497331 | DQ497367 |
| *Geochelone elegans* | India | AY081785 | AY081786 | DQ497299 | DQ497332 | DQ497368 |
| *Chelonoidis nigra* | America | AF020885 | AF020892 | DQ497300 | DQ497333 | DQ497369 |
| *Stigmochelys pardalis pardalis* | Africa | DQ497251 | DQ497271 | DQ497301 | DQ497334 | DQ497370 |
| *Stigmochelys pardalis babcocki* | Africa | DQ497252 | DQ497272 | DQ497302 | DQ497335 | DQ497371 |
| *Geochelone platynota* | IAA | DQ497253 | DQ497273 | DQ497303 | DQ497336 | DQ497372 |
| *Astrochelys radiata* | Madagascar | AF020883 | AF020890 | DQ497304 | DQ497337 | DQ497373 |
| *Centrochelys sulcata* | Africa | AY081787 | AY081788 | DQ497305 | DQ497338 | DQ497374 |
| *Astrochelys yniphora* | Madagascar | AF020882 | AF020889 | DQ497306 | DQ497339 | DQ497375 |
| *Gopherus agassizii* | America | AY434630 | | AY434562 | | |
| *Gopherus polyphemus* | America | AF020879 | AF020886 | DQ497307 | DQ497340 | DQ497376 |
| *Homopus boulengeri* | Africa | DQ497254 | DQ497274 | DQ497308 | DQ497341 | DQ497377 |
| *Homopus signatus* | Africa | DQ497255 | DQ497275 | DQ497309 | DQ497342 | DQ497378 |
| *Indotestudo elongata* | Asia IAA | DQ497256 | DQ497276 | DQ497310 | DQ497343 | DQ497379 |
| *Indotestudo forstenii* | IAA | DQ080044 | DQ080044 | DQ080044 | | |
| *Indotestudo travancorica* | India | DQ497257 | DQ497277 | DQ497311 | DQ497344 | DQ497380 |
| *Kinixys belliana* | Africa | DQ497258 | DQ497278 | DQ497312 | DQ497345 | DQ497381 |
| *Kinixys homeana* | Africa | DQ497259 | DQ497279 | DQ497313 | DQ497346 | DQ497382 |
| *Malacochersus tornieri* | Africa | DQ497260 | DQ497280 | DQ497314 | DQ497347 | DQ497383 |
| *Manouria emys emys* | IAA | DQ497261 | DQ497281 | DQ497315 | DQ497348 | DQ497384 |
| *Manouria emys phayrei* | Asia | DQ497262 | DQ497282 | DQ497316 | DQ497349 | DQ497385 |
| *Manouria impressa* | Asia IAA | DQ497263 | DQ497283 | DQ497317 | DQ497350 | DQ497386 |
| *Psammobates tentorius* | Africa | DQ497264 | DQ497284 | DQ497318 | DQ497351 | DQ497387 |
| *Pyxis arachnoides* | Madagascar | AF020880 | AF020887 | DQ497319 | DQ497352 | DQ497388 |
| *Pyxis planicauda* | Madagascar | AF020881 | AF020888 | DQ497320 | DQ497353 | DQ497389 |
| *Testudo graeca* | Africa Arabic Peninsula | AF175330 | DQ497285 | DQ497321 | DQ497354 | DQ497390 |
| *Agrionemys horsfieldii* | Asia | AB090020 | DQ497286 | DQ497322 | DQ497355 | DQ497391 |
| *Testudo kleinmanni* | Africa | AF175332 | DQ497287 | DQ497323 | DQ497356 | DQ497392 |
| *Cylindraspis indica* | MI | | | AF371247 | | |
| *Cylindraspis peltastes* | MI | | | AF371256 | | |
| *Cylindraspis vosmaeri* | MI | | | AF371260 | | |
| *Glyptemys insculpta* | America | DQ497265 | DQ497288 | AF258876 | DQ497357 | DQ497393 |

| | | | | | | |
|---|---|---|---|---|---|---|
| *Deirochelys reticularia* | America | DQ497266 | DQ497289 | AF258877 | DQ497358 | DQ497394 |
| *Rhinoclemmys melanosterna* | America | DQ497267 | DQ497290 | AY434590 | DQ497359 | DQ497395 |
| *Rhinoclemmys nasuta* | America | DQ497268 | DQ497291 | DQ497324 | DQ497360 | DQ497396 |
| *Platysternon megacephalum* | Asia | DQ256377 | DQ256377 | DQ256377 | | KC683679 |

The calibration points for this analysis were taken from (Joyce, Parham, Lyson, Warnock, & Donoghue, 2013): a calibration point was set for the outgroup based on the Emydidae-*Platysternon* (lognormal distribution; M = 3.6 and S = 0.289; offset = 32.0 m.y.a.) and another calibration point was set to the root based on the age of the Testudinoidea (lognormal distribution; M = 3.12 and S = 0.4; offset = 50.3 m.y.a.). These parameters were set in a way that the offset was the minimum age constraints used by Joyce et al. (2013) and the median of the distribution was the lower bound of the 95% confidence interval obtained by their analyses. In this way, we allowed soft constraints for both upper and lower bounds of the calibration distribution to avoid age overestimation. We made three independent runs and combined their results through LogCombiner.

#### 18. Falconidae

Table S15: Genbank numbers for the sequences used in the Falconidae analyses with the distribution attributed to each terminal

| Species | Location | FGB | MB | ND1 | PEPCK | PER2 | RAG1 | TGFB2 | VIM |
|---|---|---|---|---|---|---|---|---|---|
| *Accipiter striatus* | America | KM875747 | KM875875 | KM875999 | | | | KM876435 | |
| *Caracara cheriway* | America | KM875858 | KM875961 | KM876089 | KM876202 | | KM876413 | KM876543 | KM876660 |
| *Caracara plancus* | America | JN650312 | JN650325 | JN650427 | JF909810 | JN650357 | JN650344 | JN650374 | JN650392 |
| *Cathartes aura* | America | KX534487 | KM875884 | KM876007 | KM876121 | KM876224 | KM876316 | KM876444 | KM876576 |
| *Coragyps atratus* | America | KM875756 | KM875883 | KM876006 | KM876120 | KM876223 | KM876315 | KM876443 | KM876574 |
| *Daptrius ater* | America | JN650302 | JN650317 | JN650434 | JN650402 | JN650350 | JN650333 | JN650366 | JN650385 |
| *Falco alopex* | Africa | KM875801 | KM875978 | KM876043 | | | | KM876492 | KM876610 |
| *Falco amurensis* | Africa Asia | KM875836 | KM875942 | KM876070 | KM876180 | KM876291 | KM876372 | KM876521 | KM876637 |
| *Falco araeus* | Seychelles | KM875807 | KM875915 | KM876044 | KM876159 | KM876262 | KM876351 | KM876493 | KM876611 |
| *Falco ardosiaceus* | Africa | KM875781 | KM875969 | KM876023 | | | | KM876471 | |
| *Falco berigora* | Australia | KM875735 | KM875869 | KM875987 | KM876098 | KM876302 | KM876386 | KM876423 | KM876553 |
| *Falco biarmicus* | Africa Asia | KM875788 | KM875954 | KM876082 | KM876194 | KM876303 | KM876378 | KM876477 | KM876653 |
| *Falco cenchroides* | Australia | KM875734 | KM875868 | KM875986 | KM876099 | KM876212 | KM876387 | KM876424 | KM876555 |
| *Falco cherrug* | Asia | KM875798 | KM875911 | KM876036 | KM876154 | KM876275 | KM876403 | KM876506 | KM876606 |
| *Falco chicquera* | Africa India | KM875856 | KM875968 | KM876087 | | | | KM876541 | |
| *Falco columbarius* | America Asia Europe | KM875753 | KM875876 | KM876000 | KM876110 | KM876220 | KM876321 | KM876436 | KM876567 |
| *Falco concolor* | Africa | KM875759 | KM875887 | KM876011 | KM876125 | KM876228 | KM876395 | KM876448 | KM876581 |
| *Falco cuvierii* | Africa | KM875731 | KM875972 | KM875983 | | | | KM876420 | |
| *Falco deiroleucus* | America | KM875815 | KM875922 | KM876051 | KM876165 | KM876271 | KM876357 | KM876500 | KM876619 |
| *Falco dickinsoni* | Africa | KM875793 | KM875967 | KM876031 | | | | KM876480 | |
| *Falco eleonorae* | Africa Madagascar | KM875774 | KM875896 | KM876085 | KM876138 | KM876243 | KM876338 | KM876464 | KM876657 |
| *Falco fasciinucha* | Africa | KM875822 | KM875930 | KM876057 | KM876171 | KM876277 | KM876408 | KM876509 | KM876626 |
| *Falco femoralis* | America | KM875764 | KM875952 | KM876080 | KM876192 | KM876301 | KM876376 | KM876532 | KM876651 |

| | | | | | | | | | |
|---|---|---|---|---|---|---|---|---|---|
| *Falco hypoleucos* | Australia | KM875732 | KM875866 | KM875984 | | | | KM876421 | |
| *Falco jugger* | India | | | KM875998 | | | | | |
| *Falco longipennis* | Australia | KM875741 | KM875873 | KM875994 | KM876106 | KM876213 | KM876390 | KM876431 | KM876562 |
| *Falco mexicanus* | America | KM875752 | KM875879 | KM876002 | KM876114 | KM876219 | KM876324 | KM876495 | KM876570 |
| *Falco moluccensis* | IAA | KM875831 | KM875937 | KM876066 | | | | KM876516 | |
| *Falco naumanni* | Africa Asia | KM875839 | KM875951 | KM876079 | KM876191 | KM876299 | KM876375 | KM876531 | KM876650 |
| *Falco newtoni* | Madagascar Seychelles | KM875777 | KM875900 | KM876024 | KM876143 | KM876247 | KM876340 | KM876467 | KM876597 |
| *Falco novaeseelandiae* | New Zealand | KM875766 | KM875892 | KM876014 | KM876130 | KM876237 | KM876333 | KM876455 | KM876584 |
| *Falco pelegrinoides* | America Asia Europe | KM875768 | KM875981 | | KM876129 | KM876235 | KM876330 | KM876452 | KM876582 |
| *Falco peregrinus* | Cosmopolite | KM875751 | KM875979 | KM876019 | KM876137 | KM876242 | KM876416 | KM876445 | KM876590 |
| *Falco punctatus* | Mascarene | | KM875925 | KM876053 | KM876167 | KM876272 | KM876358 | KM876503 | KM876621 |
| *Falco rufigularis* | America | KM875804 | KM875908 | KM876033 | KM876151 | KM876256 | KM876347 | KM876490 | KM876604 |
| *Falco rupicoloides* | Africa | KM875851 | KM875924 | KM876052 | KM876195 | KM876304 | KM876379 | KM876502 | KM876654 |
| *Falco rupicolus* | Africa | KM875785 | KM875902 | KM876026 | KM876146 | KM876250 | KM876344 | KM876527 | KM876645 |
| *Falco rusticolus* | America Asia Europe | KM875827 | KM875932 | KM876058 | KM876173 | KM876279 | KM876362 | KM876510 | KM876628 |
| *Falco severus* | India IAA | KM875799 | KM875971 | KM876037 | | | | | |
| *Falco sparverius* | America | KM875749 | KM875877 | KM876004 | KM876112 | KM876215 | KM876382 | KM876540 | KM876568 |
| *Falco subbuteo* | Africa Asia | KM875845 | KM875950 | KM876078 | KM876190 | KM876297 | KM876411 | KM876529 | KM876665 |
| *Falco subniger* | Australia | KM875733 | KM875867 | KM875985 | KM876096 | KM876240 | KM876336 | KM876461 | KM876551 |
| *Falco tinnunculus* | Africa Asia IAA India | KM875865 | KM875949 | KM876077 | KM876188 | KM876296 | KM876374 | KM876528 | KM876646 |
| *Falco vespertinus* | Africa Asia Europe | KM875805 | KM875914 | KM876042 | KM876158 | KM876261 | KM876350 | KM876491 | KM876609 |
| *Falco zoniventris* | Madagascar | KM875802 | | KM876038 | | | | KM876546 | |
| *Herpetotheres cachinnans* | America | KM875770 | | KM876040 | KM876157 | KM876260 | KM876398 | KM876459 | KM876608 |
| *Ibycter americanus* | America | KM875857 | KM875960 | KM876088 | KM876201 | | KM876417 | KM876542 | KM876659 |
| *Micrastur buckleyi* | America | | | JF909874 | | | | | |
| *Micrastur gilvicollis* | America | JF899902 | JF909632 | JF909866 | JF909800 | JF909714 | JF909670 | JF909843 | JF909759 |
| *Micrastur mintoni* | America | JF899891 | JF909616 | JF909860 | JF909783 | JF909693 | JF909654 | JF909824 | JF909740 |
| *Micrastur mirandollei* | America | JF899906 | JF909631 | JF909873 | JF909798 | JF909712 | JF909669 | JF909842 | JF909758 |
| *Micrastur plumbeus* | America | KM875860 | | | KM876204 | | KM876415 | KM876545 | KM876662 |
| *Micrastur ruficollis* | America | KM875769 | | KM876015 | KM876133 | | KM876397 | KM876458 | KM876661 |
| *Micrastur semitorquatus* | America | JF899914 | JF909623 | JF909870 | JF909808 | JF909721 | JF909676 | JF909850 | JF909765 |
| *Microhierax caerulescens* | Asia IAA India | KM875833 | KM875939 | KM876068 | KM876177 | KM876283 | KM876365 | KM876518 | KM876632 |
| *Microhierax erythrogenys* | IAA | KM875784 | KM875906 | KM876025 | KM876144 | KM876248 | KM876343 | KM876473 | KM876598 |
| *Microhierax fringillarius* | IAA | KM875792 | KM875975 | KM876064 | | | | KM876515 | |
| *Microhierax latifrons* | IAA | KM875791 | KM875977 | KM876030 | | | | KM876479 | |
| *Microhierax melanoleucos* | Asia India | KM875780 | KM875976 | KM876022 | | | | KM876470 | |
| *Milvago chimachima* | America | JN650309 | JN650326 | JN650430 | JN650411 | JN650358 | JN650342 | JN650375 | JN650393 |
| *Milvago chimango* | America | | | | | | | | KM876636 |
| *Phalcoboenus albogularis* | America | JN650316 | KM875935 | JN650423 | | | | | JN650401 |
| *Phalcoboenus australis* | America | | JN650319 | JN650435 | JN650403 | JN650351 | JN650334 | JN650367 | JN650386 |
| *Phalcoboenus carunculatus* | America | KM875758 | KM875886 | KM876010 | KM876124 | | KM876394 | KM876447 | KM876577 |
| *Phalcoboenus megalopterus* | America | JF899903 | JF909628 | JN650422 | JN650417 | JN650362 | JN650347 | JN650380 | JN650398 |
| *Polihierax insignis* | IAA | KM875829 | KM875936 | KM876063 | | | | KM876514 | |

| | | | | | | | | | |
|---|---|---|---|---|---|---|---|---|---|
| *Polihierax semitorquatus* | Africa | KM875834 | KM875940 | KM876069 | KM876178 | KM876284 | KM876366 | KM876519 | KM876633 |
| *Spiziapteryx circumcincta* | America | JN650306 | JN650323 | JN650432 | JN650408 | JN650355 | JN650339 | JN650371 | JN650390 |

#### 19. *Aphananthe*

Table S16: Genbank numbers for the sequences used in the *Aphananthe* analyses with the distribution attributed to each terminal.

| Species | Locality | psbA-trnH | trnL-trnF | ETS | ITS |
|---|---|---|---|---|---|
| *Aphananthe cuspidata* | Asia | KR086759 | KR086777 | KR086725 | KR086741 |
| *A. cuspidata* | IAA | KR086760 | KR086778 | KR086726 | KR086742 |
| *A. monoica* | America | KR086763 | KR086779 | KR086729 | KR086745 |
| *A. monoica* | America | KR086764 | KR086780 | KR086730 | KR086746 |
| *A. philippinensis* | Australia IAA | KR086765 | KR086781 | KR086731 | KR086747 |
| *A. philippinensis* | Australia IAA | KR086766 | JN040357 | KR086732 | KR086748 |
| *A. philippinensis* | Australia IAA | KR086767 | KR086782 | KR086733 | KR086749 |
| *A. philippinensis* | Australia IAA | KR086768 | KR086783 - | | KR086750 |
| *A. sakalava* | Madagascar | KR086770 | KR086784 - | | KR086752 |
| *A. aspera* | Asia | KR086753 | KR086771 | KR086719 | KR086735 |
| *A. aspera* | Asia | KR086754 | KR086772 | KR086720 | KR086736 |
| *A. aspera* | Asia | KR086755 | KR086773 | KR086721 | KR086737 |
| *A. aspera* | Asia | KR086756 | KR086774 | KR086722 | KR086738 |
| *Celtis bungeana* | Asia | KR086757 | KR086775 | KR086723 | KR086739 |
| *C. julianae* | Asia | KR086758 | KR086776 | KR086724 | KR086740 |
| *Gironniera subequalis* | Asia | KR086761 | JN040375 | KR086727 | KR086743 |
| *Lozanella permollis* | America | KR086762 | JN040379 | KR086728 | KR086744 |
| *Pteroceltis tatarinowii* | Asia | KR086769 | JN040385 | KR086734 | KR086751 |

#### *20. Exacum*

Table S17: Genbank numbers for the sequences used in the *Exacum* analyses with the distribution attributed to each terminal.

| Species | Locality | trnL | ITS |
|---|---|---|---|
| *Exacum affine* | Arabic Peninsula | AJ490202 | AJ489877 |
| *Exacum affine* | Arabic Peninsula | AJ490203 | AJ489878 |
| *Exacum affine* | Arabic Peninsula | AJ490204 | AJ489879 |
| *Exacum atropurpureum* | India | AJ490205 | AJ489880 |
| *Exacum bulbilliferum* | Madagascar | AJ490206 | AJ489881 |
| *Exacum caeruleum* | Arabic Peninsula | AJ490207 | AJ489882 |
| *Exacum dolichantherum* | Madagascar | AJ490208 | AJ489883 |
| *Exacum exiguum* | Madagascar | AJ490209 | AJ489884 |
| *Exacum fruticosum* | Madagascar | AJ490210 | AJ489885 |
| *Exacum gracilipes* | Arabic Peninsula | AJ490211 | AJ489886 |
| *Exacum hamiltonii* | India | AJ490212 | AJ489887 |

| | | | |
|---|---|---|---|
| *Exacum humbertii* | Madagascar | AJ490213 | AJ489888 |
| *Exacum intermedium* | Madagascar | AJ490214 | AJ489889 |
| *Exacum linearifolium* | Madagascar | AJ490215 | AJ489890 |
| *Exacum macranthum* | India | AJ490216 | AJ489891 |
| *Exacum macranthum* | India | AJ490217 | AJ489892 |
| *Exacum marojejyense* | Madagascar | AJ490218 | AJ489893 |
| *Exacum microcarpum* | Madagascar | AJ490219 | AJ489894 |
| *Exacum millotii* | Madagascar | AJ490220 | AJ489895 |
| *Exacum nummularifolium* | Madagascar | AJ490221 | AJ489896 |
| *Exacum oldenlandioides* | Africa | AJ490222 | AJ489897 |
| *Exacum pallidum* | India | AJ490223 | AJ489898 |
| *Exacum pedunculatum* | India | AJ490224 | AJ489899 |
| *Exacum quinquenervium* | Madagascar | AJ490225 | AJ489900 |
| *Exacum sessile* | India | AJ490226 | AJ489901 |
| *Exacum stenophyllum* | Madagascar | AJ490227 | AJ489902 |
| *Exacum subacaule* | Madagascar | AJ490228 | AJ489903 |
| *Exacum subteres* | Madagascar | AJ490229 | AJ489904 |
| *Exacum subverticillatum* | Madagascar | AJ490230 | AJ489905 |
| *Exacum sutaepense* | IAA | AJ490231 | AJ489906 |
| *Exacum tetragonum* | India | AJ490232 | AJ489907 |
| *Exacum tetragonum* | India | AJ490233 | AJ489908 |
| *Exacum trinervium* | India | AJ490234 | AJ489909 |
| *Exacum trinervium var. ritigalense* | India | AJ490235 | AJ489910 |
| *Exacum walkeri* | India | AJ490236 | AJ489911 |
| *Exacum walkeri* | India | AJ490237 | AJ489912 |
| *Exacum wightianum* | India | AJ490238 | AJ489913 |
| *Gentianothamnus madagascariensis* | Madagascar | AJ490240 | AJ489914 |
| *Tachiadenus carinatus* | Madagascar | AJ490249 | AJ489923 |
| *Tachiadenus longiflorus* | Madagascar | AJ490250 | AJ489924 |
| *Ornichia madagascariensis* | Madagascar | AJ490242 | AJ489917 |
| *Ornichia trinervis* | Madagascar | AJ490243 | AJ489918 |

#### 21. *Ficus*

Table S18: Genbank numbers for the sequences used in the *Ficus* analyses with the distribution attributed to each terminal.

| Species | Locality | ITS | ETS | G3pdh | ncpGS |
|---|---|---|---|---|---|
| Ficus alongensis 1 | Asia | KJ845962 | KJ845902 | KJ846015 | |
| Ficus alongensis 2 | Asia | KJ845963 | KJ845903 | | |
| Ficus arnottiana 1 | India | | KJ845879 | | |
| Ficus arnottiana 2 | India | | KJ845880 | | |
| Ficus caulocarpa 1 | IAA | KJ845953 | | | |
| Ficus caulocarpa 2 | IAA | KJ845954 | KJ845894 | KJ846009 | |

| | | | | | |
|---|---|---|---|---|---|
| Ficus caulocarpa 3 | IAA | KJ845955 | KJ845895 | KJ846010 | |
| Ficus concinna 1 | IAA | KJ845989 | KJ845928 | KJ846035 | |
| Ficus concinna 2 | IAA | KJ845990 | KJ845929 | KJ846036 | |
| Ficus concinna 3 | IAA | KJ845991 | KJ845930 | KJ846037 | KJ846071 |
| Ficus concinna 4 | IAA | KJ845992 | KJ845931 | KJ846038 | KJ846072 |
| Ficus cordata 1 | Africa | KJ845973 | KJ845912 | KJ846020 | |
| Ficus cordata 2 | Africa | KJ845974 | KJ845913 | KJ846021 | |
| Ficus cordata 3 | Africa | KJ845975 | KJ845914 | KJ846022 | KJ846063 |
| Ficus densifolia 1 | Mascarene | KJ845983 | KJ845922 | KJ846030 | KJ846068 |
| Ficus densifolia 2 | Mascarene | KJ845984 | KJ845923 | KJ846031 | KJ846069 |
| Ficus densifolia 3 | Mascarene | KJ845985 | KJ845924 | KJ846032 | |
| Ficus densifolia 4 | Mascarene | KJ845986 | KJ845925 | KJ846033 | KJ846070 |
| Ficus geniculata geniculata 1 | IAA | KJ845940 | KJ845882 | KJ845999 | KJ846044 |
| Ficus geniculata geniculata 2 | IAA | KJ845941 | KJ845883 | KJ846000 | KJ846045 |
| Ficus geniculata geniculata 3 | IAA | KJ845942 | KJ845884 | | KJ846046 |
| Ficus geniculata insignis | Australia | KJ845943 | KJ845885 | KJ846001 | KJ846047 |
| Ficus glaberrima siamensis 1 | IAA | KJ845996 | KJ845935 | KJ846041 | KJ846076 |
| Ficus glaberrima siamensis 2 | IAA | KJ845997 | KJ845936 | KJ846042 | KJ846077 |
| Ficus glaberrima siamensis 3 | IAA | KJ845998 | KJ845937 | KJ846043 | |
| Ficus henneana 1 | Australia | KJ845967 | | KJ846016 | KJ846058 |
| Ficus henneana 2 | Australia | KJ845968 | KJ845907 | | KJ846059 |
| Ficus hookeriana hookeriana | India | KJ845988 | KJ845927 | | |
| Ficus ingens 1 | Africa | KJ845964 | KJ845904 | | KJ846056 |
| Ficus ingens 2 | Africa | KJ845965 | KJ845905 | | |
| Ficus ingens 3 | Africa | KJ845966 | KJ845906 | | KJ846057 |
| Ficus lecardii1 | Africa | KJ845971 | KJ845910 | KJ846018 | KJ846061 |
| Ficus lecardii2 | Africa | KJ845972 | KJ845911 | KJ846019 | KJ846062 |
| Ficus madagascariensis | Madagascar | KJ845956 | KJ845896 | | KJ846053 |
| Ficus middletonii | IAA | KJ845952 | KJ845893 | KJ846008 | KJ846052 |
| Ficus orthoneura 1 | IAA | KJ845987 | KJ845926 | KJ846034 | |
| Ficus prasinicarpa 1 | IAA | KJ845947 | | | |
| Ficus prasinicarpa 2 | Australia | KJ845948 | KJ845889 | | |
| Ficus prolixa 1 | Pacific | KJ845949 | KJ845890 | KJ846005 | KJ846051 |
| Ficus prolixa 2 | Pacific | KJ845950 | KJ845891 | KJ846006 | |
| Ficus pseudoconcinna | IAA | KJ845946 | KJ845888 | KJ846004 | KJ846050 |
| Ficus religiosa 1 | Asia / India | KJ845980 | KJ845919 | KJ846027 | KJ846066 |
| Ficus religiosa 2 | IAA | KJ845981 | KJ845920 | KJ846028 | |
| Ficus religiosa 3 | IAA | KJ845982 | KJ845921 | KJ846029 | KJ846067 |
| Ficus cf rumphii | IAA | KJ845995 | KJ845934 | | KJ846075 |
| Ficus rumphii 1 | IAA | KJ845993 | KJ845932 | KJ846039 | KJ846073 |
| Ficus rumphii 2 | IAA | KJ845994 | KJ845933 | KJ846040 | KJ846074 |
| Ficus salicifolia 1 | Africa | KJ845976 | KJ845915 | KJ846023 | KJ846064 |
| Ficus salicifolia 2 | Arabic Peninsula | KJ845977 | KJ845916 | KJ846024 | |

| | | | | | |
|---|---|---|---|---|---|
| Ficus subpisocarpa pubipoda 1 | IAA | KJ845969 | KJ845908 | | |
| Ficus subpisocarpa pubipoda 2 | IAA | KJ845970 | KJ845909 | KJ846017 | KJ846060 |
| Ficus superba 1 | IAA | KJ845944 | KJ845886 | KJ846002 | KJ846048 |
| Ficus superba 2 | IAA | KJ845945 | KJ845887 | KJ846003 | KJ846049 |
| Ficus tsjakela | India | KJ845951 | KJ845892 | KJ846007 | |
| Ficus verruculosa 1 | Africa | KJ845978 | KJ845917 | KJ846025 | |
| Ficus verruculosa 2 | Africa | KJ845979 | KJ845917 | KJ846026 | KJ846065 |
| Ficus virens glabella 1 | IAA | KJ845960 | KJ845900 | KJ846013 | KJ846055 |
| Ficus virens glabella 2 | IAA | KJ845961 | KJ845901 | KJ846014 | |
| Ficus virens virens 1 | Australia | KJ845957 | KJ845897 | KJ846011 | KJ846054 |
| Ficus virens virens 2 | IAA | KJ845958 | KJ845898 | KJ846012 | |
| Ficus virens virens 3 | Australia | KJ845959 | KJ845899 | | |
| Ficus virens virens 4 | India | KJ845938 | KJ845881 | | |
| Ficus virens virens 5 | India | KJ845939 | | | |

#### 22. Loranthaceae

Table S19: Genbank numbers for the sequences used in the Loranthaceae analyses with the distribution attributed to each terminal

| Species | Location | LSU | SSU | matK | rbcL | trnL-F |
|---|---|---|---|---|---|---|
| *Actinanthella menyharthii* | Africa | EU544352 | EU544313 | EU544408 | | |
| *Aetanthus nodosus* | America | | EU544314 | EU544409 | | |
| *Agelanthus sansibarensis* | Africa | EU544353 | U59946 | EU544410 | EU544464 | DQ340573 |
| *Agelanthus zizyphifolius* | Africa | MG999385 | MG999460 | MG999409 | MG999430 | MG999482 |
| *Alepis flavida* | New Zealand | EF464474 | L24139 | EF464508 | KT626664 | EF464481 |
| *Amyema queenslandica* | Australia | EU544355 | EU544315 | EU544412 | | |
| *Amyema glabra* | Australia | EU544354 | AF039073 | EU544411 | EU544465 | EU544476 |
| *Amylotheca duthieana* | IAA | EU544356 | EU544316 | EU544413 | KF496311 | EU544477 |
| *Anacolosa papuana* | Australia | | DQ790104 | DQ790181 | DQ790144 | |
| *Antidaphne viscoidea* | America | | L24080 | EF464500 | L26068 | |
| *Arjona tuberosa* | America | EF464480 | EF464468 | EF464513 | EF464532 | EF464483 |
| *Atkinsonia ligustrina* | Australia | EF464475 | EF464464 | DQ787444 | EF464526 | DQ788714 |
| *Bakerella sp.* | Madagascar | EU544358 | EU544318 | EU544415 | EU544466 | EU544479 |
| *Baratranthus axanthus* | IAA | EU544357 | EU544317 | EU544414 | | EU544478 |
| *Benthamina alyxifolia* | Australia | EU544359 | EU544319 | EU544416 | AY957440 | EU544480 |
| *Berhautia senegalensis* | Africa | EU544360 | EU544320 | EU544417 | | |
| *Cathedra acuminata* | America | | FJ848847 | DQ790182 | DQ790145 | |
| *Cecarria obtusifolia* | Australia | EU544361 | EU544321 | EU544418 | EU544467 | EU544481 |
| *Cladocolea gracilis* | America | EU544362 | EU544322 | EU544419 | | EU544482 |
| *Dactyliophora novaeguineae* | Australia | EU544363 | EU544323 | EU544420 | | EU544483 |
| *Decaisnina triflora* | IAA | EU544364 | EU544324 | EU544421 | EU544468 | EU544484 |
| *Decaisnina aherniana* | IAA | MG999386 | MG999461 | MG999410 | | |
| *Dendropemon bicolor* | America | EU544365 | AF039075 | EU544422 | EU544469 | |

| | | | | | | |
|---|---|---|---|---|---|---|
| *Dendrophthoe curvata* | IAA | EU544367 | EU544325 | EU544424 | | |
| *Dendrophthoe longituba* | IAA | EU544366 | | EU544423 | | EU544485 |
| *Dendrophthoe gangliiformis* | IAA | | MG999462 | MG999411 | MG999431 | MG999483 |
| *Dendrophthoe pentandra* | Asia | MG999387 | MG999463 | MG999412 | MG999432 | MG999484 |
| *Desmaria mutabilis* | America | EF464476 | EF464465 | EF464509 | EF464527 | EF464486 |
| *Diplatia furcata* | Australia | EU544368 | L24088 | EU544425 | KF496292 | EU544486 |
| *Elytranthe albida* | Asia | MG999388 | MG999464 | MG999413 | MG999433 | MG999485 |
| *Emelianthe panganensis* | Africa | EU544369 | EU544326 | EU544426 | MG999434 | MG999486 |
| *Englerina ramulosa* | Africa | EU544370 | L24140 | EU544427 | EU544470 | DQ340577 |
| *Englerina woodfordioides* | Africa | MG999389 | MG999465 | MG999414 | MG999435 | MG999487 |
| *Erianthemum dregei* | Africa | EU544371 | L25679 | EU544428 | | EU544488 |
| *Eubrachion ambiguum* | America | AF389273 | L24141 | EF464498 | | |
| *Gaiadendron punctatum* | America | DQ790209 | L24143 | DQ787445 | L26072 | DQ340617 |
| *Globimetula dinklagei* | Africa | EU544372 | AF039076 | EU544429 | | EU544489 |
| *Helicanthes elastica* | India | EU544375 | EU544328 | EU544432 | | |
| *Helixanthera parasitica* | Asia | MG999391 | MG999466 | | MG999437 | MG999488 |
| *Helixanthera cylindrica* | IAA | EU544374 | EU544327 | EU544431 | | |
| *Helixanthera sampsonii* | Asia | | | KP093921 | KP094864 | |
| *Helixanthera coccinea* | IAA | EU544373 | | EU544430 | | EU544490 |
| *Helixanthera kirkii* | Africa | MG999390 | AF039077 | | MG999436 | |
| *Ileostylus micranthus* | New Zealand | EU544376 | EU544329 | EU544433 | EU544471 | EU544491 |
| *Lepeostegeres lanceifolius* | IAA | EU544379 | | EU544435 | | |
| *Lepidaria forbesii* | IAA | EU544378 | EU544330 | EU544434 | | EU544492 |
| *Lepidoceras chilense* | America | | EF464459 | EF464499 | EF464519 | |
| *Ligaria cuneifolia* | America | EF464477 | L24152 | EF464510 | EF464528 | DQ442940 |
| *Loranthus europaeus* | Europe | EU544380 | L24153 | EU544436 | JQ933393 | EU544493 |
| *Loranthus odoratus* | Asia | EU544381 | EU544331 | | | EU544494 |
| *Loxanthera speciosa* | IAA | EU544382 | EU544332 | EU544437 | | EU544495 |
| *Lysiana filifolia* | Australia | EU544383 | EU544333 | EU544438 | | EU544496 |
| *Macrosolen cochinchinensis* | Asia IAA | EU544384 | EU544334 | EU544439 | MG999439 | MG999490 |
| *Macrosolen tricolor* | IAA | MG999393 | MG999468 | MG999416 | MG999440 | MG999491 |
| *Macrosolen bibracteolatus* | Asia | MG999392 | MG999467 | MG999415 | MG999438 | MG999489 |
| *Misodendrum linearifolium* | America | DQ790211 | L24397 | DQ787438 | L26074 | DQ788712 |
| *Misodendrum punctulatum* | America | KP263250 | KP263284 | DQ787443 | EF464531 | DQ788711 |
| *Moquiniella rubra* | Africa | DQ790207 | AF039078 | DQ790171 | DQ790132 | EF464489 |
| *Muellerina eucalyptoides* | Australia | EU544385 | EU544335 | EU544440 | EU544472 | EU544498 |
| *Notanthera heterophylla* | America | EF464478 | EF464466 | EF464511 | EF464529 | DQ442939 |
| *Nuytsia floribunda* | Australia | DQ790210 | DQ790103 | DQ787446 | DQ790134 | DQ788716 |
| *Oedina pendens* | Africa | EU544386 | EU544336 | EU544441 | | EU544499 |
| *Oliverella rubroviridis* | Africa | EU544387 | EU544337 | EU544442 | | |
| *Oncella ambigua* | Africa | | EU544338 | EU544443 | | |
| *Oncocalyx sulfureus* | Africa | EU544388 | EU544339 | EU544444 | MG999442 | EU544500 |
| *Oncocalyx fischeri* | Africa | MG999394 | MG999469 | MG999417 | MG999441 | MG999492 |

| | | | | | | |
|---|---|---|---|---|---|---|
| *Oryctanthus occidentalis* | America | EU544389 | L24408 | EU544445 | | EU544501 |
| *Osyris alba* | Europe | MG999395 | | | MG999443 | MG999493 |
| *Osyris lanceolata* | Africa | MG999396 | MG999470 | MG999418 | MG999444 | MG999494 |
| *Passovia pyrifolia* | America | EU544392 | L24412 | EU544448 | | EU544504 |
| *Peraxilla tetrapetala* | New Zealand | EU544390 | EU544340 | EU544446 | JQ933439 | EU544502 |
| *Phanerodiscus capuronii* | Madagascar | DQ790219 | DQ790122 | DQ790180 | DQ790143 | |
| *Phragmanthera crassicaulis* | Africa | EU544391 | EU544341 | EU544447 | | EU544503 |
| *Phragmanthera regularis* | Africa | MG999397 | MG999471 | MG999419 | MG999445 | DQ340579 |
| *Plicosepalus sagittifolius* | Africa | EU544393 | EU544342 | EU544449 | MG999447 | MG999496 |
| *Plicosepalus curviflorus* | Africa | MG999398 | MG999472 | MG999420 | MG999446 | MG999495 |
| *Psittacanthus calyculatus* | America | EU544394 | L24414 | EU544450 | | DQ340610 |
| *Quinchamalium chilense* | America | KP263257 | EF464469 | EF464514 | EF464533 | EF464491 |
| *Santalum album* | Asia | MG999399 | MG999473 | MG999421 | MG999448 | MG999497 |
| *Schoepfia fragrans* | Asia | MG999400 | MG999474 | MG999422 | MG999449 | MG999498 |
| *Schoepfia jasminodora* | Asia | MG999401 | JQ613226 | HQ415321 | MG999450 | AY191152 |
| *Schoepfia schreberi* | America | AF389261 | L24418 | AY957454 | L11205 | DQ788717 |
| *Scurrula ferruginea* | IAA | EU544395 | EU544343 | EU544451 | KF114863 | EU544505 |
| *Scurrula parasitica* | Asia - IAA | EU544397 | EU544345 | EU544451 | MG999454 | MG999502 |
| *Scurrula pulverulenta* | Asia | EU544396 | EU544344 | EU544452 | | |
| *Scurrula chingii* | Asia | MG999403 | MG999476 | MG999424 | MG999452 | MG999500 |
| *Scurrula chingii* | Asia | MG999404 | MG999477 | MG999425 | MG999453 | MG999501 |
| *Scurrula philippensis* | Asia | MG999405 | MG999478 | MG999426 | MG999455 | MG999503 |
| *Scurrula buddleioides* | Asia | MG999402 | MG999475 | MG999423 | MG999451 | MG999499 |
| *Septulina glauca* | Africa | EU544398 | EU544346 | | AM235022 | EU544506 |
| *Socratina bemarivensis* | Madagascar | EU544399 | EU544347 | EU544454 | | EU544507 |
| *Sogerianthe sesailiflora* | IAA | EU544400 | EU544348 | EU544455 | | EU544508 |
| *Spragueanella rhamnifolia* | Africa | | | EU544456 | | |
| *Struthanthus oerstedii* | America | EU544402 | L24421 | EU544457 | JQ594622 | EU544509 |
| *Struthanthus woodsonii* | America | EU544403 | EU544349 | EU544458 | EU544474 | EU544510 |
| *Tapinanthus constrictiflorus* | Africa | EU544404 | L24422 | EU544459 | EU213529 | EU544511 |
| *Taxillus chinensis* | IAA | EU544405 | EU544350 | EU544460 | MG999456 | MG999504 |
| *Taxillus thibetensis* | Asia | MG999407 | MG999480 | MG999428 | MG999458 | MG999506 |
| *Taxillus tsaii* | Asia | MG999408 | MG999481 | MG999429 | MG999459 | MG999507 |
| *Taxillus sutchuenensis* | Asia | MG999406 | MG999479 | MG999427 | MG999457 | MG999505 |
| *Tolypanthus involucratus* | Asia | | | EU544461 | | |
| *Tripodanthus acutifolius* | America | EU544406 | L24424 | EU544462 | EU544475 | EU544513 |
| *Tristerix corymbosus* | America | EF464479 | EF464467 | EF464512 | EF464530 | DQ340605 |
| *Tupeia antarctica* | New Zealand | DQ790208 | L24425 | DQ790172 | DQ790133 | EF464494 |
| *Vanwykia remota* | Africa | EU544407 | EU544351 | EU544463 | | EU544514 |

#### 23. Rubiaceae

Sample list was taken from the supplementary material of the original article (available at: https://onlinelibrary.wiley.com/action/downloadSupplement?doi=10.1111%2Fjbi.12981&file=jbi12981-sup-0001-AppendixS1.xls). In order to simplify the analysis, we chose the tribes that presented a diverse biogeographic distribution (Coffeae, Bertiereae, Octotropideae, and Pavetteae) and built the trees independently. In the alignments, we excluded the large gaps that represented apomorphic indels (of a single terminal). The calibration was made by extrapolating the dates obtained by the original article. For the Coffeae-Bertiereae analysis three calibration points were set: Bertiereae (lognormal distribution; M = 0.5 and S = 0.87; offset = 10.0 m.y.a.), Coffeae (lognormal distribution; M = 0.5 and S = 0.87; offset = 11.0 m.y.a.), and a root calibration (lognormal distribution; M = 0.5 and S = 0.95; offset = 15.0 m.y.a.). For the Octotropideae analysis three calibration points were set: root (lognormal distribution; M = 1.65 and S = 0.345; offset = 16.7 m.y.a.), *Didymosalpinx-Mantalania* (lognormal distribution; M = 1.74 and S = 0.33; offset = 14.0 m.y.a.), and *Burchellia-Polysphaeria* (lognormal distribution; M = 1.64 and S = 0.344; offset = 15.7 m.y.a.). For the Pavetteae-Sherbournieae analysis three calibration points were set: root (lognormal distribution; M = 1.48 and S = 0.34; offset = 16.4 m.y.a.), Pavetteae (lognormal distribution; M = 1.22 and S = 0.352; offset = 10.8 m.y.a.), and Sherbournieae (lognormal distribution; M = 1.73 and S = 0.35; offset = 9.97 m.y.a.).
