## Supplementary material S0 to S4 for "Ecological changes have driven biotic exchanges across the Indian Ocean": S3.pdf

### Appendix 3: Coordinates used in ancestral range reconstruction

Table S20: Geographic coordinates for the regions as used in BayArea analyses.

| Region | Latitude | Longitude |
|---|---|---|
| Africa | -1.2500 | 25.7225 |
| Asia | 39.7054 | 81.8409 |
| Australia | -24.1385 | 133.3899 |
| Comoros | -12.1837 | 44.0731 |
| IAA | 1.2500 | 113.4847 |
| India | 20.6835 | 79.1195 |
| Madagascar | -19.4258 | 46.6878 |
| Mascarene | -20.2856 | 57.5659 |
| Seychelles | -4.7005 | 55.4899 |
| America | 10.7769 | -79.1874 |
| Pacific | -13.9791 | -173.2304 |
| New Zealand | -43.6889 | 171.00271 |
| Europe | 46.309 | 20.4464 |
| Arabic Peninsula | 22.2594 | 47.8352 |
| Mediterranean | 36.637 | -2.9501 |
